# Esr1-Dependent Signaling and Transcriptional Maturation in the Medial Preoptic Area of the Hypothalamus Shapes the Development of Mating Behavior during Adolescence

**DOI:** 10.1101/2025.02.26.640339

**Authors:** Koichi Hashikawa, Yoshiko Hashikawa, Brandy Briones, Kentaro Ishii, Yuejia Liu, Mark A. Rossi, Marcus L. Basiri, Jane Y. Chen, Omar R. Ahmad, Rishi V. Mukundan, Nathan L. Johnston, Rhiana C. Simon, James C. Soetedjo, Jason R. Siputro, Jenna A. McHenry, Richard D. Palmiter, David R. Rubinow, Larry S. Zweifel, Garret D. Stuber

## Abstract

Mating and other behaviors emerge during adolescence through the coordinated actions of steroid hormone signaling throughout the nervous system and periphery. In this study, we investigated the transcriptional dynamics of the medial preoptic area (MPOA), a critical region for reproductive behavior, using single-cell RNA sequencing (scRNAseq) and *in situ* hybridization techniques in male and female mice throughout adolescence development. Our findings reveal that estrogen receptor 1 (Esr1) plays a pivotal role in the transcriptional maturation of GABAergic neurons within the MPOA during adolescence. Deletion of the estrogen receptor gene, *Esr1*, in GABAergic neurons (Vgat+) disrupted the developmental progression of mating behaviors in both sexes, while its deletion in glutamatergic neurons (Vglut2+) had no observable effect. In males and females, these neurons displayed distinct transcriptional trajectories, with hormone-dependent gene expression patterns emerging throughout adolescence and regulated by *Esr1*. *Esr1* deletion in MPOA GABAergic neurons, prior to adolescence, arrested adolescent transcriptional progression of these cells and uncovered sex-specific gene-regulatory networks associated with *Esr1* signaling. Our results underscore the critical role of *Esr1* in orchestrating sex-specific transcriptional dynamics during adolescence, revealing gene regulatory networks implicated in the development of hypothalamic controlled reproductive behaviors.

**One Sentence Summary:** Single cell RNA sequencing reveals how adolescent sex hormones sculpt hypothalamic cell types required for mating behavior.

---

Adolescence is a secondary critical period, following the perinatal steroid surge, during which juvenile animals undergo extensive physiological maturation of both the body and nervous system as they transition into adulthood ^1,2^. This maturation is primarily orchestrated by the hypothalamus-pituitary-gonad (HPG) axis, which activates gonadal cells in the testes and ovaries to release testosterone and estrogen, initiated during adolescence ^1,3,4^. These circulating sex steroids are essential not only for reproductive maturation but also for guiding the development and execution of sex-specific behaviors through their direct actions on the central nervous system ^5–7^. Despite the well-established role of these hormones in shaping behavior, the molecular mechanisms underlying their influence on brain development during adolescence are still limited to brain-region level (bulk)^8^ in humans and model organisms and adolescent transcriptional dynamics at single cell resolution in the brain remain poorly understood (but see a pioneering study in the human testis^9^).

The medial preoptic area (MPOA) is a critical region in the brain, known for its involvement in mating and other social behaviors ^7,10–16^. Within the MPOA, specific molecularly defined subpopulations of neurons express various neuropeptides and hormone receptors that are closely tied to reproductive behavior ^16–20^. Notably, the MPOA is enriched with the expression of steroid hormone receptor genes such as *Esr1,* or estrogen receptor 1 ^21,22^, leading us to hypothesize that gonadal steroid signaling during adolescence facilitates the transcriptional maturation of these genetically defined MPOA neurons, which are crucial for the development of mating behaviors. While recent advances have shed light on the molecular diversity of neurons in sexually dimorphic brain regions during the perinatal period ^23^ and adulthood ^22,24,25^, the transcriptional dynamics that occur during adolescence, particularly at the single-cell level, remain largely unexplored. This gap in our understanding represents a critical frontier in unraveling how hormonal signaling shapes the neural circuits that govern sex-specific behaviors.

In this study, we used single-cell RNA sequencing (scRNAseq) and hybridized chain reaction fluorescent *in situ* hybridization (HCR-FISH) to investigate the transcriptional dynamics of the MPOA throughout adolescence in both male and female mice. We show that *Esr1* plays a critical role in the transcriptional maturation of GABAergic neurons (Vgat+) within the MPOA, which is essential for the development of mating behaviors. Deleting *Esr1* in Vgat+ neurons disrupted the development of mating behaviors in both sexes, while deletion in glutamatergic neurons (Vglut2+) had no observable effect. Our findings highlight the importance of *Esr1* in guiding sex-specific transcriptional programs during adolescence, revealing distinct hormone-dependent gene expression patterns and gene-regulatory networks crucial for reproductive behaviors.

## Results

### Selective knockout of *Esr1* in GABAergic or glutamatergic MPOA neurons at pre-adolescence

Previous studies have shown that knocking out *Esr1*, either globally or in specific cell types, reduces mating behavior ^7,26^. However, the role of *Esr1* in genetically defined MPOA neurons during adolescent development remains unclear, as these studies could not easily differentiate between effects occurring during perinatal and adolescent periods. To address this gap and directly test whether the emergence of mating behavior requires steroid hormone signaling in molecularly defined MPOA cell types, we developed an AAV-frtFlex-Cre virus. We injected this virus into the MPOA of *Slc32a1*^Flp^::*Esr1*^lox/lox^ (MPOA^Vgat-Esr1KO^) or *Slc17a6*^Flp^::*Esr1*^lox/lox^ (MPOA^Vglut2-Esr1KO^) male and female mice at postnatal day (P) 14-18. This viral strategy allowed for *Esr1* deletion in MPOA GABAergic or glutamatergic neurons, depending on the transgenic mouse line, throughout adolescence (**Fig. 1a-b,i-j and Supplementary Fig. 1a-b**). The onset of genital changes, the maturation of the HPG axis (as determined by the first day of estrus in female mice^27^ and first day of initiation of sexual behaviors in male mice^28^), and locomotor behaviors were not altered in MPOA^Vgat-Esr1KO^ and MPOA^Vglut2-Esr1KO^ mice (**Fig. 1c-d,f,k-l,n and Supplementary Fig. 1e-f**). The establishment of mating behavior was abolished in MPOA^Vgat-Esr1KO^ mice in both sexes (**Fig. 1d-e,g-h and Supplementary Fig. 1a-b,g**), but unaffected in MPOA^Vglut2-Esr1KO^ mice (**Fig. 1l-m,o-p and Supplementary Fig. 1h**). These results extend prior work to show that *Esr1* signaling in MPOA GABAergic neurons is required for the emergence of mating behavior during adolescence ^7,14,15^.

**Fig. 1:**
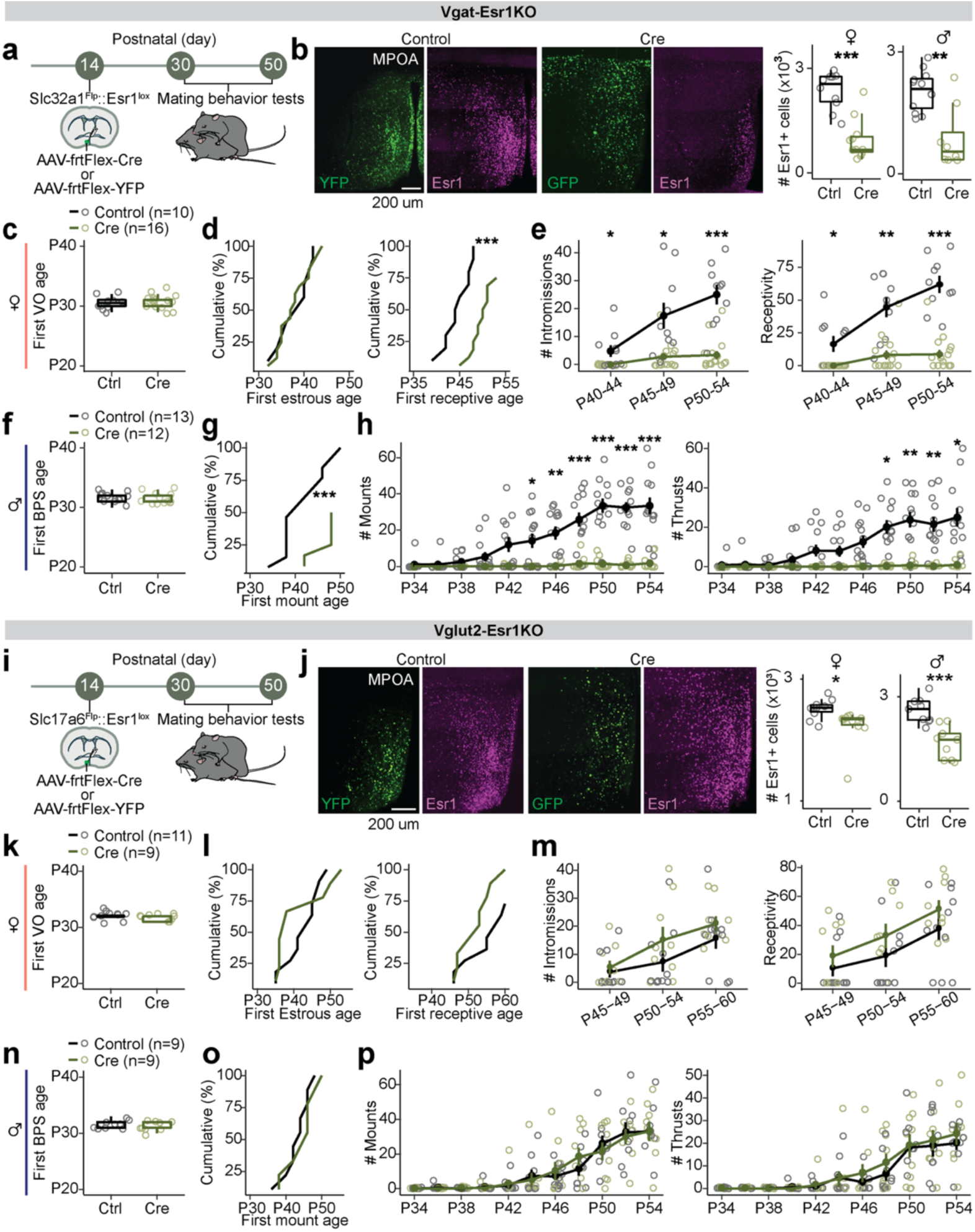
*Esr1* in MPOA^Vgat+Esr1+^ neurons governs adolescent maturation of sexual behaviors **a,** Schematic timeline for behavioral experiments testing the role of Esr1 in MPOA^Vgat+Esr1+^ during adolescent maturation of mating behaviors in *Slc32a1*^Flp^::*Esr1*^lox/lox^ mice (Vgat-Esr1KO), refers to panels **b-h**. **b,** Representative images of viral-reporter and Esr1 expression from Control (left) and Cre (right) groups. Quantification of Esr1 in the MPOA. **c-e,** Quantitative comparisons of female mice: first vaginal opening (VO) age (**c**), estrous age (**d**, left), receptive age (**d**, right), number of intromissions received (**e**, left), and receptivity (**e**, right). **f-h,** Quantitative comparisons of male mice: balanopreputial separation (BPS) age (**f**), first mount age (**g**), number of mounts (**h**, left) and thrusts (**h**, right). **i,** Schematic timeline for behavioral experiments testing the role of Esr1 in MPOA^Vglut2+Esr1+^ during adolescent maturation of mating behaviors in *Slc17a6*^Flp^::*Esr1*^lox/lox^ mice (Vglut2-Esr1KO), refers to panels **j-p**. **j,** Representative images of viral-reporter and Esr1 expression from Control (left) and Cre (right) groups. Quantification of Esr1 in the MPOA. **k-m,** Quantitative comparisons of female mice: first vaginal opening (VO) age (**k**), estrous age (**l**, left), receptive age (**l**, right), number of intromissions received (**m**, left), and receptivity (**m**, right). **n-p,** Quantitative comparisons of male mice: balanopreputial separation (BPS) age (**n**), first mount age (**o**), number of mounts (**p**, left) and thrusts (**p**, right). Line plots are shown in mean ± S.E.M and analyzed with a two-way repeated measures ANOVA followed by multiple comparisons. Box plots are shown with box (25%, median line, and 75%) and whiskers and analyzed with unpaired t-test. ***p < 0.001, **p < 0.01. *p < 0.05. Statistical details in Methods.

## Transcriptional dynamics of MPOA cell types during adolescent development

To characterize the transcriptional dynamics occurring in the MPOA specific to the adolescent period, we collected tissue from male and female wildtype mice at pre- (P23), mid- (P35) and post- (P50) adolescence timepoints, as well as from mice gonadectomized (GDX) prior to adolescence onset. We used droplet-based scRNAseq to recover the transcriptomes of 58,921 cells and combined the data from all samples across the 8 experimental groups (**Fig. 2a and Supplementary Fig. 2a-e and Supplementary Table 1**). Using known neuronal marker genes *Stmn2* and *Thy1*, we performed high-resolution clustering to focus our subsequent analyses on neuronal cell data (24,627 cells). Thirty-five neuron specific clusters were resolved with 434 marker genes in total (UMIs per cell: 5,395; median genes per cell: 2,592), resulting in 20 GABAergic clusters (Vgat+ clusters, 13,334 cells; 54.4% of neurons), 12 glutamatergic clusters (Vglut2+ clusters, 7,658 cells; 31.1% of neurons), and 3 clusters with mixed glutamatergic and GABAergic markers (3,635 cells; 14.8% of neurons) (**Fig. 2b and Supplementary Figs. 2f-l, 3a,j and Supplementary Table 2**), largely consistent with previously published data ^22,29^. To further quantify transcriptional associations between MPOA neuronal cell types, we performed partition-based graph abstraction (PAGA) analysis ^30^ to identify cell cluster similarities through a graphical representation of cell differentiation (**Fig. 2c**). Consistent with their distinct roles, Vgat+ and Vglut2+ neurons were readily separated in the PAGA graph. Additionally, Vgat+ clusters 2, 4 and 16, all of which expressed *Esr1*, showed higher connectivity compared to other Vgat+ cell type clusters (**Fig. 2c and Supplementary Figs. 2m, 3h-i**), suggesting that Vgat+Esr1+ clusters form a distinct transcriptional unit.

**Fig. 2:**
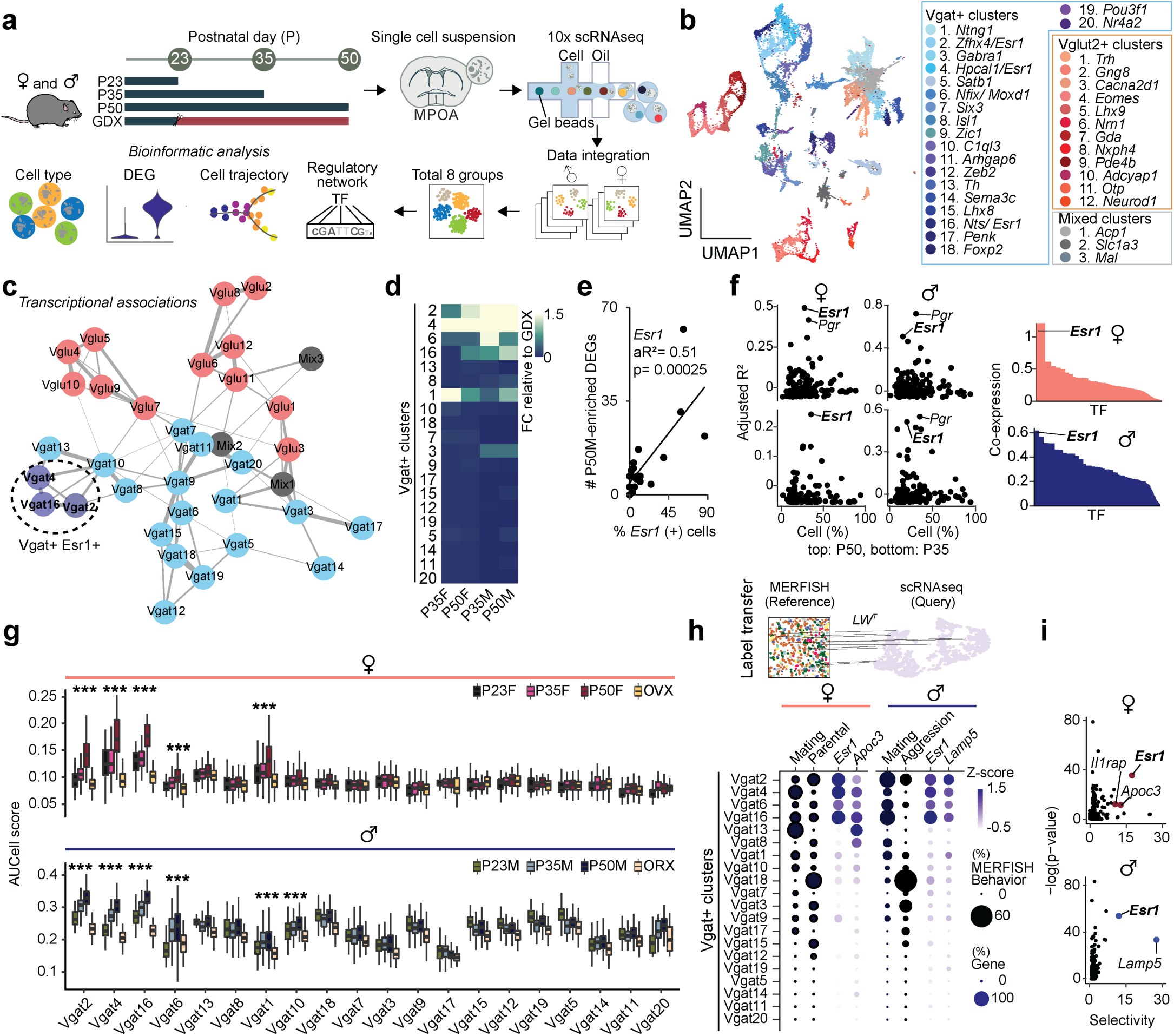
Single cell RNA sequencing identifies neuronal cell types across adolescence in the MPOA a, Schematic illustrating MPOA scRNAseq experiment methods. **b,** Joint clustering of 24,627 neurons from all groups (P23, P35, P50, GDX). Total identified clusters = 32. UMAP plot is color coded by neuronal cluster type, listed in legend. **c,** Visualization of neuron cluster transcriptional associations via PAGA analysis. Neuron cluster similarity is represented by the thickness of each connecting line. Vgat+Esr1+ clusters (Vgat 2, 4, & 16) demonstrate high connectedness and are highlighted within the dotted circle. Clusters: Pink = Vglut2+, Blue = Vgat+Esr1-, Purple = Vgat+Esr1+, Grey = Mixed. **d,** Heatmap showing the scaled sum of DEG log fold changes for each Vgat+ cluster in comparison to GDX samples. **e,** Linear regression analysis of percentage of *Esr1* expressing cells (x-axis) and the number of P50M-enriched DEGs in comparison with GDX samples (y-axis), where each dot is a Vgat+ cluster. **f,** Left: Scatter plots showing aR^2^ values of hormone receptor genes (dots) in females (left) and males (right) comparing the percentage of percent expressing cells to the number of DEGs in comparison with GDX samples at each Vgat+ cluster for P50-enriched (top) and P35-enriched (bottom) DEGs. Right: SCENIC analysis-computed ranked sums of TFs associated with HA-DEGs and their co-expression scores in females (top, 952 TFs) and males (bottom, 915 TFs). HA-DEGs in both sexes show highest co-expression with Esr1. Each TF is plotted along the x-axis in descending rank order. **g,** Quantitative comparison of aggregate HA-DEG expression (represented by AUCell score) within each Vgat+ cluster across P23, P35, P50, and GDX groups. **h,** Top: Schematic illustrating integrative analysis, establishing correspondence between MERFISH^22^ (Moffitt et al., 2018) and scRNAseq datasets. The MERFISH label transfer algorithm (details in Methods) identified behaviorally-relevant cells within the defined scRNAseq clusters. Bottom: The dot plot graph represents the percent of cells, indicated by dot size, identified as behaviorally-relevant (selected female behaviors: mating & parental; selected male behaviors: mating & aggression) in each Vgat+ cluster. Data integration revealed the top two marker genes related to mating as *Esr1* and *Apoc3* (female, left) and *Esr1* and *Lamp5* (male, right) (see panel **i**). The dot plot graph also represents the percent of cells, indicated by dot size, expressing the marker gene in each Vgat+ cluster, in addition to its z-scored gene expression indicated by dot color intensity. **i,** Scatter plots of enriched genes in mating-related scRNAseq clusters in females (top) and males (bottom). Box plots are shown with box (25%, median line, and 75%) and whiskers and analyzed with Kruskal-Wallis H test followed by multiple comparisons test. p-values were Bonferroni corrected. ***p < 0.001. Statistical details in Methods. aR^2^: adjusted R squared; GDX: gonadectomy; OVX: ovariectomy; ORX: orchiectomy.

**Fig. 3:**
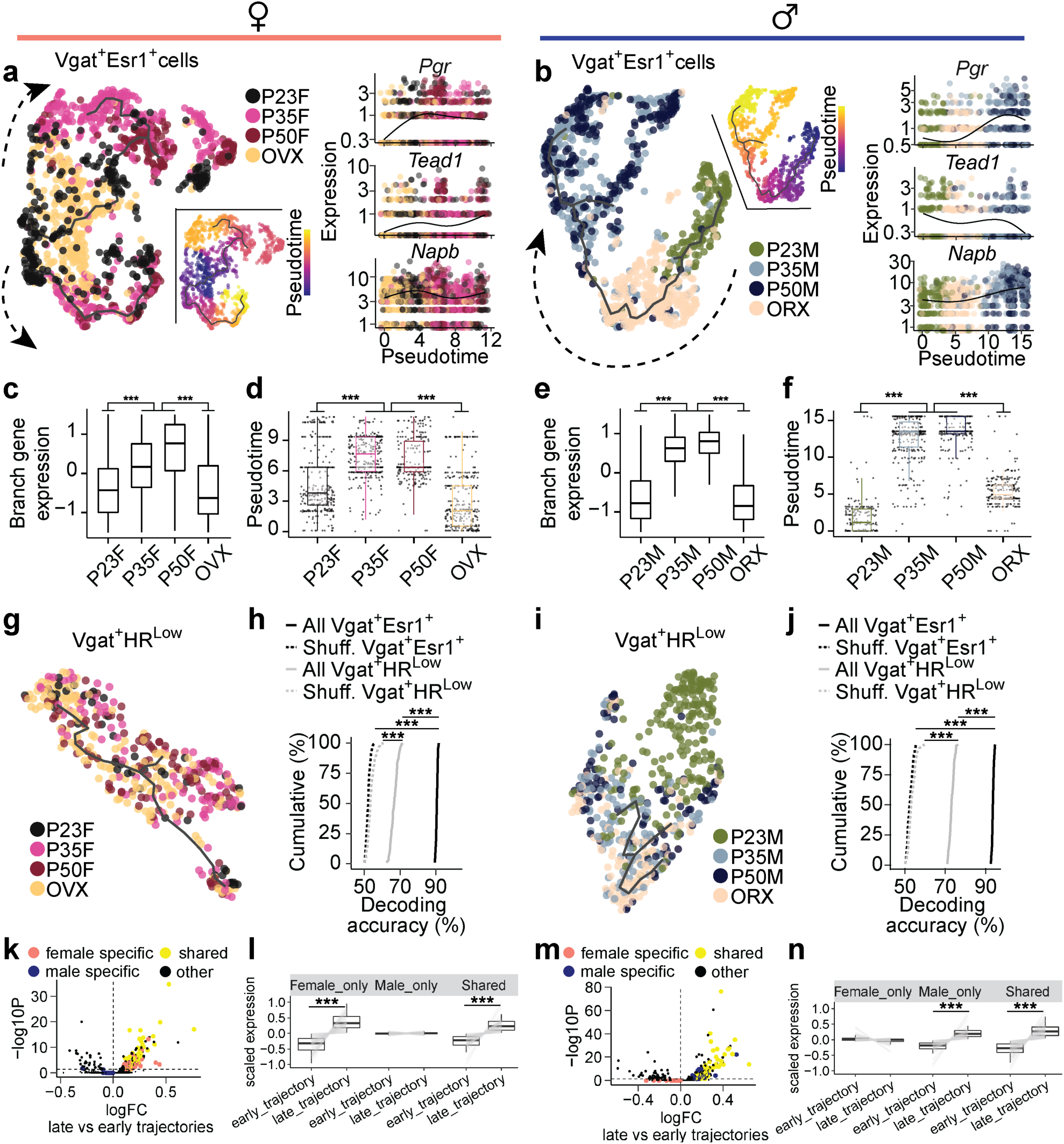
Identification of adolescent transcriptional trajectories in MPOA^Vgat+Esr1+^ neurons **a-b,** UMAP visualization of Vgat+Esr1+ cells and their transcriptional trajectories depicted by a solid black line. Vgat+Esr1+ cells are color coded by group (left) and pseudotime (right), where progression of time is delineated from dark to bright coloring. Dashed arrows indicate the direction of transcriptional progression. Kinetics plots show the relative expression of HA-DEGs *Pgr* (shared), *Tead1* (female-related), and *Napb* (male-related) in Vgat+Esr1+ cells for each group across adolescent pseudotime (x-axis). **a**: females; **b**: males. **c, e,** Box plots showing scaled gene expression of Vgat+Esr1+ branch enriched genes in female (**c**) and male (**e**) mice. **d, f,** Box plots showing Vgat+Esr1+ cell placement along pseudotime for each group. **g, i,** UMAP visualization of Vgat+HR^Low^ cells and their transcriptional trajectories depicted by a solid black line. Vgat+HR^Low^ cells are color coded by group in females (**g**) and males (**i**). **h, j,** Cumulative distributions of decoding accuracy by SVM classification between mature groups (P50, P35) and immature groups (P23, GDX) using expression data from Vgat+Esr1+ (black, real line), Vgat^+^ HR^Low^ (grey) or shuffled data (dashed line) in females (**h**) and males (**j**). **k, m,** Volcano plots comparing gene expression between late and early trajectories in female (**k**) and male (**m**). Sex-specific and shared gene programs are highlighted (shared: yellow; female-specific: salmon; male-specific: blue). **l, n,** Box and line plots showing scaled expression of sex-specific and shared enriched genes from early to late trajectories in females (**l**) and males (**n**). Box plots are shown with box (25%, median line, and 75%) and whiskers and analyzed with Kruskal-Wallis H test followed by multiple comparisons test with p-values Bonferroni corrected (**c-f**) or Wilcoxon rank-sum test (**l, n**). Cumulative distributions were analyzed one-way ANOVAs followed by multiple comparisons. ***p < 0.001. OVX: ovariectomy; ORX: orchiectomy; GDX: gonadectomy; HR^Low^: hormone receptor-low; SVM: support vector machine.

Irrespective of age, sex, and hormonal states, we found that each identified cell type was represented across all 8 experimental groups (**Supplementary Figs. 2d,h-i, 3a**), indicating that new cell types do not emerge during adolescence. However, when assessing gene expression differences across timepoints and sex, we found that hormone-associated differentially expressed genes (HA-DEGs) showed substantial variability across neuronal cell types, ranging from 0 to 88 HA-DEGs per cell type (**Fig. 2d-f and Supplementary Fig. 3a-g and Supplementary Tables 3-5**). We then used these HA-DEGs in regression, co-expression, and AUCell analyses to identify the neuronal types that are transcriptionally sensitive at different hormonal states ^31^. Regression analysis revealed a strong enrichment of HA-DEGs in Vgat+Esr1+ neurons (**Fig. 2e-f)**. Consistent with this result, co-expression analysis demonstrated that out of more than 1,000 transcription factors (TFs), *Esr1* was one of the most co-expressed in both sexes (**Fig. 2f and Supplementary Table 6)**. Furthermore, in predominantly Vgat+Esr1+ clusters, the aggregate expression of HA-DEGs (AUCell) displayed patterns sensitive to both adolescence and hormonal changes, with the highest expression observed in adult mice (P50) and notably lower expression at P23 and during hormonal deprivation (GDX) (**Fig. 2g**). Together, these analyses demonstrate that sex-hormone signaling during adolescence development significantly influences the transcriptional states of specific MPOA neuronal cell types, particularly cells belonging to Vgat+Esr1+ clusters.

Having identified the distinct transcriptional sensitivity of Vgat+Esr1+ neurons to hormone signaling during adolescence, we next examined whether these neurons might play a direct role during behavior. Given that mating behavior was abolished in MPOA^Vgat-Esr1KO^ mice in both sexes (**Fig. 1**), this finding, along with our scRNAseq analyses, suggests that Vgat+Esr1+ neurons are critically involved during mating behaviors. To test this hypothesis, we conducted an integrative analysis using our scRNAseq dataset and a publicly available multiplexed error-robust fluorescence *in situ* hybridization (MERFISH) dataset ^22^ which identified social behavior-activated POA cells via mating, parenting, and fighting behavior assays using *Fos* (**Fig. 2h**). We clustered the neuronal cells in the MERFISH data and computed the enrichment of *Fos* in the derived clusters. Consistent with previous observations, only a subset of MERFISH Gad1*+* inhibitory clusters showed *Fos* enrichment (**Supplementary Fig. 4a,b**). After integrating our scRNAseq clusters with the MERFISH clusters (Details in Methods. Seurat V3 Integrative Analysis: reference-based integration; label transfers ^32^) we were able to identify mating, parenting, and fighting-relevant neuronal clusters within our dataset (**Fig. 2h and Supplementary Fig. 4c-d and Supplementary Table 7**). We found that *Esr1* was the most enriched and selective gene in the mating-relevant scRNAseq Vgat+ clusters (**Fig. 2h-i and Supplementary Table 7**). This observation is consistent with a previous study demonstrating that optogenetic stimulation of Vgat+Esr1+ neurons in the MPOA is sufficient to induce sexual behaviors in male mice ^18^. Collectively, these findings suggest that a subset of Vgat+ clusters expressing *Esr1*, which exhibit transcriptional changes during adolescence, are also functionally linked to mating behaviors in both males and females.

While these differential gene expression (DE) analyses quantify changes in individual gene expression at the pseudo-bulk level (**Fig. 2d and Supplementary Fig. 3b-c**), it does not capture the transitions in transcriptional states of individual cells as they progress through distinct biological stages. These transitions, driven by complex interactions between multiple genes, are key to understanding how single cells change over time ^33^. To resolve these dynamic changes, we applied pseudotime analysis (**Fig. 3 and Supplementary Figs. 5-6**), a method that infers the transcriptional progression of individual cells through a biological process, such as adolescence, by constructing a principal graph based on combinatorial gene expression patterns ^34,35^.

Recognizing the potential for sex-specific differences in transcriptional progression, we created separate pseudotime manifold models for males and females to capture nuanced dynamics during adolescence ^36^. In both sexes, Vgat+Esr1+ trajectories revealed that transcriptional states at P35 and P50 branch and diverge from preadolescence **(Fig. 3a-f and Supplementary Fig. 5a-f)**, suggesting an acceleration of transcriptional dynamics between these timepoints. Neurons from GDX mice, on the other hand, showed arrested transcriptional progression, closely resembling preadolescent states, demonstrating the importance of circulating sex hormones in the maturation of these cells during adolescence. In contrast, Vgat+ cells lacking steroid hormone receptor gene expression (Vgat+ hormone R^Low^) exhibited minimal transcriptional changes in response to hormone changes (**Fig. 3g,i and Supplementary Fig. 6a-c**). Pair-wise DEG analysis consistently showed that larger number of DEGs between P35 and P23 in Vgat+Esr1+ (male: 146 genes; female: 162 genes) than Vgat+ hormone R^Low^ (male: 26 genes; female: 1 gene). Furthermore, all Vgat+Esr1+ clusters were found to co-express the androgen receptor gene (*Ar*) to a high degree (68.3 – 88.3 %, **Supplementary Fig. 6a**). However, Vgat+ clusters expressing *Ar* but not *Esr1* (Vgat+ Esr1- AR+) showed only a moderate transcriptional progression in males and minimal changes in females (**Supplementary Fig. 6b-d**), suggesting that *Esr1* is more influential in driving transcriptional changes during adolescence than *Ar* alone. Lastly, we analyzed Vglut2+ clusters enriched with *Esr1*, and again found only moderate transcriptional progression in both sexes (**Supplementary Fig. 7**), altogether indicating that cell types outside of Vgat+Esr1+ clusters are less transcriptionally dynamic during this period.

Two major branches of transcriptional trajectories emerged from subpopulations of Vgat+Esr1+ cells in both sexes. In females, Branch1 was dominated by Vgat+ cluster 4, while Branch2 was occupied by clusters 2 and 16. In males, Branch1 was composed of Vgat+ clusters 4 and 16, while Branch 2 was primarily formed by Vgat+ cluster 2 (**Supplementary Fig. 5g-j**). The analysis of DEGs along these branches revealed distinct gene enrichment, indicating that specific subpopulations of Vgat+Esr1+ cells undergo unique transcriptional progressions during adolescence (**Supplementary Fig. 5c-d and Supplementary Table 8**). When we combined the trajectories for both sexes into a joint manifold, the model failed to capture hormone-dependent dynamics (**Supplementary Fig. 6e-g**), emphasizing that separate models are necessary for accurate analysis of sex-specific and sex-shared transcriptional progressions (**Fig. 3k-n**).

To strengthen our understanding of the developmental trajectory and transcriptional maturity of Vgat+Esr1+ clusters, we employed a support vector machine (SVM) classifier to predict the developmental state of single cells based solely on HA-DEGs ^37^. The SVM accurately classified Vgat+Esr1+ single cells as either transcriptionally mature (intact adolescent or adult) or immature (preadolescent or hormonally deprived) with high accuracy (median prediction accuracy 90.6 – 93.6 %). In contrast, Vgat+ hormone R^Low^ cells had significantly lower prediction accuracy (median 66.7 - 73.0 %) (**Fig. 3h,j**), further underscoring the importance of Vgat+Esr1+ clusters in the development of MPOA transcriptional states.

Finally, we validated our pseudotime trajectories (Monocle V3) using Manifold Enhancement Latent Dimension (MELD) analysis, which measures continuous transcriptional progression in response to experimental conditions (P23, P35, P50, GDX) ^38,39^. MELD analysis corroborated our findings from pseudotime, showing that intact adolescents and adults were spatially segregated in transcriptional space and exhibited a higher likelihood of reaching mature transcriptional states compared to preadolescent and hormonally deprived groups (**Supplementary Fig. 6h-m**). These results, visualized in Potential of Heat-diffusion for Affinity-based Trajectory Embedding (PHATE) space ^38^, further highlight that while MPOA neuronal cell types are terminally diversified by preadolescence (P23) (**Supplementary Figs. 2h, 3e**), Vgat+Esr1+ neurons continue to undergo hormonally dependent transcriptional refinement from adolescence into adulthood.

## Spatial phenotyping of MPOA cell types during adolescence

To assess whether adolescent hormonal changes affect spatially resolved gene expression in the MPOA and to cross-validate our scRNAseq trajectory analyses, we performed highly multiplexed hybridization chain reaction fluorescent *in situ* hybridization (HM-HCR FISH, V3)^40,41^. This technique allowed us to detect transcripts of ∼12 genes across 41,549 MPOA cells at single-molecule resolution (**Fig. 4a-c and Supplementary Figs. 8-9).** The rationale for selecting this gene panel is related to **Supplementary Fig. 6k,m,** with details provided in the Methods. In addition to the experimental groups used in the scRNAseq experiments, we prepared tissue from hormonally supplemented mice. These mice received testosterone (males) or estrogen (females) from P23-P27 (see timeline in **Fig. 4a**) to investigate whether early sex steroid supplementation accelerated adolescent transcriptional trajectories.

**Fig. 4:**
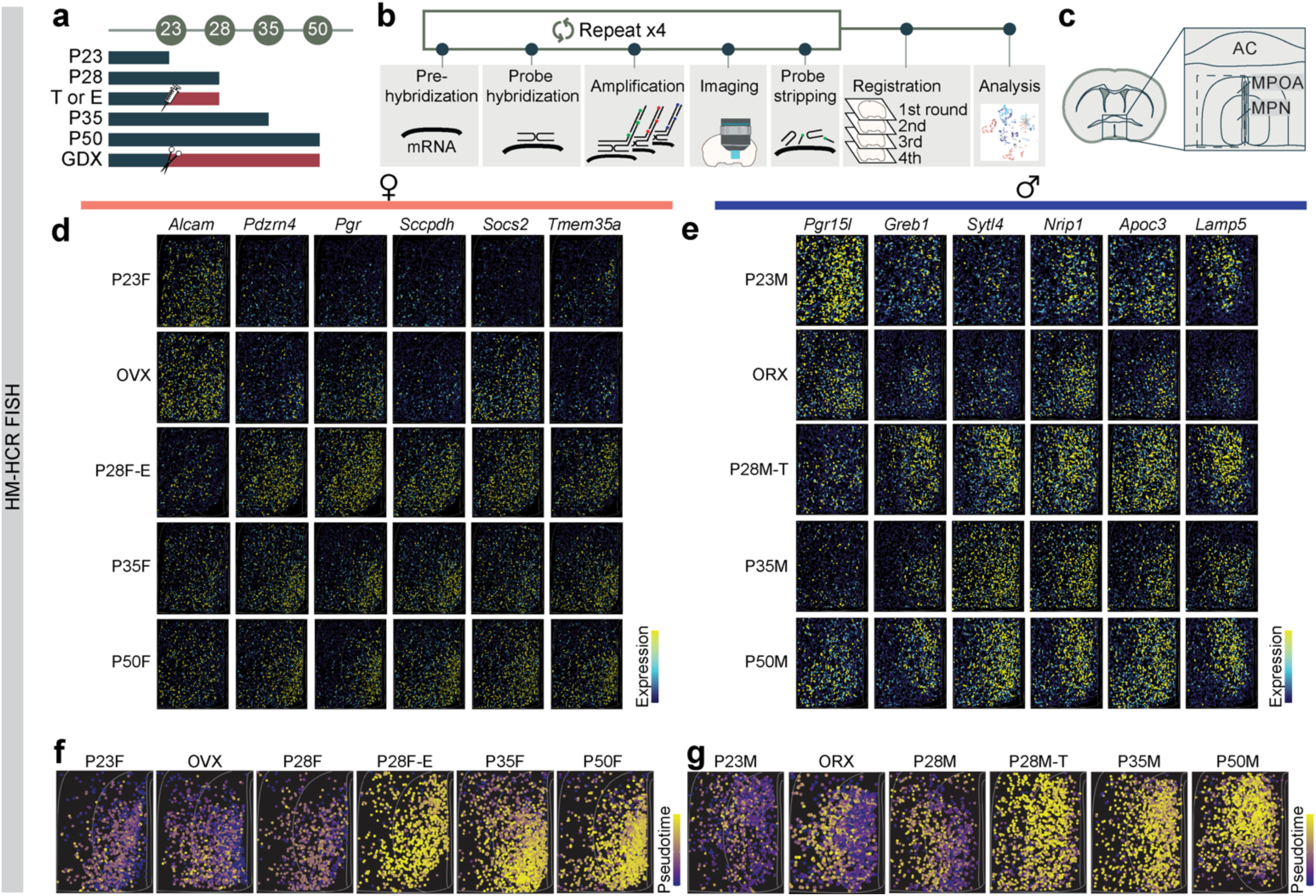
HM-HCR FISH reveals spatial transcriptional trajectories of MPOA^Vgat+Esr1+^ neurons across adolescence and hormonal state **a,** Schematic timeline for highly multiplexed-HCR FISH (HM-HCR FISH) experiments with tissue collected at different combinations of ages and hormone manipulation. **b,** Schematic illustrating HM-HCR FISH procedure. **c,** Schematic of a representative mouse coronal brain section for analyzing MPOA. A total of 41,549 cells from MPOA were analyzed. **d, e,** Representative images of reconstructed cells in MPOA, color coded by scaled expression of genes at P23, GDX, P28 with hormone supplementation, and P50. **d:** females; **e:** males. Quantitative analysis of each gene is reported in **Supplementary Fig. 9a-b, i-j**. **f, g,** Pseudotime spatial visualization of Vgat+Esr1+cells across all 6 groups in females (**f**) and males (**g**), where progression of time is delineated from dark to bright coloring. Pseudotime score was computed using HM-HCR FISH gene expression data, described in detail in Methods. Quantitative comparisons of this data is reported in **Supplementary Fig. 9r-u**. T: testosterone; E: estradiol; AC: anterior commissure; MPOA: medial preoptic area; MPN: medial preoptic nucleus; GDX: gonadectomy; OVX: ovariectomy; ORX: orchiectomy.

To distinguish Vgat+Esr1+ from Vgat+ hormone R^Low^ cells, we measured *Slc32a1*, *Esr1, Ar,* and 8-9 DEGs during adolescence (**Fig. 4d-e and Supplementary Figs. 8-9.** Details in Methods). Consistent with the scRNAseq data, we observed a transcriptional progression from the preadolescent state (P23) to a matured state (P35 and P50), where the dynamics were bidirectionally influenced by circulating steroids– accelerated by early testosterone or estrogen supplementation (P28 T or E2) and delayed in a hormonally-deprived state (GDX) (**Fig. 4f-g and Supplementary Fig. 9a-c,f,i-q**). Interestingly, the transcriptional trajectories of female and male adolescents differed spatially, but both were regulated by sex hormones **(Supplementary Fig. 9r-u).** This suggests that neurons defined transcriptionally as adult female or male occupy partially overlapping spatial distributions in the MPOA. SVM classification revealed that the expression of ≥10 genes in Vgat+Esr1+ cells from the HM-HCR FISH data was sufficient to accurately classify individual cells as transcriptionally mature or immature, with high accuracy (84.4–85.5 %). However, the same set of genes significantly underperformed in classifying Vgat+ hormone R^Low^ cells (63.6–64.6 %) (**Supplementary Fig. 9d,g**). Classification accuracy further decreased when individual genes were iteratively removed (**Supplementary Fig. 9e,h**), further indicating that a relatively small combinatorial set of HA-DEGs is sufficient for accurately identifying age- and sex-specific transcriptional states.

Molecular profiling of the MPOA in adult mice has established the presence of sexual dimorphism ^42,43^. Our scRNAseq and *in situ* analyses further show hormone-dependent transcriptional progression during adolescence in both females and males. However, critical questions remain about whether (1) sexual dimorphism is evident within specific MPOA cell types, (2) sexually dimorphic genes overlap with adolescent gene sets, and (3) the degree of sexual dimorphism shifts during adolescence. To address these questions, we first identified sexually dimorphic genes across Vgat+ clusters (**Supplementary Fig. 10a and 11**). Like HA-DEGs, sexually dimorphic genes were enriched within Vgat+Esr1+ clusters. The number of these dimorphic genes correlated with *Esr1* expression, where sexually dimorphic genes co-expressed most often with *Esr1* in both sexes (**Fig. 5a-e and Supplementary Fig. 10a and Supplementary Table 6**). Notably, only subsets of sexually dimorphic genes were also classified as adolescent genes (**Figs. 5f, 6a-b**) and some of these adolescent genes were shared across sexes (**Fig. 5g**).

**Fig. 5:**
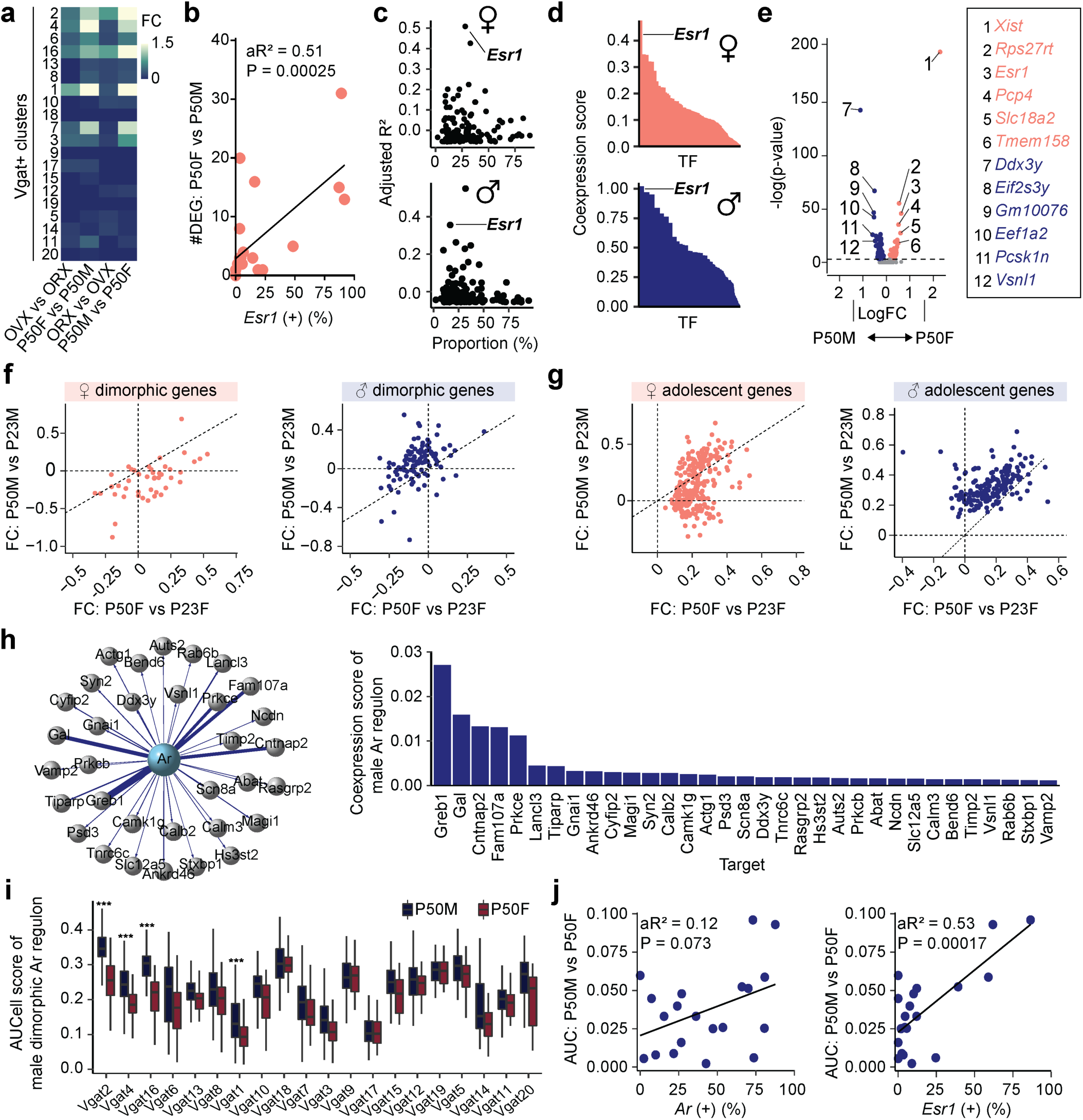
Identification of sexually dimorphic genes in MPOA **a,** Heatmap representing the scaled sum of log fold changes of sexually dimorphic genes for each Vgat+ cluster comparing females and males (bidirectionally) in hormonally-intact P50 mice or GDX mice. **b,** Linear regression analysis comparing the percentage of *Esr1* expressing cells and the number of female dimorphic genes within each Vgat+ cluster (individual dots). **c,** Scatter plots showing adjusted R^2^ values of hormone receptor genes (dots) in females (top) and males (bottom) as a result of linear regression analysis comparing the percentage of hormone receptor gene expressing cells and number of sexually dimorphic genes across Vgat+ clusters. **d,** SCENIC analysis-computed ranked sums of TFs associated with sexually dimorphic genes and their co-expression scores in females (top) and males (bottom). Sexually dimorphic genes in both sexes show highest co-expression with Esr1. Each TF is plotted along the x-axis in descending rank order. **e,** Volcano plot comparing P50M and P50F gene expression in Vgat+Esr1+ cells. Dimorphic genes are numbered and color coded (P50F-rich: salmon; P50M-rich: blue). **f, g,** Scatter plots showing fold change differences of sexually dimorphic genes (**f**) or adolescence-related genes (**g**) between P50F and P23F (x-axis) and P50M and P23M (y-axis). Female-rich genes plotted on the left and male-rich genes plotted on the right. Adolescent genes and dimorphic genes were only partially overlapping. Adolescent genes were partially shared between sexes. **h,** Motif-enrichment analysis of male-rich dimorphic genes reveals deconstructed *Ar*-regulons. The co-expression score between *Ar* and a regulon gene is represented by the thickness of their connecting line in the visualization (left) and via a bar graph (right). **i**, Box plots comparing P50M to P50F expression of male-rich dimorphic *Ar*-regulon genes at each Vgat+ cluster via AUCell analysis. **j,** Linear regression analysis between the percentage of *Ar* (left) or *Esr1* (right) expressing cells and AUCell score of male-rich dimorphic *Ar*-regulon genes at each Vgat^+^ cluster (each dot). Box plots are shown with box (25%, median line, and 75%) and whiskers and analyzed with Wilcoxon rank-sum test. p-values were Bonferroni corrected. ***p < 0.001. Statistical details in Methods. aR^2^: adjusted R squared; TF: transcription factor; FC: fold change; Ar: androgen.

**Fig. 6:**
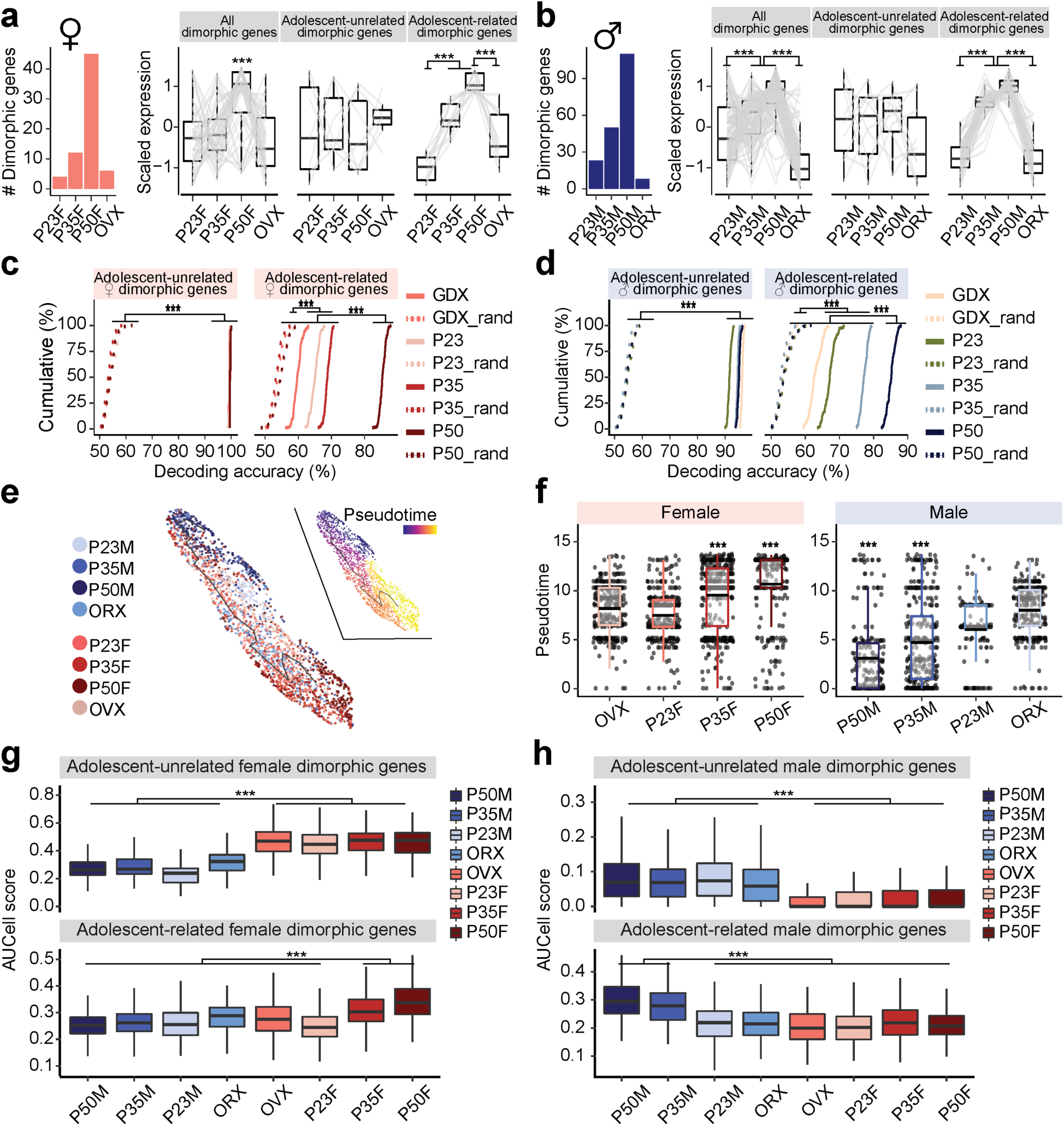
A**d**olescent dynamics of sexually dimorphic genes in MPOA^Vgat+Esr1+^ neurons **a, b,** Bar plot of adult dimorphic genes in females (**a**) and males (**b**) in each group. Box plots show scaled gene expression of sexually dimorphic genes (left: all; middle: adolescent-unrelated; right: adolescent-related) within Vgat+Esr1+ cells in females (**a**) and males (**b**). **c, d,** Cumulative distributions of decoding accuracy by SVM classification for adolescent-unrelated (left) and -related (right) dimorphic genes between females and males of matched groups using female (**c**) or male (**d**) dimorphic genes. Shuffled data depicted with dashed lines. **e,** UMAP transcriptional trajectory visualization of combined female and male Vgat+Esr1+ cells (dots), color coded by group (left) or pseudotime (right) where progression of time is delineated from dark to bright coloring. **f,** Box plots showing pseudotime assignment of Vgat+Esr1+ cells across groups in females (left) and males (right). **g, h,** Box plots comparing AUCell aggregate expression of female dimorphic (**g**) or male dimorphic (**h**) genes (top: adolescent-unrelated; bottom: adolescent-related) across groups. Box plots are shown with box (25%, median line, and 75%) and whiskers and analyzed with Kruskal-Wallis H test followed by multiple comparisons test. p-values were Bonferroni corrected. Cumulative line graphs were analyzed with one-way ANOVAs followed by multiple comparisons. ***p < 0.001. Statistical details in Methods. GDX: gonadectomy; OVX: ovariectomy; ORX: orchiectomy; SVM: support vector machine.

Previous research has shown that *Esr1* plays a role in regulating sexual dimorphisms within sexually dimorphic brain nuclei ^23^. However, androgen receptor (AR) activation may also be essential for the expression of sexually dimorphic genes. To determine whether AR regulates male dimorphic genes, we performed single-cell regulatory network inference (SCENIC) analysis ^31^. SCENIC identified a regulon of 33 AR-regulated genes, which were highly co-expressed with *Ar* and enriched with *Ar*’s consensus DNA regulatory element within their gene loci, in the male dimorphic gene set (**Fig. 5h**). AUCell analysis, however, showed that male dimorphic AR-regulon genes were enriched within male Vgat+Esr1+ clusters but not in Vgat+ Esr1- Ar+ clusters (specifically Vgat+ 5, 8, and 13) (**Fig. 5i**). Consistent with this observation, regression analysis showed that *Esr1* expression, rather than *Ar*, predicted the expression of male dimorphic AR-regulon genes (**Fig. 5j**).

As previously noted, a subset of sexually dimorphic genes overlaps with adolescent-related genes (**Fig. 5f**). The number of sexually dimorphic genes increases during adolescence but reverts following gonadectomy (**Fig. 6a-b**). Among 45 female dimorphic genes, 12 showed increased expression during adolescence (adolescence-related dimorphic genes), while 7 were not associated with adolescence (**Fig. 6a, right**). Of the 110 dimorphic genes in males, 52 were adolescence-related dimorphic genes and 10 were not (**Fig. 6b, right**). The expression levels of adolescence-related dimorphic genes were the highest at P50 and were again reduced by gonadectomy in both sexes (**Fig. 6a-b**). SVM analysis revealed that adolescence-related dimorphic genes decoded sex with the highest accuracy in intact P50 males and females, followed by P35, and were less accurate in P23 and GDX groups. In contrast, adolescence-unrelated dimorphic genes successfully predict the sexes of single cells with high accuracy (>90%) irrespective of age and conditions (**Fig. 6c-d**), emphasizing the influence of adolescence-related dimorphic genes in shaping MPOA sex-specific transcriptional profiles. AUCell and trajectory analyses were then performed to quantify the transcriptional dynamics of sexual dimorphic genes during adolescence. These analyses consistently demonstrated that transcriptional states were the most sexually dimorphic at P50 driven by adolescence-related dimorphic genes and were the least dimorphic at P23 and GDX (**Fig. 6e-h**). Thus, combined trajectory analysis of all Vgat+Esr1+ neurons from both sexes, along with AUCell analysis, revealed that P50F and P50M cells were the most sexually dimorphic from each other. However, these transcriptional states were bridged via a common immature state observed largely prior to adolescence onset or in hormonally depleted conditions, suggesting that sexually dimorphic MPOA gene expression is bimodal in adults but largely continuous between males and females prior to adolescence. These analyses (**Figs. 5 and 6**) indicate that: 1) sexual dimorphism is most pronounced in Vgat+Esr1+ cells; 2) a subset of sexually dimorphic genes are linked to adolescence, with a fraction of adolescence-related genes shared between sexes; and 3) sexual dimorphism in the MPOA expands during adolescence as sex steroid hormone levels rise.

### *Esr1* knockout at preadolescence alters the transcriptional dynamics of Vgat+ MPOA neurons

Co-expression of *Esr1* with adolescence genes and dimorphic genes, along with partially distinct transcriptional dynamics in Vgat+Esr1+ cells between adolescent females and males, suggests that *Esr1* uniquely regulates gene expression in each sex. To further test this, we virally deleted *Esr1* from Vgat+ MPOA neurons in female and male mice prior to the onset of adolescence and conducted scRNAseq at P50, comparing Esr1KO to Esr1-intact controls (**Fig. 7a and Supplementary Fig. 11.** Details in Methods). Esr1KO in Vgat+ MPOA neurons reduced the DEGs in males and females (Esr1-DEGs) to 600 and 824, respectively, while only reducing 5.3% and 3.3% of male and female Esr1-DEGs in Vgat+ hormone R^Low^ cells (**Fig. 7b and Supplementary Fig. 10c and Supplementary Table 9**). Previously identified HA-DEGs (**Fig. 2**) were largely represented in this list of Esr1-DEGs, suggesting that *Esr1* deletion recapitulates hormone deprivation effects on the transcriptomes of Vgat+Esr1+ cells (males: 56.5%, 78/138 genes; females: 73.3%, 124/169 genes; **Fig. 7b**). Again, because of the complex nature of this relationship between dimorphic and adolescent genes, we generated a separate manifold for each sex to assess MPOA adolescent transcriptional dynamics. Pseudotime trajectory analysis indicated that Esr1KO near-completely prevented adolescent transcriptional maturation in both sexes (**Fig. 7c-h and Supplementary Fig. 12**). DE analysis also identified sex-shared and -specific processes (**Fig. 7i-l and Supplementary Table 10**). In addition, dimorphic genes in the Esr1-DEGs of Vgat+Esr1+ cells were sufficient to predict the sex in P50 females and males with the highest accuracy (81.4 ± 0.11 %), followed by P35, and lowest accuracy in P23 (**Supplementary Fig. 10d**). Like our previous observation (**Supplementary Fig. 6**), we observed two major branches of transcriptional trajectories with branch specific gene expression profiles, primarily originating from one to two subpopulations of Vgat+Esr1+ clusters (**Supplementary Fig. 12c-d and Supplementary Table 10**) highlighting *Esr1* as a key regulator of adolescent transcriptional dynamics in the MPOA of both sexes.

**Fig. 7:**
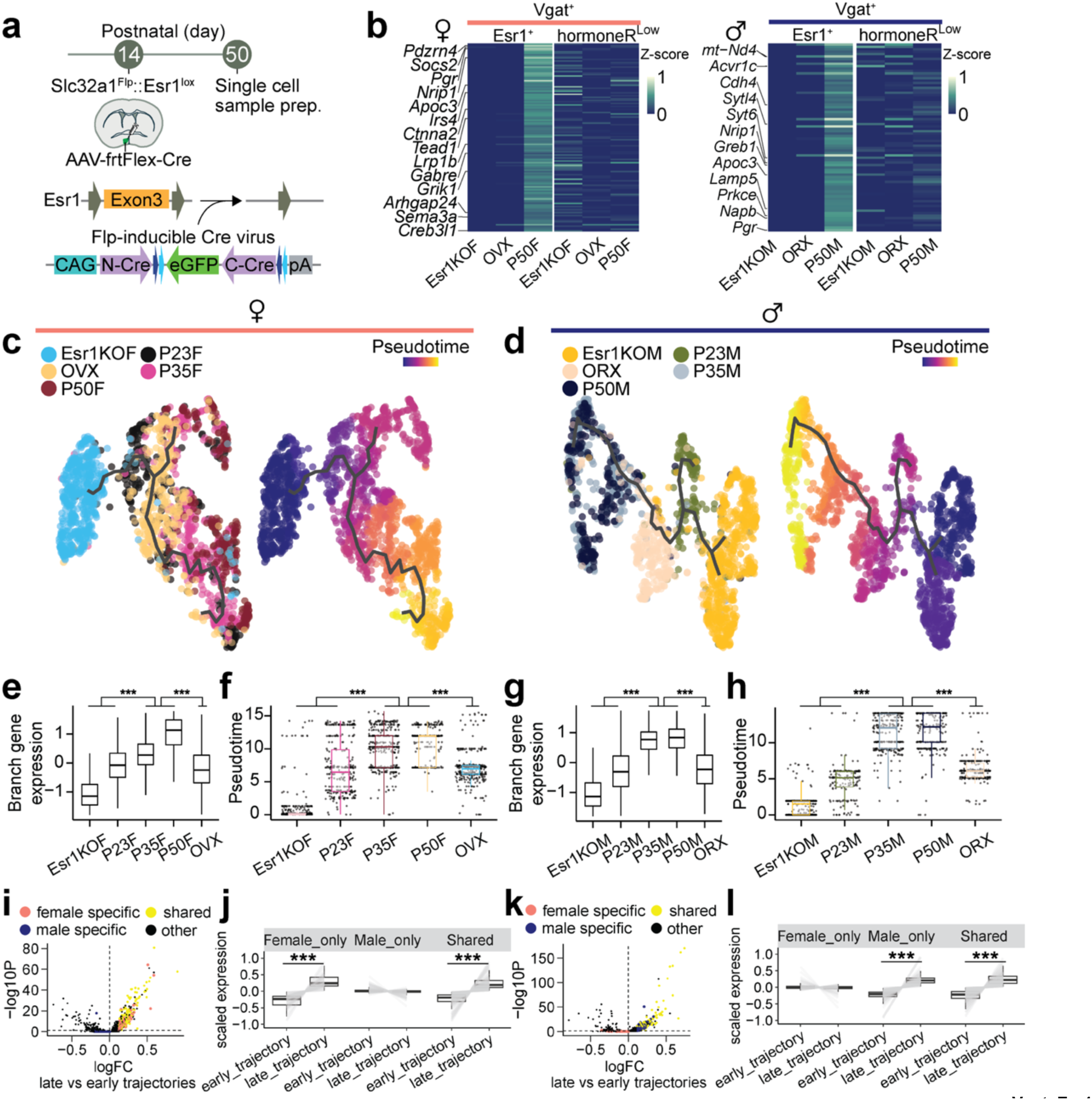
Regulation of adolescent transcriptional dynamics via *Esr1* activation in MPOA^Vgat+Esr1+^ neurons **a,** Schematic illustrating cell type and site-specific deletion of *Esr1* in MPOA using viral vector AAV-frtFlex-Cre in *Slc32a1*^Flp^::*Esr1*^lox/lox^ mice, followed by scRNAseq. **b,** Heatmaps show the scaled average expression (z-score) of Vgat+ HA-DEGs for Esr1+ and hormoneR^Low^ cells across Esr1KO, GDX, and P50 groups for females (left) and males (right). **c, d,** UMAP visualization of Vgat+Esr1+ cells and their transcriptional trajectories depicted by a solid black line in females (**c**) and males (**d**). Vgat+Esr1+ cells are color coded by group (left) and pseudotime (right), where progression of time is delineated from dark to bright coloring. **e, g,** Box plots showing scaled gene expression of Vgat+Esr1+ branch enriched genes for each group in female (**e**) and male (**g**) mice. **f, h,** Box plots showing Vgat+Esr1+ cell placement along pseudotime for each group. **i, k,** Volcano plot comparing gene expression between late and early trajectories in female (**i**) and male (**k**). Sex-specific and shared gene programs are highlighted (shared: yellow; female-specific: salmon; male-specific: blue). **j, l,** Box and line plots showing scaled expression of sex-specific and shared enriched genes from early to late trajectories in females (**j**) and males (**l**). Box plots are shown with box (25%, median line, and 75%) and whiskers and analyzed with Kruskal-Wallis H test followed by multiple comparisons test. p-values were Bonferroni corrected. ***p < 0.001. Statistical details in Methods. HA-DEG: hormone-associated differentially expressed gene; GDX: gonadectomy; OVX: ovariectomy; ORX: orchiectomy.

Gene regulatory networks (GRNs) are complex systems of genes, transcription factors, and other molecules that interact to control gene expression within a cell. To identify GRNs that are shared between sexes or specific to one sex, we used functional-SCENIC to examine if *Esr1* cis-regulates its own differentially expressed genes (Esr1-DEGs) and/or transcription factors (Esr1-TFs) (**Fig. 8a**). Our analysis revealed that a significant portion of Esr1-DEGs contained *Esr1* binding sites–14.9% in females (123 out of 824 genes) and 16.3% in males (98 out of 600 genes) (**Fig. 8b-c**)–indicating that *Esr1* directly regulates many target genes. Additionally, 9 female and 6 male Esr1-DEGs encoded transcription factors that function as central regulatory hubs within the network (**Fig. 8d-e**), suggesting that Esr1’s influence extends indirectly through these secondary transcription factors. 54.1% of female and 28.3% of male Esr1-DEGs were cis-regulated by Esr1-TFs, and collectively, 56.1 % female and 36.7% male Esr1-DEGs were cis-regulated by *Esr1*, Esr1-TFs, or a combination of the two (Esr1-GRNs; **Fig. 8d-e and Supplementary Fig. 13**).

**Fig. 8:**
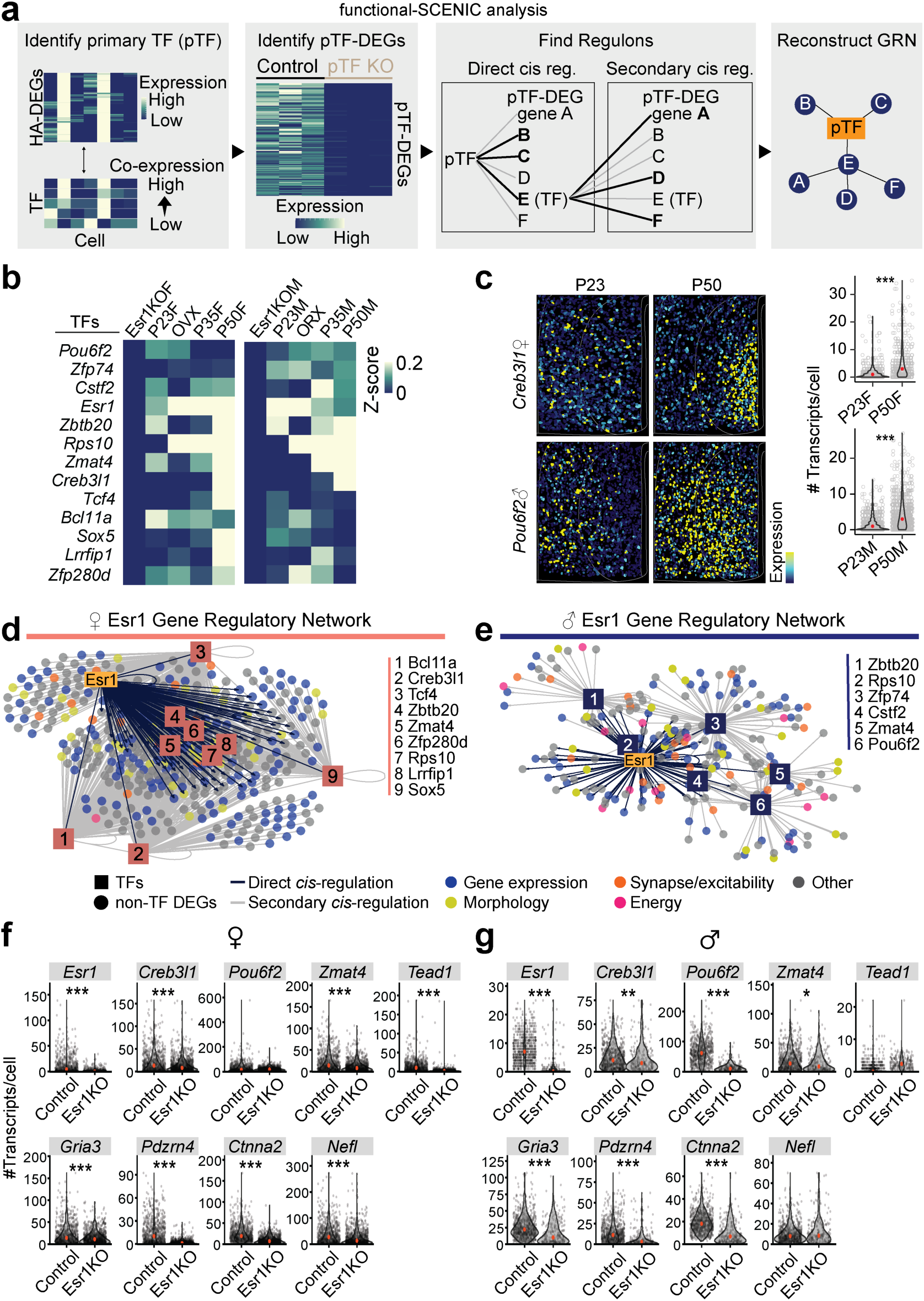
D**e**constructing sex-specific gene regulatory networks underlying adolescent transcriptional dynamics in MPOA^Vgat+Esr1+^ neurons **a,** Schematic demonstrating the deconstruction of causal GRNs through the combination of functional genomics and SCENIC analysis. (Left to right) Functional-SCENIC analysis involves identification of pTFs that co-express previously identified HA-DEGs, then those pTF-DEGs are analyzed for regulons and sorted into primary and secondary cis-regulated genes. With that data, a GRN can be reconstructed for each pTF. **b,** Heatmap for Vgat+Esr1+ cells scaled expression (z-score) of *Esr1* and Esr1-regulated TFs in Esr1KO, P23, GDX, P35, and P50 groups for females (left) and males (right). **c,** Representative images of reconstructed cells color coded by scaled expression of *Creb3l1* (female, top) or *Pou6f2* (male, bottom) in the MPOA at ages P23 (left) and P50 (right). Violin plots show the number of *Creb3l1* (female, top) or *Pou6f2* (male, bottom) transcripts per cell at ages P23 and P50. **d, e,** Motif-enrichment analysis of Esr1-DEG deconstructed GRNs in females (**d**) and males (**e**). The visual representation of the GRNs show *Esr1* and Esr1-regulated TFs cis-regulate Esr1-DEGs. TFs are depicted by numbered boxes and circles denote Esr1-DEGs color coded by ontology category. **f, g,** Violin plots comparing Esr1KO to control Vgat+ cell gene expression (measured as number of transcripts per cell) in MPOA of females (**f**) and males (**g**). Violin plots are outlined with a distribution line, individual dots represent each cell, and depict a red dot indicating the median. Violin plots are analyzed with Wilcoxon rank-sum test. ***p < 0.001, **p < 0.01. *p < 0.05. Statistical details in Methods. GRN: gene regulatory network; pTF: primary transcription factor; HA-DEG: hormone-associated differentially expressed gene; GDX: gonadectomy; OVX: ovariectomy; ORX: orchiectomy.

To validate these Esr1-GRNs *in situ*, we virally deleted *Esr1* from MPOA neurons in female and male mice prior to the onset of adolescence and conducted HM-HCR FISH on brain sections at P50, comparing Esr1KO to Esr1-intact controls. The gene panel included genes identified via SCENIC (Details in Methods)–3 Esr1-TFs: *Zmat4* (sex-shared), *Pou6f2* (male-specific), and *Creb3l1* (female-specific), and 6 Esr1-DEGs regulated by *Esr1* or Esr1-TFs: *Gria3*, *Pdzrn4*, *Ctnna2*, *Nefl*, *Tead1*, and *Esr1*. Consistent with our scRNAseq analysis (**Fig. 8b-e**), sex-shared Esr1-TF, *Zmat4*, was reduced by Esr1KO in both sexes, while male-specific Esr1-TF, *Pou6f2*, was decreased only in males. Female-specific Esr1-TF, *Creb3l1*, was reduced in females, but also in males to a lesser degree, perhaps because of high sensitivity in HM-HCR FISH assays. The expression of sex-shared Esr1-DEGs: *Gria3*, *Pdzrn4*, and *Ctnna2*, were decreased in Esr1KO females and males, and female-specific Esr1-DEG, *Tead1*, was reduced only in females (**Fig. 8f-g**). These results further suggest that *Esr1* orchestrates sex-shared and -specific transcriptional states via an *Esr1*-specific gene regulatory network. These dynamic gene regulatory networks may in turn dictate the sex-shared and -specific adolescent changes that facilitate the maturation of sexually dimorphic neuronal circuits for mating.

## Discussion

Adolescence represents a critical period for the maturation of behaviors and brain functions necessary for reproduction and social interactions. Unlike homeostatic processes such as eating and breathing, many behaviors that define adulthood emerge through hormonally driven neural circuit refinements during adolescence ^1,3,4^. Our study provides a detailed analysis of transcriptional dynamics in MPOA Esr1+ neurons, revealing their essential role in the adolescent maturation of mating behaviors in both sexes. Using a combination of single-cell RNA sequencing, advanced *in situ* hybridization techniques, and functional genomic manipulations, we show that Esr1 is a central regulator of these adolescent transcriptional changes. We show that MPOA Esr1+ neurons are transcriptionally dynamic during adolescence, progressing from an immature state in preadolescence to mature adult-like states in a hormone-dependent manner. This maturation process was arrested in mice lacking *Esr1* in MPOA Vgat+ neurons during adolescence, leading to profound deficits in mating behaviors. Importantly, these findings highlight that transcriptional programs driving sexual maturation are cell-type specific and temporally constrained, dependent on precise hormonal signaling during adolescence. Adolescent transcriptional dynamics in MPOA Esr1+ neurons appear to be continuous, with trajectories shaped by circulating sex hormones. This hormone-dependent maturation was confirmed by pseudotime analysis, which revealed that hormonal deprivation arrests transcriptional progression, while hormone supplementation accelerates it. These findings align with the broader view that the adolescent brain undergoes extensive molecular and circuit-level adaptations to support the emergence of sexually dimorphic behaviors ^3,16,44^.

## Integration of MPOA sexual dimorphism and adolescence-related transcriptional dynamics

One of the major contributions of this study is the disentanglement of sexual dimorphism and adolescent dynamics in Esr1+ neurons. We identified subsets of genes that are both sexually dimorphic and adolescently dynamic, as well as genes that are unique to each process. This complex interplay underscores the importance of analyzing these processes separately in a sex-specific fashion to uncover their contributions to neural development. Our findings suggest that Esr1 not only regulates common gene programs shared by males and females but can also orchestrate sexually distinct transcriptional networks. The male-specific enrichment of androgen receptor-related genes and the identification of sexually dimorphic Esr1-dependent transcription factors (e.g., *Pou6f2* in males, *Creb3l1* in females) further highlight how Esr1 integrates hormonal signals to shape sex-specific neural circuits ^42,43^.

## Functional implications for neural circuitry and behavior

The adolescent transcriptional changes observed in MPOA Esr1+ neurons likely translate to functional adaptations in mating-relevant neural circuits. Previous studies have shown that MPOA neurons are critical for a wide range of sexually dimorphic behaviors, including mounting, vocalization, and parental care ^12,16,20,45^. Our study provides molecular evidence linking the transcriptional maturation of MPOA Esr1+ neurons to these behaviors. This link is especially compelling considering our integration with published MERFISH datasets, which further validated the association of Vgat+Esr1+ neurons with mating behaviors in both sexes ^18,22^. Dynamic changes in the circuit properties of MPOA neurons during adolescence may involve alterations in cellular excitability, synaptic connectivity, and interactions with downstream targets. Ontology analysis of Esr1-dependent genes revealed enrichment for pathways related to synaptic transmission and axonal structure, supporting the hypothesis that transcriptional maturation directly contributes to circuit remodeling ^29,46^. Future studies combining transcriptional profiling and CRISPR gene editing with *in vivo* imaging and circuit manipulation will be essential to dissect these connections. Our study complements previous work on perinatal transcriptional dynamics in sexually dimorphic brain regions, such as the BNST ^23^. While the perinatal period is critical for establishing primary sexual differentiation, our findings emphasize adolescence as a secondary critical period where sexually dimorphic transcriptional programs are refined. Notably, both periods rely heavily on Esr1-mediated regulation, suggesting a conserved role for estrogen signaling across developmental stages.

## Limitations of the study

While these results suggest that male and female adolescent transcriptional maturation has unique and overlapping components, some technical limitations of the results are worth noting. First, UMIs per cell are low due to older scRNAseq technology being used. We used HM-HCR FISH to corroborate our findings because this technique is very sensitive for detecting targeted genes. However, it is likely that the transcriptional state changes we observed are blunted due to sparse gene expression detection in 10X V2 kits compared to now-standard versions of these reagents. Additionally, while KO of Esr1 has dramatic impacts on mating behavior in both sexes, due to the massive number of genes that are regulated by Esr1, this is not surprising. Now that we have a list of targeted genes that are likely directly under the influence of Esr1, future studies are needed to resolve how individual or groups of Esr1-associated DEGs are critical for specific aspects of mating behavior.

## Technical limitations of the study

1. Although HM-HCR experiments showed the bidirectional control of transcriptional progression during adolescence, it is unclear if the facilitation in male by testosterone supplement is via activation of AR or Esr1 or both because testosterone will likely be converted to estrogen in the brain. Future studies using dihydrotestosterone (DHT) and estrogen to males may address this issue.
2. Although we have identified hormone/Esr1 dependent transcriptional trajectories during adolescence, the relations and interplay with genetically determined perinatal event, which is earlier and robust, are unclear. Some sex differences during adolescence might be an extension of perinatally established sex differences while others might be unique adolescent changes.
3. While we have observed robust effect of Esr1-KO in scRNAseq experiment which was further validated with FISH experiment, it is possible that there are further heterogeneous Vgat-Esr1 populations in the MPOA which might be differentially targeted in each virally injected sample. To mitigate this, 3-4 mice were pooled for each sample in scRNAseq experiment and in HCR-FISH experiment, in addition to confirming recombinase RNA expression within the MPOA, we included samples with robust Esr1 deletion in the MPOA. Interestingly, due to the technical challenge, Esr1 deletion tends to be more robust than weakly detected recombinase RNA expression (data not shown).
4. While we have observed robust transcriptional progression in Vgat^+^ Esr1^+^ neurons during adolescence, we observed more mild alternations in VgluT2^+^ neurons. Although the scale of our study is comparable or exceeds prior scRNAseq studies in MPOA^22,29^, future larger studies may have more sensitivity to detect adolescent transcriptional dynamics in VgluT2^+^ neurons.
5. Although we demonstrated adolescent transcriptional changes were observed as early as P35, and either hormonal deprivation or Esr1 KO in prior to adolescence prevented the transcriptional progression (arrested transcriptional state even at adult), given the viral incubation time and permanent deletion of Esr1 after viral injection, it is challenging to disambiguate the role of Esr1 during adolescence and adult. Future studies injecting the virus at adult may provide additional insights on the similarity and difference between transcriptional changes during puberty and maintained transcriptional states at adult.

## Conclusions and future directions

These findings have implications beyond mating behaviors. The role of Esr1+ neurons in the MPOA likely extends to other hormonally sensitive behaviors and physiological processes, such as thermoregulation, metabolic control, and social bonding ^17,21,47^. Additionally, understanding how disruptions in these transcriptional programs contribute to developmental disorders—such as delayed puberty, hypogonadism, or neuropsychiatric conditions with sex-biased prevalence—represents an important avenue for future research ^1,2^. While our study focused on Esr1, it is likely that other transcription factors and hormone receptors synergistically contribute to adolescent transcriptional dynamics. For example, the androgen receptor may play a role in males, while additional regulatory networks may be governed by sex chromosome-linked genes ^43^. Advanced genomic approaches, such as perturb-seq, could help unravel the complexity of these gene regulatory networks in a cell-type-specific manner ^31^. Although we focused on adolescent transcriptional trajectories of Vgat+Esr1+ neurons, this dataset provides a rich resource for studying transcriptional states across other neuronal, glial, and stromal MPOA cell types during adolescent brain development. Understanding steroid-induced changes in transcriptional trajectories during two major developmental periods (perinatal and adolescence periods) has implications for the healthy and maladaptive development of human brains, and provides additional insight into neurobiological mechanisms that underlie sex differences ^48–50^.

## Acknowledgements

We thank M. Borrego for critical comments on the manuscript. We thank C. Trapnell for helpful discussions. We thank Y. Tao and Z. Hu (University of North Carolina) for assistance with initial 10x Genomics library generation.

## Author contributions

Conceptualization: GDS, KH, YH, KI, BB, JM, DR Methodology: KH, YH, KI, BB, MLB, RDP, LSZ

Investigation: KH, YH, KI, BB, YL, MAR, JYC, ORA, RVM, NLJ, RCS, JCS, JRS,

Data analysis: KH, YH, KI, BB, GDS, JCS, JRS Manuscript writing: GDS, KH, YH, BB, KI Manuscript editing: all authors

Supervision: GDS

## Data and materials availability

The NCBI Gene Expression Omnibus accession number for the scRNAseq data reported in this paper for the reviewers is GEO: GSE172177 All the codes used to analyze scRNAseq, and HM-HCR FISH are available at a Github repository affiliated with Stuber Laboratory group and this manuscript title (http://www.github.com/stuberlab/).

## Abbreviation

medial preoptic area (MPOA)

single-cell RNA sequencing (scRNAseq) estrogen receptor 1 (Esr1)

GABAergic neurons (Vgat+) glutamatergic neurons (Vglut2+) hybridized chain reaction fluorescent *in situ* hybridization (HCR-FISH) gonadectomized (GDX) partition-based graph abstraction (PAGA) hormone-associated differentially expressed genes (HA-DEGs) multiplexed error-robust fluorescence *in situ* hybridization (MERFISH) differential gene expression (DE) differentially expressed genes (DEGs) support vector machine (SVM) Manifold Enhancement Latent Dimension (MELD) Potential of Heat-diffusion for Affinity-based Trajectory Embedding (PHATE) androgen receptor (AR) single-cell regulatory network inference (SCENIC)

## Materials and Methods

### Mice

Several lines of male and female mice were used for scRNAseq, HM-HCR FISH, immunohistochemistry (IHC), and behavioral experiments, with specific ages and experimental conditions described in their respective sections. All mice used in these experiments were on a C57BL/6 background. Wild-type C57BL/6J mice were originally obtained from Jackson Laboratory (JAX) and bred in-house. Transgenic *Slc32a1*^Flp^::*Esr1*^lox/lox^ mice were generated by crossing *Slc32a1*^Flp^ mice (Jackson Laboratory stock no. 029591) with *Esr1*^lox/lox^ mice (kindly provided by Kenneth Korach; Jackson Laboratory stock no. 032173), and then breeding the resulting progeny with *Slc32a1*^Flp^::*Esr1*^lox/+^ mice. Similarly, *Slc17a6*^Flp^::*Esr1*^lox/lox^ mice were generated by crossing *Slc17a6*^Flp^ mice (Jackson Laboratory stock no. 030212) with *Esr1*^lox/lox^ mice and then crossing the progeny with *Slc17a6*^Flp^::*Esr1*^lox/+^ mice. For both lines, only mice heterozygous for Flp and homozygous for Esr1^lox/lox^ were used in experiments. For scRNAseq and HCR experiments, mice were group-housed except for the 1–2 days prior to tissue isolation, during which they were singly housed. Behavioral experiment mice were singly housed for the duration of the experiments. Wild-type Swiss Webster (SW) mice, originally obtained from Taconic and bred in-house, were used as lactating foster dams to rear pups following brain surgery, as C57BL/6 dams frequently committed infanticide after such procedures. All mice were maintained with *ad libitum* access to food and water under a reverse 12-hour light-dark cycle. All experiments were conducted in accordance with the National Institutes of Health’s *Guide for the Care and Use of Laboratory Animals* and approved by the Institutional Animal Care and Use Committees (IACUC) of the University of North Carolina and University of Washington.

## Stereotaxic viral injection

Juvenile mice aged P14–18 were separated from their dams, anesthetized with isoflurane, and positioned in a stereotaxic frame (Kopf Instruments). Isoflurane anesthesia was maintained at less than 0.8% throughout the procedure. Following a craniotomy above the target region, viral injections were performed using glass capillaries at a controlled rate of 60 nl/min with a Nanoject system (Drummond). To ensure accurate delivery and minimize backflow, injections were paused for 30 seconds after every 60 nl infusion, and the capillaries were left in place for 10 minutes before withdrawal. The stereotaxic coordinates used were 0.2 mm AP and 0.23 mm ML from Bregma; -4.95 mm DV from the brain surface. After recovery from anesthesia, mice were transferred to the care of lactating Swiss Webster (SW) foster dams. SW dams effectively nursed surgically treated pups, whereas C57BL/6 dams frequently committed infanticide following such procedures.

## Gonadectomy

Gonadectomized mice were prepared for scRNAseq and HCR experiments. At P22–23, mice were anesthetized with isoflurane and maintained under anesthesia at less than 1.0%. After shaving the hair around the flank area for females or the scrotum for males, local anesthesia was administered, and a small incision was made through the skin and muscle above the target gonads. The gonads were identified, gently exteriorized with forceps, and removed using a heat cautery pen (Bovie Medical). The remaining tissues were returned to the abdominal cavity (females) or scrotum (males). The incision site was closed with sutures, and tissue adhesive (Vetbond) was applied to ensure secure skin closure. Brain tissues were collected at P50 ± 2, ensuring the mice were deprived of sex hormones throughout adolescence. Gonadectomized adult female mice were also prepared as stimulus animals for Resident Intruder (RI) and three-chamber sociability assays. For these experiments, 7–8-week-old C57BL/6J female mice were ovariectomized using the same procedure. After a three-week recovery period, a standard hormonal priming protocol was used to induce estrus. This involved subcutaneous injections of β-Estradiol 3-Benzoate (Sigma, 10 µg at 48 hours and 5 µg at 24 hours before the assay) and Progesterone (Sigma, 500 µg) administered six hours prior to the RI assay.

## Generation of AAV-frtFlex-Cre virus

The split-Cre design used in this study was previously described ^51,52^. In this system, an intron is introduced into the middle of the Cre sequence, with the second exon flanked by frt sites. The action of FLP recombinase inverts the second exon, enabling splicing to generate functional Cre. For these experiments, the construct was incorporated into an AAV vector. The frtFlex-Cre-EGFP cassette was subcloned into the pAAV-CAG-WPRE vector between AscI and XhoI restriction sites. AAV1 virus carrying frtFlex-Cre-EGFP was produced as described previously ^52^. Briefly, pAAV-CAG-frtFlex-Cre-EGFP-WPRE was transiently transfected into HEK293T/17 cells (ATCC) along with the pDG1 packaging plasmid. Twenty-four hours post-transfection, cells were transferred to serum-free media, and viral capsids were harvested 48 hours later. Purification of AAV1 capsids was performed using gradient centrifugation. Viral titer was estimated at approximately 2 x 10^12^ particles/mL based on densitometry analysis following gel electrophoresis, using a known standard for reference.

## Validation of efficiency and specificity of AAV-frtFlex-split-Cre virus

To confirm that AAV-frtFlex-Cre does not produce functional Cre in the absence of FLP recombinase, 300 nl of AAV-frtFlex-Cre was unilaterally injected into the MPOA of *Esr1*^lox/lox^ mice (n = 3 males and 3 females; data combined across sexes) (**Supplementary Fig. 1a**). To validate the specificity of AAV-frtFlex-Cre, 300 nl of the virus was unilaterally injected into the MPOA of *Slc32a1*^Flp^ and *Slc17a6*^Flp^ mice. Two weeks post-transduction, brains were collected and analyzed using FISH, as described in the RNAscope section (**Supplementary Fig. 1b**). Six weeks after the injection, brain tissue was collected and analyzed via IHC, as detailed in the Histology section. To assess the efficiency of AAV-frtFlex-Cre, 300 nl of the virus was bilaterally injected into the MPOA of *Slc32a1*^Flp^::*Esr1*^lox/lox^ and *Slc17a6*^Flp^::*Esr1*^lox/lox^ mice. These mice were first used for behavioral experiments, as described in the Behavioral Experiment section, and efficiency was quantified post-behavioral testing. Quantitative results for each mouse line and sex are presented in **Fig. 1 and Supplementary Fig. 1**.

## Single-cell preparation and cDNA library construction for scRNAseq

Male and female wild-type mice (n = 3–4 animals pooled per group) were prepared at P23 ± 1 (preadolescence, hormonally intact), P35 ± 1 (mid-adolescence, hormonally intact), and P50 ± 2 (hormonally intact early adulthood or gonadectomy groups). For gonadectomy groups, mice were gonadectomized at P22–23. Additionally, male and female *Slc32a1*^Flp^::*Esr1*^lox/lox^ mice (n = 3–4 per group), which had received bilateral MPOA injections of 300 nl AAV-frtFlex-Cre at P14–18, were collected at P50 ± 2 (early adulthood). All mice were singly housed 1–2 days before tissue dissection.

Single-cell preparation followed previously published procedures ^53^. Immediately after removal from their home cages, mice were deeply anesthetized with an intraperitoneal injection of 0.2 mL sodium pentobarbital (39 mg/mL) and phenytoin sodium (5 mg/mL), followed by transcardial perfusion with ice-cold NMDG-aCSF containing transcription and translation inhibitors to minimize transcriptional events induced by anesthesia, perfusion, or brain extraction. NMDG-aCSF consisted of 96 mM NMDG, 2.5 mM KCl, 1.35 mM NaH2PO4, 30 mM NaHCO3, 20 mM HEPES, 25 mM glucose, 2 mM thiourea, 5 mM Na+ ascorbate, 3 mM Na+ pyruvate, 0.6 mM glutathione-ethyl-ester, 2 mM N-acetyl-cysteine, 0.5 mM CaCl2, and 10 mM MgSO4 (pH 7.35–7.40, 300–305 mOsm, oxygenated with 95% O2 and 5% CO2). Inhibitor cocktails included 500 nM TTX, 10 μM APV, 10 μM DNQX, 5 μM actinomycin, and 37.7 μM anisomycin. Brains were extracted, and coronal sections containing the MPOA were sliced at 300 µm using a vibratome (Leica VT1200). Slices (3–4 per animal) were recovered on ice in a chamber for 30 minutes.

MPOA tissues were dissected under a light microscope (Leica MZFL) using a micro scalpel (Feather) from 3–4 animals per group (10–15 slices pooled per group). The tissue was enzymatically digested with 1 mg/mL pronase (Sigma-Aldrich) for 30 minutes at room temperature, followed by mechanical trituration using fire-polished glass capillaries (tip diameter 200–300 µm). Cell suspensions were filtered twice through 40 µm strainers to remove aggregates, and dead cells were eliminated using a dead cell removal kit (Miltenyi Biotec). After centrifugation, the supernatant was removed, and cells were resuspended (20–30 mL). A fraction (∼5 mL) was mixed with trypan blue for viability assessment and cell concentration measurement using a hemocytometer. Samples with >80% viability were used for cDNA library preparation, and final cell concentrations were adjusted to 800–1,000 cells/µL. Typically, 30,000–80,000 cells were collected from 3–4 mice per group, exceeding the number required for cDNA library construction and reducing biological variability between subjects.

cDNA libraries were prepared using the Chromium Single Cell 3’ Reagent Kits V2 according to the manufacturer’s instructions (10x Genomics). Approximately 17,000 dissociated cells were mixed with reverse transcription mix and loaded onto a chip to recover up to 10,000 single cells. mRNAs from single cells were captured by barcoded beads in droplets using a Chromium Controller. Reverse-transcribed cDNAs were PCR amplified, fragmented, and ligated with adapters, followed by sample indexing via PCR. cDNA libraries were sequenced on an Illumina NextSeq 500 (v2.5) or HiSeq system. Sequencing reads were aligned to the mouse genome using the 10x Genomics Cell Ranger pipeline (V3) to generate cell-by-gene count matrices for downstream analysis.

## Highly Multiplexed HCR FISH

HM-HCR FISH assays were conducted to validate trajectory inference from scRNAseq data (HCR1; **Fig. 4 and Supplementary Fig. 8–9**) and to cross-validate the Esr1-GRN analysis (HCR2; **Fig. 8**).

For HCR1, we analyzed groups included in the scRNAseq experiments, including preadolescence (P23 ± 1), adolescence (P35 ± 1), intact adult (P50 ± 2), and gonadectomy groups (P50 ± 2, gonadectomy performed at P22–23). Additionally, a hormone-supplemented group received daily injections of sex hormones (females: 5 µg β-Estradiol 3-Benzoate in 50 µl sesame oil; males: 200 µg Testosterone Propionate in 50 µl sesame oil) from P22 to P27, with tissue collected at P28. A control group for hormone supplementation received daily injections of sesame oil (50 µl) during the same period, and brains were harvested at P28. The timing of hormone supplementation aligned with the onset of adolescence in peripheral tissues (P28–33; **Fig. 1c, f**) and reports from the pituitary (P25–30; ^54^). Adolescence onset was verified in hormone-supplemented animals by assessing male balanopreputial separation (BPS) and female vaginal opening (VO). All hormone-supplemented animals displayed BPS or VO by P28, whereas none of the controls reached adolescence by this age. All mice in the 12 groups were singly housed 1–2 days prior to tissue collection.

For HCR2, AAV-Cre-YFP or AAV-Flp-YFP was unilaterally injected into the MPOA of Esr1^lox/lox^ mice of both sexes at P30–35, and brain tissues were harvested three weeks later. For both HCR1 and HCR2, mice (n = 2 per group) were deeply anesthetized with isoflurane, and brains were rapidly extracted and frozen on dry ice. Coronal sections (20 µm) were prepared using a cryostat (Leica) and stored at - 80°C until use.

Sequential HCR FISH followed a modified protocol based on ^40^. Tissue sections were fixed in pre-chilled 4% paraformaldehyde (PFA) for 30 minutes on ice, rinsed twice in 1x PBS at room temperature, dehydrated in ethanol (50%, 70%, 100%, 100%; 5 minutes each), and air-dried for 5 minutes. Sections were then permeabilized with protease IV (ACD, 322336) for 5 minutes at room temperature, followed by rinses in 1x PBS and 2x SSC. Probes, amplifiers, and buffers (Molecular Instruments) were prepared as per the manufacturer’s guidelines (Choi et al., 2018). Tissue was equilibrated in probe hybridization buffer for 10 minutes at 37°C in a humidified chamber, then incubated overnight with probe mixtures (final oncentration 4–10 nM) at 37°C. Coverslips were removed in 100% pre-warmed wash buffer at 37°C, and tissues were sequentially washed in dilutions of wash buffer in 5x SSCT (75%, 50%, 25%; 15 minutes each), then in 5x SSCT at 37°C (15 minutes) and at room temperature (5 minutes).

For signal amplification, tissues were equilibrated in amplification buffer for 30 minutes at room temperature, then incubated overnight with snap-cooled hairpins conjugated to Alexa488, 546/594, and 647 (final concentration 60 nM). Coverslips were removed in 5x SSCT at room temperature, followed by two 30-minute washes in fresh 5x SSCT and rinses in 2x SSC. Autofluorescence was minimized with a quenching kit (Vector Laboratories, SP-8400-15). Sections were treated with quenching solution for 2 minutes, rinsed in 2x SSC, counterstained with DAPI (ACD, 320858) for 30 seconds, mounted in Vectashield (Vector Laboratories, H-1700-10), and stored at 4°C. DAPI staining was performed only in the first round, and all images were acquired within 24 hours.

Probes and amplifiers were stripped between rounds to enable multiplexing. Coverslips were floated off in 2x SSC for 30 minutes at room temperature, and tissue was incubated in DNase I (250 U/mL in 1x DNase I buffer, Roche, 04716728001) for 1.5 hours at room temperature, followed by six washes in 2x SSC (5 minutes each). Tissue was then equilibrated in pre-hybridization buffer for subsequent rounds. This process was repeated for up to four rounds, allowing detection of up to 12 different mRNAs.

Images were acquired using an Axio Imager M2 fluorescence microscope (Zeiss) equipped with Zen software. Channels for green (Alexa488), red (Alexa546/594), and far-red (Alexa647) fluorescence were captured, along with brightfield images to aid registration. DAPI signals from the first round defined nuclear regions of interest (ROIs), which were expanded to include cytoplasmic transcripts. Image registration was performed using HiPlex software (ACD), with brightfield images from subsequent rounds aligned to the first round using transformation matrices. Up to 17 images were overlaid, and overlapping regions were cropped to generate a single 12-plex image.

For quantification, HCR FISH images were analyzed in ImageJ. DAPI-defined ROIs were transferred to the 12-plex image to measure transcript numbers for each gene. For HCR1, genes enriched in Vgat+Esr1+ neurons and HA-DEGs from scRNAseq data were analyzed, with validation by SVM (**Supplementary Fig. 6k, m**). For HCR2, Esr1-TFs and Esr1-DEGs regulated by Esr1-TFs or Esr1 were selected from the Esr1-GRN (**Fig. 8**). To disambiguate the MPOA and adjacent brain regions, quantitative analysis is restricted to Vgat+Esr1+ neurons and is devoid of posterior BNST. For HCR2, AAV was injected unilaterally so that successful targeting of the MPOA with AAV-Cre-YFP (detection of recombinase RNA within the MPOA) and the deletion of Esr1 were confirmed for inclusion of samples.

## RNAscope

AAV-frtFlex-Cre (300 nl) was unilaterally injected into the MPOA of VgatFlp or Vglut2Flp mice (n = 2 per experiment). After two weeks of viral incubation, mice were deeply anesthetized with isoflurane, and brains were rapidly extracted and frozen on dry ice. Coronal sections (20 µm) were prepared using a cryostat (Leica) and stored at -80°C until further use. Probe hybridization and signal detection were performed according to the manufacturer’s instructions (ACDbio). In VgatFlp mice, probes targeted *Slc32a1* and *eGFP*, while in Vglut2Flp mice, probes targeted *Slc17a6* and *eGFP*. Tiled images of the MPOA were acquired using a Zeiss ApoTome2 fluorescence microscope with a 20x objective and Zen software (Zeiss). Image acquisition settings were consistent across all experiments. The resulting CZI files were analyzed using Fiji. The number of cells expressing *Slc32a1* or *Slc17a6* and *eGFP*, as well as double-positive cells expressing both markers, were quantified for each mouse line (**Fig.1 and Supplementary Fig. 1**).

## Behavioral experiments

Male and female *Slc32a1*^Flp^*::Esr1*^lox/lox^ or *Slc17a6*^Flp^*::Esr1*^lox/lox^ mice were bilaterally injected with 300 nl of AAV-frtFlex-Cre (Cre group; virus generated in-house, detailed in the Generation of AAV-frtFlex-Cre virus section) or AAV-fDIO-eYFP (control group; UNC Vector Core) at P14–18. Starting at P25, sexual organ development was inspected daily to determine the age of first balano-preputial separation (BPS) in males or vaginal opening (VO) in females. For female subjects, vaginal smears were also collected daily following vaginal opening to monitor estrous cycles. Behavioral experiments, including the Resident Intruder (RI) assay, locomotion tests, sociability assays, and the Elevated Plus Maze (EPM), were conducted during the second half of the dark cycle under red-light illumination. Body weights were recorded daily from P30 to P54. Upon completion of the behavioral experiments, subjects were deeply anesthetized, and brain tissues were collected for histological analysis (detailed in the Histology section). Mice that failed to gain weight or received mistargeted viral injections were excluded from the analysis.

To assess the adolescent maturation of sexual behaviors, male subjects underwent RI assays from P34 to P54 every other day. In this assay, a hormonally primed adult female mouse was introduced into the male subject’s home cage for a 15-minute interaction (Cre groups: *Slc32a1*^Flp^*::Esr1*^lox/lox^, n = 12; *Slc17a6*^Flp^*::Esr1*^lox/lox^, n = 9; control groups: *Slc32a1*^Flp^*::Esr1*^lox/lox^, n = 13; *Slc17a6*^Flp^*::Esr1*^lox/lox^, n = 9). To prevent ejaculation, male subjects were separated from the female intruder as soon as thrusting began. The interactions were video-recorded, and the number of mounting and thrusting behaviors was manually counted.

Female sexual receptivity was assessed in RI assays conducted from P35 to P54 for *Slc32a1*^Flp^*::Esr1*^lox/lox^ mice (Cre group: n = 16; control group: n = 10) and from P40 to P60 for *Slc17a6*^Flp^*::Esr1*^lox/lox^ mice (Cre group: n = 9; control group: n = 11). Assay timing was guided by pilot experiments and conducted only when vaginal smears indicated the female was in proestrus or estrus. Female subjects were introduced into the home cage of a sexually experienced adult male mouse and allowed to interact freely for 10 minutes. To avoid ejaculation, females were separated from the male after intromission began or approximately 5 seconds after an unsuccessful mounting attempt. These interactions were recorded using an IR camera controlled by Ethovision (Noldus), and behaviors such as being mounted, intromitted, escaping, or displaying submissive postures were manually counted. Receptivity was calculated as the proportion of intromission bouts out of total mounting attempts. Stress levels were minimized and equalized across groups by selecting highly sexually experienced but minimally aggressive male residents. In rare instances of aggression, male residents were promptly removed. Unlike mice with brain-wide *Esr1* deletions ^27^, the male subjects in this study, with *Esr1* knockouts limited to MPOA Vgat+ neurons during adolescence, showed no aggressive behaviors, consistent with previous findings ^15^.

Sociability tests were conducted after P45 on days when RI assays were not performed. Subjects were placed in a standard three-chamber choice arena, where one side contained a caged social stimulus and the other an object. Stimulus mice were either adult males or hormonally primed females. After a 5-minute habituation period without stimuli, a social stimulus and an object were introduced, and the subject mouse was allowed to explore for 10 minutes. The stimuli were then exchanged for a new mouse of the opposite sex and a novel object, and the subject mouse explored for an additional 10 minutes. Movements and locations were tracked using an IR camera controlled by Ethovision (Noldus). Total distance traveled during the habituation period was used to assess locomotion, while the time spent in the chamber with the social stimulus was used for statistical comparisons.

The Elevated Plus Maze (EPM) test was performed after P45 on days when neither RI assays nor sociability tests were conducted. Subjects were placed in a standard EPM arena and allowed to explore freely. After a 5-minute habituation period, movements were tracked using an IR camera controlled by Ethovision (Noldus). Time spent in open and closed arms of the maze was recorded for statistical analysis.

## Histology

Histological experiments were conducted to: (1) examine viral infection in the MPOA of mice used in behavioral experiments (*Slc32a1*^Flp^*::Esr1*^lox/lox^ or *Slc17a6*^Flp^*::Esr1*^lox/lox^), (2) validate the specificity of AAV-frtFlex-Cre in *Esr1*^lox/lox^ mice, and (3) confirm the specificity of reporter gene expression in *Slc32a1*^Flp^*::Esr1-Cre::RC-FLTG* mice. Mice were deeply anesthetized with pentobarbital and transcardially perfused with 40 mL of 4% paraformaldehyde (PFA) in PBS. Brains were extracted and post-fixed in 4% PFA in PBS on ice for 5–7 hours before being transferred to PBS with 0.05% sodium azide (Sigma) for storage at 4°C until sectioning. Free-floating coronal brain sections (50 µm thick) were prepared using a vibratome (Leica) and stored in PBS with 0.05% sodium azide until immunohistochemistry (IHC). Sections were first washed in PBS (3 × 5 minutes) and blocked with 15% normal donkey serum (NDS) in PBST (0.3% Triton X-100) for 2 hours at room temperature. Following blocking, sections were incubated with an anti-Esr1 primary antibody (1:500, Santa Cruz, sc-542) diluted in 15% NDS in PBST for 72 hours at 4°C. After incubation, sections were washed in PBST (0.3% Triton X-100, 3 × 30 minutes) and incubated with a secondary antibody (1:500, Life Technologies donkey anti-rabbit 568 or Jackson ImmunoResearch donkey anti-rabbit 647) diluted in 15% NDS in PBST for 2 hours at room temperature. Sections were then washed in PBST (2 × 15 minutes), incubated with DAPI for 2 minutes, rinsed in PBS (2 × 15 minutes), mounted on slides, and coverslipped with mounting medium. Tiled images of the MPOA were acquired using a Zeiss ApoTome2 fluorescence microscope with a 20x objective and Zen software (Zeiss). Image acquisition settings were consistent across all experiments. The resulting CZI files were analyzed with HALO software using the ISH-IF version 1.2 module (Indica Labs). Regions of interest (ROIs) corresponding to the MPOA were manually outlined in both hemispheres across three adjacent sections from each brain. GFP- or YFP-positive cells, Esr1-positive cells, and double-positive cells were automatically detected and quantified using specific threshold settings to define phenotypes.

## Data analysis for scRNAseq, MERFISH, and HM-HCR data

Analysis of scRNAseq, MERFISH, and HM-HCR FISH data utilized several R and Python packages, which were adapted and modified as needed. Previously published MERFISH data, deposited in DRYAD, was acquired from https://datadryad.org/stash/dataset/doi:10.5061/dryad.8t8s248. The data analysis workflow encompassed clustering, differential gene expression analysis, integration of differential conditions and modalities, lineage inference, gene set activity measurements, and gene regulatory network (GRN) deconstruction.

Clustering and differential gene expression analyses were performed using the Seurat V3 package ^32^, while LISI ^55^ was applied to evaluate integration performance across conditions. Lineage inference, representing transcriptional connectivity of neuronal clusters, was conducted with PAGA (partition-based graph abstraction; Wolf et al., 2019). Single-cell gene set activity was quantified using AUCell ^31^. GRNs were inferred and deconstructed using the SCENIC package, which integrates GENIE3 ^31,56^ and RcisTarget. These analyses depicted cis-regulatory relationships between transcription factors and differentially expressed genes. Gene co-expression networks were identified using scWGCNA ^57^. To examine transcriptional state progressions by age and hormonal state, Monocle V3 ^34^ and MELD ^38^ were applied. Ontology analysis was conducted using the online Enrichr platform ^58,59^.

The data analysis workflow consisted of four major steps: (1) clustering and identifying cell types that exhibit transcriptional dynamics during adolescence and are relevant for reproductive behaviors; (2) trajectory analysis to quantify adolescent transcriptional progression; (3) transcriptional analysis in spatial contexts; and (4) GRN deconstruction of adolescent transcriptional dynamics. Each step is detailed in subsequent sections.

## Data preprocessing and doublet removal for scRNAseq data

Preprocessing followed the procedures described in our previously published study ^53^. Low-expression genes, defined as those detected in fewer than three cells, and low-quality cells (total UMI < 700, total UMI > 15,000, or >20% mitochondrial gene expression) were excluded from downstream analysis. Suspected doublets were computationally removed using the DoubletDecon package (Version 1.1.5; ^60^) with default settings, applying a doublet rate of 5.6 ± 1.0%.

## Integrative clustering and differential gene expression analysis

To minimize the effects of batch variability and differences due to sex, age, and hormonal state while preserving the global similarity of transcriptional states within cell types, we applied the Seurat V3 integrative approach ^32^. This method combines canonical correlation analysis (CCA; ^61^) and mutual nearest neighbor analysis ^62^. After preprocessing the data, which included 58,921 cells in total, gene counts were normalized using the NormalizeData function to scale counts by total UMI with a constant scale factor (10,000), followed by natural-log transformation (log1p). The FindVariableFeatures function was used to select 2,000 highly variable genes from each sample based on variance stabilizing transformation. Integration was performed pairwise using the FindIntegrationAnchors function (CCA1-40) to identify anchors and assign scores, followed by the IntegrateData function to compute an integrated gene expression matrix by iteratively constructing and subtracting transformation matrices from the original data.

The integrated expression data was used for clustering. Expression matrices were scaled, centered, and reduced using principal component analysis (PCA). A nearest-neighbor graph was constructed in PCA space (FindNeighbors; default settings), followed by Louvain clustering at a resolution of 0.8 (FindClusters). Uniform Manifold Approximation and Projection (UMAP) was then generated for visualization (RunUMAP). Initial clustering identified 32 clusters, 13 of which were neuronal, expressing canonical markers such as *Thy1* or *Stmn2*. Cell types were determined based on marker expression (Neuron: *Stmn2, Thy1;* Astrocyte: *Ntsr2;* OPC: *Gpr17;* Oligodendrocyte: *Mog;* Microglia: *C1qc;* Mural cell: *Tagln;* Endothelial cell: *Flt1;* Intermediate cell: *B2m;* Ependymal cell: *Foxj1*) as reported in previous studies ^53,63,64^. Neuronal and non-neuronal cell types were combined for visualization (**Supplementary Fig. 2a-d**).

To identify conserved markers, we used the FindConservedMarkers function to compare gene expression in each cluster against the rest of the dataset using the Wilcoxon rank-sum test. P-values were adjusted for the number of genes tested, and genes with an adjusted p-value < 0.05 were considered significantly enriched. Clustering robustness was assessed by sub-sampling 10–100% of cells and re-clustering using the same pipeline, repeated 10 times at each sampling rate.

For higher-resolution analysis, the 24,831 cells in the 13 neuronal clusters were extracted and re-clustered using the same integrative clustering approach. This resulted in 36 clusters, of which one (204 cells) was excluded due to low expression of neuronal markers. The remaining 24,627 cells in 35 neuronal clusters were analyzed using the FindConservedMarkers function to identify cluster-specific marker genes. These neuronal clusters included cells from all groups across different ages, sexes, and hormonal states.

Differentially expressed genes (DEGs) between groups were calculated using the FindMarkers function. Pairwise comparisons were performed within aggregate Vgat+ or Vglut2+ clusters for each sex (criteria: >10% expression, logFC > 0, adjusted p-value < 0.05). Hierarchical clustering of DEGs generated dendrogram trees, with a threshold (h = 3.15) used to identify DEG clusters. This analysis revealed DEG clusters with higher expression in P50 and P35 groups compared to P23 and gonadectomy (GDX) groups in both Vgat+ and Vglut2+ cells. Additional analyses identified hormonally associated DEGs (HA-DEGs) by comparing gene expression between P50 or P35 and GDX groups in individual or aggregate clusters (e.g., Vgat+Esr1+, Vgat+Esr1-Ar+, Vgat+ hormone RLow; where + indicates >50% positive cells and - indicates <10% positive cells).

To assess integration performance, Local Inverse Simpson’s Index (LISI; ^55^) was computed in UMAP space, with LISI scores reflecting the effective number of distinct groups in the local neighborhood of each cell (**Supplementary Fig. 2i**). Clustering stability was further evaluated by examining lineage relationships between clusters at varying resolutions (0.2–2.0) using the clustree package ^65^. Clustering tree analysis showed the emergence of heterogeneous *Esr1* clusters at resolution 0.8 or higher (**Supplementary Fig. 2k**). Specific marker genes in neuronal clusters (**Supplementary Fig. 3a**) and Vgat+Esr1+ subclusters (**Supplementary Fig. 3h**) further validated clustering.

To explore transcriptional connectivity between neuronal clusters, PAGA ^30^ was applied. In the resulting PAGA graph, nodes represented clusters, and edge thickness corresponded to connectivity strength. Globally, the PAGA graph revealed distinct separation between Vgat+ and Vglut2+ clusters. Locally, Vgat+Esr1+ clusters (Vgat2, Vgat4, and Vgat16) exhibited high connectivity and were distinct from other clusters (**Fig. 2c**).

## Integrative analysis of MERFISH and scRNAseq data

Seurat V3 was used to jointly analyze publicly available POA MERFISH data ^22^ and our scRNAseq data. To establish correspondence between MERFISH and scRNAseq clusters related to reproductive behaviors in adult mice, cells from behaviorally naïve subjects (lacking *Fos* expression data), lactating females, or individuals exhibiting aggression toward pups were excluded. Cells categorized as “Inhibitory” or “Excitatory” in the MERFISH metadata were extracted and normalized using the NormalizeData function. Highly variable genes were identified with FindVariableFeatures, and the top 60 variable genes (excluding *Fos*) were used for clustering, following the procedures described earlier.

Seurat clustering identified 19 neuronal clusters within the MERFISH dataset, including 10 GABAergic clusters enriched with *Gad1* and 6 excitatory clusters expressing *Slc17a6*. Because *Fos* data from behaviorally naïve animals was unavailable in the MERFISH dataset, *Fos*-enriched clusters were defined using the following approach: For each behavioral category and sex, a threshold for *Fos* enrichment was set at the one-sided 95th percentile expression level. The proportion of cells above this threshold was calculated for each cell type, and Fisher’s exact test was performed to identify *Fos*-enriched clusters (*p* < 0.05 after multiple comparison correction). As *Fos*-enriched clusters were only observed in a subset of GABAergic clusters, subsequent analyses focused exclusively on the correspondence of GABAergic clusters between modalities.

To map MERFISH clusters onto scRNAseq data, reference (MERFISH) and query (scRNAseq, P50) datasets were aligned for each sex using the FindTransferAnchors (CCA1-30) and TransferData functions. This generated a weights matrix (*W*), which was used to transfer cell-type labels through the equation P = LW^T^, where L is a binary classification matrix and P represents label predictions. The imputed MERFISH cluster labels on scRNAseq cells enabled the inference of scRNAseq clusters associated with social behaviors. Genes selectively enriched in scRNAseq clusters linked to sexual behaviors were identified using the FindMarkers function.

The integrative analysis of MERFISH and scRNAseq data, along with enrichment analysis of hormonally associated DEGs (HA-DEGs), consistently demonstrated that Vgat+Esr1+clusters (defined as clusters where >50% of cells express Esr1) were transcriptionally dynamic and strongly linked to sexual behaviors. Consequently, downstream analyses primarily focused on Vgat+Esr1+clusters, while differences with other Vgat+ populations (e.g., Vgat+ Esr1- Ar+ or Vgat+ hormone RLow) were highlighted when relevant.

## Gene set activity analysis of scRNAseq data

To assess the aggregate expression of specific gene sets in single cells, we performed AUCell analysis (Aibar et al., 2017). AUCell calculates the Area Under the Curve (AUC) to determine whether a given gene set is enriched among the expressed genes in each cell. Using this approach, we computed gene set activity for hormonally associated DEGs (HA-DEGs; **Fig. 2g**), *Ar*-regulon in male-dimorphic genes (**Fig. 5h–j**), and adolescence-related versus unrelated dimorphic genes (**Fig. 6g–h**).

## Trajectory analysis of scRNAseq and HM-HCR FISH data

Monocle 3 was used to quantify adolescent transcriptional progression in scRNAseq and HM-HCR FISH datasets ^34^. For scRNAseq data, Seurat clustering results were used to separately analyze Vgat+Esr1+, Vgat+Esr1-Ar+, Vgat+hormoneR^Low^, and Vglut2+Esr1+ clusters in each sex. The preprocessing step involved normalizing and scaling the data using the preprocess_cds function, followed by PCA dimensional reduction based on hormonally associated DEGs (HA-DEGs; using the top 10–15 principal components). The reduce_dimension function was applied to generate UMAP embeddings, and a principal graph in UMAP space was constructed using the learn_graph function with default settings. The root node of the trajectory was assigned to cells from P23 or GDX groups, and pseudotime, representing transcriptional progression from the root state, was computed as the geodesic distance of each cell from the root node in the UMAP.

To test whether hormonally dependent transcriptional trajectories observed in Vgat+Esr1+ cells were also present in other cell types, similar Monocle trajectory analyses were conducted for Vgat+Esr1-Ar+, Vgat+hormoneR^Low^, and Vglut2+Esr1+ cells. Additionally, branches of transcriptional trajectories were analyzed due to the presence of two major branches originating from one or two subpopulations of Vgat+Esr1+ cells in both sexes (**Supplementary Fig. 5, 12**). Branch-specific DEG analysis identified genes enriched in each branch and those shared across branches (**Supplementary Fig. 5, 12**).

In the HM-HCR experiments, seven genes were selected from the top 50 HA-DEGs based on fold change and selectivity (ratio of pct1 to pct2), and two genes (male) or one gene (female) were selected from the top 10 genes enriched in P23 over P50. To confirm that these selected DEGs represented pubertal and adolescent transcriptional states, decoding accuracy of experimental conditions (age and hormonal states; detailed in Decoding Conditions from scRNAseq or HM-HCR FISH) was computed and compared between HCR DEGs and control DEGs (seven from the bottom 50 DEGs and two or one from the bottom 10 genes enriched in P23 over P50; **Supplementary Fig. 6k, m**).

Trajectory analysis of HM-HCR data was performed similarly to scRNAseq data using Monocle 3, focusing on Vgat+Esr1+ cells. The primary difference was that log1p values (not normalized by total UMI) were used, as most detected genes were HA-DEGs, and equivalent transcript numbers were not assumed for single cells. To quantify spatial-temporal dynamics of pubertal and adolescent transcriptional progression across ages, hormonal states, and sexes, the MPOA size was normalized across groups. Pseudotime values for the medial and lateral MPOA were computed to identify spatially specific patterns. To examine sex differences in the transcriptional onset of puberty and adolescence, Vgat+Esr1+ cells from P23, P28 (non-hormonally treated controls), and P50 groups were projected into the normalized MPOA space. A transcriptional maturation index was calculated for each P28 cell based on the pseudotime of its neighborhood P23 and P50 cells.

To jointly compute transcriptional trajectories from scRNAseq and HM-HCR data, scRNAseq data was imputed using HM-HCR data via Seurat V3 (**Supplementary Fig. 9**). In Vgat+Esr1+ cells, *Slc32a1* and one HM-HCR gene were excluded from the scRNAseq dataset. The FindTransferAnchors (CCA) and TransferData functions were used to identify anchors between the HM-HCR reference and the scRNAseq query, generating a weights matrix (*W*). Gene expression features were transferred as P = FW^T^, where F is the gene expression matrix and P is the predicted expression matrix. This process was iterated for all HM-HCR genes to generate predicted scRNAseq data (11–12 genes). The predicted scRNAseq data was validated against the real dataset using Pearson correlation coefficient analysis. Transcriptional trajectories were then learned from the predicted data using the same Monocle pipeline as described above.

## MELD analysis

To computationally validate the monocle trajectory analysis of scRNAseq data, we used the MELD Python package, which learns the transition of transcriptional states as the relative likelihood of conditions for each cell in the PHATE space (Potential of Heat-diffusion for Affinity-based Trajectory Embedding) ^38,39^. PHATE preserves local and distal relationships of transcriptional states, outperforming other embedding techniques in maintaining the structure of single-cell data. The input data consisted of log-normalized expression values and clustering metadata for Vgat+Esr1+ cells, generated from Seurat V3 analysis. The analysis proceeded in two main steps. First, data was embedded into low-dimensional PHATE space. Principal components (PCs) were computed using scprep.reduce.pca (number of components = 8–15), and PHATE embeddings were generated with phate.PHATE and phate_op.fit_transform using default settings. Second, the kernel density for each condition was estimated to quantify the likelihood of cells belonging to specific states. Conditions were categorized as mature (P50, P35) or immature (P23, GDX) (**Supplementary Fig. 6h-i, j, l**). Kernel density estimation was performed using meld.MELD (beta = 67, knn = 7) and meld_op.fit_transform with default settings. Relative likelihoods for each condition (the probability of observing a cell in each condition) were computed using the replicate_normalize_densities function, which applies L1 normalization across samples. These MELD analyses were used to measure hormonally associated adolescent trajectories, providing computational cross-validation for the monocle trajectory analysis (**Supplementary Fig. 6h-i, j, l**).

## DEG and trajectory analysis of scRNAseq in Esr1KO experiments

Data from *Esr1*KO groups were independently processed for quality control, including the removal of low-quality cells and doublets ^60^. Seurat V3 was used to perform clustering and identify neuronal clusters (detailed in Integrative Clustering and Differential Gene Expression Analysis). The dataset included 6,727 cells from *Esr1*KOF (female) and 6,464 cells from *Esr1*KOM (male). Neuronal cells were re-clustered to identify Vgat+ clusters. In females, six Vgat+ clusters (eVgat1–6), seven Vglut2+ clusters (eVglut1–7), and one mixed cluster (eMix1) were identified, while in males, 12 Vgat+ clusters (eVgat1–12), six Vglut2+ clusters (eVglut1–6), and two mixed clusters (eMix1–2) were identified (**Supplementary Fig. 11**; *Esr1*KOF: 2,182 neuronal cells; *Esr1*KOM: 4,761 neuronal cells).

To identify Vgat+ clusters in the *Esr1*KO group corresponding to specific clusters in the intact (viral-free) group, a Pearson correlation coefficient analysis was performed for each sex. This correlation matrix determined which clusters in the *Esr1*KO dataset corresponded to Vgat+Esr1+ clusters (e.g., Vgat2, 4, 16) or Vgat+hormoneR^Low^ clusters (e.g., Vgat14, 17, 20) in the intact group (**Supplementary Fig. 11**). To address potential experimental variations introduced by viral manipulations, this correspondence was cross-validated using anchor-based integrative analysis. Using the FindTransferAnchors (CCA1-30) and TransferData functions, anchors were identified between the intact group (reference) and *Esr1*KO group (query), constructing a weights matrix (*W*). Labels (cell-type classifications) were transferred using the equation P = LW^T^, where L is a binary classification matrix and P is the label prediction. Label imputation confirmed that eVgat1 and eVgat3 (female), and eVgat3 and eVgat4 (male), corresponded to the Vgat+Esr1+ population of intact groups. Similarly, eVgat5 (female) and eVgat9 (male) were closely related to the Vgat+hormoneR^Low^ clusters of P50 groups (**Supplementary Fig. 11**). Differentially expressed genes (DEGs) specific to *Esr1* were computed using Seurat’s FindMarkers function by comparing the gene expression of Vgat+Esr1+ clusters between intact P50 and *Esr1*KO groups. For trajectory analysis, Vgat+Esr1+ cells from all groups were combined and analyzed using Monocle 3. Principal components were computed based on the same HA-DEGs as described previously, and these were used to construct UMAP embeddings. In UMAP space, a principal graph was learned, root nodes were defined, pseudotime was computed, and branch-specific analysis was performed (detailed in Trajectory Analysis of scRNAseq and HM-HCR FISH Data).

## Deconstruction of gene regulatory network

To infer the cis-regulation of genes by transcription factors (TFs), we used the SCENIC computational framework ^31,53,66^. SCENIC was combined with cell-type-specific gene manipulation to causally infer the gene regulatory network (GRN). First, co-expression scores between TFs and target genes were computed using GENIE3 (runGenie3) based on scRNAseq expression data from Vgat+ cells at P50. TFs were ranked by the sum of their weights for hormonally associated DEGs (HA-DEGs; logFC > 0.25). Among all 952 TFs (female) and 915 TFs (male) in the dataset, *Esr1* achieved the highest sum score in both sexes. These results prompted further scRNAseq analysis of subjects with *Esr1* deleted in Vgat+ cells in the MPOA (detailed in Trajectory Analysis of scRNAseq in *Esr1*KO Experiments). After identifying Vgat+Esr1+ clusters, genes reduced by *Esr1* deletion (*Esr1*-DEGs) were identified using the FindMarkers function in Seurat V3. To determine which *Esr1*-DEGs were cis-regulated by *Esr1*, we applied the RcisTarget package. The RcisTarget database (mm9-tss-centered-10kb-7species.mc9nr.feather) was used to score motifs within the transcription start site (TSS) ±10 kb region and annotate associated TFs. Overrepresentation of each motif was assessed using the calcAUC function, and significantly enriched motifs were identified using the addMotifAnnotation function (normalized enrichment score threshold: NES ≥ 2). This analysis identified 123 genes in females and 98 genes in males that were significantly enriched with *Esr1* motifs. Of these, 9 (female) and 6 (male) genes were TFs (*Esr1*-TFs). To investigate whether *Esr1*-TFs could cis-regulate *Esr1*-DEGs, we repeated RcisTarget analysis for pairs of *Esr1*-TFs and *Esr1*-DEGs. These analyses allowed us to infer genes cis-regulated by *Esr1* or each *Esr1*-TF, collectively defining regulons for each TF. Using these regulons, we constructed the *Esr1*-GRN with the igraph package. In the *Esr1*-GRN, regulons were categorized and highlighted based on gene ontology (detailed in Ontology Analysis, **Fig. 8d-e, and Supplementary Fig. 13**). This approach provided a causal framework to understand the regulatory influence of *Esr1* and its downstream TFs on hormonally associated transcriptional networks.

## Single-cell consensus weighted gene co-expression network analysis

Single-cell consensus-weighted gene co-expression network analysis (scWGCNA) was employed to identify sex-specific co-expression networks in Vgat+Esr1+ cells ^57^. The analysis aimed to detect modules (groups of coexpressed genes) highly expressed in Vgat+Esr1+ clusters. To begin, metacells were constructed using the MetacellsByGroups function. Co-expression networks were then generated with the ConstructNetwork function, using the optimal soft-power threshold identified through TestSoftPowers. Module eigengenes (MEs), which represent the principal component of gene expression for each module, were computed using the ModuleEigengenes function. Module connectivity, calculated with the ModuleConnectivity function, was used to identify hub genes—those highly connected within each module. The top 25 hub genes and their connectivity were visualized using the ModuleNetworkPlot function. This analysis revealed nine modules in females and six modules in males that were highly expressed in Vgat+Esr1+ cells (Vgat2, 4, 16). Functional enrichment for each module was assessed using Enrichr via the RunEnrichr function ^58,59^. To directly compare modules between sexes, Jaccard similarity was computed across all module pairs to quantify sex-specific similarities and differences in gene co-expression networks.

## Ontology analysis

Ontology analysis was performed using Enrichr to examine the enrichment of functionally related genes within *Esr1*-DEGs (**Supplementary Fig. 13**). Gene ontology (GO) terms were identified from the GO Biological Process, GO Molecular Function, and Cellular Component databases. Detailed GO terms and their associated genes are reported in **Supplementary Fig. 13**. To simplify interpretation, GO terms were grouped into four categories: Gene Expression, Synapse/Excitability, Morphology, and Energy. Genes associated with multiple categories were manually assigned to the most relevant category to streamline classification and analysis.

## Decoding conditions from scRNAseq or HM-HCR FISH

To evaluate whether the selected features from trajectory analyses of scRNAseq and HM-HCR FISH experiments were sufficient to decode cellular conditions (e.g., age, sex, hormonal states, transcriptional maturity), we performed support vector machine (SVM) classification using the scikit-learn library. GridSearch with 10-fold cross-validation was applied to optimize the classification as described in previous studies ^67,68^. SVM models were tested using both linear and radial basis function (rbf) kernels.

Hyperparameters were optimized across a range of values for γ(10^-3^, 10^-2^, 10^-1^, 10^1^, 10^2^, 10^3^) and C (10^-3^, 10^-2^, 10^-1^, 10^1^, 10^2^, 10^3^). Input features for classification included differentially expressed genes (DEGs) used in constructing trajectories for scRNAseq and HM-HCR FISH. Decoding accuracies were compared against baseline accuracies calculated using randomized features (see Detailed Statistical Procedures).

## Detailed statistical procedures

Statistical analyses for each experiment are described in the methods and summarized below. Behavioral data and SVM classification analyses were performed using GraphPad Prism v9.0.2. Non-parametric tests were used to compare gene expression level distributions in scRNAseq and RNAscope data, conducted in R. No formal statistical tests were used to predetermine sample sizes.

**Fig. 1b**: Female: unpaired t-test. t(19)=5.7941, p < 0.001., Male: unpaired t-test. t(18)=3.8128, p=0.0029.

**Fig. 1c**: unpaired t-test. t(24)=0.7317, p=0.4714.

**Fig. 1d left:** unpaired t-test. t(24)=0.1302, p=0.8975.

**Fig. 1d right:** unpaired t-test. Subjects, which did not become receptive by P55, were given the value 60. T(24)=4.138, p=0.0004.

**Fig. 1e left:** Two-way repeated measures ANOVA followed by multiple comparisons. ANOVA revealed main effect of age (F (1.92778, 46.2667) = 21.8041, p<0.0001), main effect of group (F (1, 24) = 36.9300, p<0.0001) and interaction between group and age (F (2, 48) = 10.8824, p=0.0010). Holm-Sidak multi-comparison test was conducted. *p < 0.05, ***p < 0.001.

**Fig. 1e right:** Two-way repeated measures ANOVA followed by multiple comparisons. ANOVA revealed main effect of age (F (1.88961, 45.3507) = 23.1245, p<0.0001), main effect of group (F (1, 24) = 75.2342, p<0.0001) and interaction between group and age (F (2, 48) = 10.3659, p=0.0002). Holm-Sidak multi-comparison test was conducted. *p < 0.05, **p < 0.01, ***p < 0.001.

**Fig. 1f**: unpaired t-test. t(23)=0.09601, p=0.9243.

**Fig. 1g**: unpaired t-test. t(23)=4.534, p=0.0001.

**Fig. 1h left:** Two-way repeated measures ANOVA followed by multiple comparisons. ANOVA revealed main effect of age (F (3.94868, 90.8196) = 22.7095, p<0.0001), main effect of group (F (1, 23) = 66.9792, p<0.0001) and interaction between group and age (F (10, 230) = 18.7448, p<0.0001). Holm-Sidak multi-comparison test was conducted. *p < 0.05, **p < 0.01, ***p < 0.001.

**Fig. 1h right:** Two-way repeated measures ANOVA followed by multiple comparisons. ANOVA revealed main effect of age (F (4.27919, 98.4214) = 13.4311, p<0.0001), main effect of group (F (1, 23) = 19.1087, p=0.0002) and interaction between group and age (F (10, 230) = 8.95218, p<0.0001). Holm-Sidak multi-comparison test was conducted. *p < 0.05, **p < 0.01.

**Fig. 1j**: Female: unpaired t-test. t=2.523, p=0.025. Male: unpaired t-test. t= 4.7744, p=0.00024.

**Fig. 1k**: First VO age: unpaired t-test. t(18)=1.285, p=0.2152.

**Fig. 1l left:** First estrous age: unpaired t-test. t(18)=0.5142, p=0.6133.

**Fig. 1l right:** First receptive age: unpaired t-test. Subjects, which did not become receptive by P55, were given the value 65. t(18)=1.664, p=0.1134.

**Fig. 1m left:** The number of received intromissions: Two-way repeated measures ANOVA. ANOVA revealed main effect of age (F (1.933, 34.79) = 10.92, p=0.0002), no effect of group (F (1, 18) = 2.471, p=0.1334) and no interaction between group and age (F (2, 36) = 0.5656, p=0.5730).

**Fig. 1m right:** Receptivity: Two-way repeated measures ANOVA. ANOVA revealed main effect of age (F (1.990, 35.82) = 13.80, p<0.0001), no effect of group (F (1, 18) = 2.221, p=0.1534) and no interaction between group and age (F (2, 36) = 0.1445, p=0.8660).

**Fig. 1n**: First BPS age: unpaired t-test. t(16)=1.000, p=0.3322.

**Fig. 1o**: First mount age: unpaired t-test. t(16)=0.6030, p=0.5549.

**Fig. 1p left:** The number of mounts: Two-way repeated measures ANOVA. ANOVA revealed main effect of age (F (3.494, 55.90) = 39.72, p<0.0001), no effect of group (F (1, 16) = 0.001556, p=0.9690) and no interaction between group and age (F (10, 160) = 0.5442, p=0.8566).

**Fig. 1p right:** The number of thrusts: Two-way repeated measures ANOVA. ANOVA revealed main effect of age (F (3.831, 61.29) = 24.70, p<0.0001), no effect of group (F (1, 16) = 0.4016, p=0.5352) and no interaction between group and age (F (10, 160) = 0.3325, p=0.9713).

**Fig. 2e**: Linear regression analysis. Adjusted R-squared=0.51, p=0.00025.

**Fig. 2f**: Linear regression analysis. Adjusted R-squared for each hormone receptor gene was reported.

**Fig. 2g**: Kruskal-Wallis rank sum test followed by multiple comparisons at each Vgat+ cluster. Female:

Vgat2: Kruskal-Wallis chi-squared= 235.66, df=3, p<0.0001. Vgat4: Kruskal-Wallis chi-squared= 273.36, df=3, p<0.0001. Vgat16: Kruskal-Wallis chi-squared= 95.768, df=3, p<0.0001. Vgat6: Kruskal-Wallis chi-squared= 96.91, df=3, p<0.0001.

Vgat13: Kruskal-Wallis chi-squared= 8.2627, df=3, p= 0.04088. Vgat8: Kruskal-Wallis chi-squared= 5.4462, df=3, p= 0.1419.

Vgat1: Kruskal-Wallis chi-squared= 124.49, df=3, p<0.0001. Vgat10: Kruskal-Wallis chi-squared= 11.429, df=3, p= 0.009619. Vgat18: Kruskal-Wallis chi-squared= 8.5876, df=3, p= 0.03531. Vgat7: Kruskal-Wallis chi-squared= 2.5441, df=3, p= 0.4674.

Vgat3: Kruskal-Wallis chi-squared= 5.9714, df=3, p= 0.113. Vgat9: Kruskal-Wallis chi-squared= 6.7542, df=3, p= 0.08016. Vgat17: Kruskal-Wallis chi-squared= 7.4935, df=3, p= 0.05773. Vgat15: Kruskal-Wallis chi-squared= 3.2132, df=3, p= 0.3599. Vgat12: Kruskal-Wallis chi-squared= 5.316, df=3, p= 0.1501.

Vgat19: Kruskal-Wallis chi-squared= 1.7054, df=3, p= 0.6357.

male:

Vgat5: Kruskal-Wallis chi-squared= 3.6455, df=3, p= 0.3024. Vgat14: Kruskal-Wallis chi-squared= 1.9379, df=3, p= 0.5854. Vgat11: Kruskal-Wallis chi-squared= 12.301, df=3, p=0.006419. Vgat20: Kruskal-Wallis chi-squared= 5.3732, df=3, p = 0.1464.

Vgat2: Kruskal-Wallis chi-squared= 309.73, df=3, p<0.0001. Vgat4: Kruskal-Wallis chi-squared= 235.14, df=3, p<0.0001. Vgat16: Kruskal-Wallis chi-squared= 147.85, df=3, p<0.0001. Vgat6: Kruskal-Wallis chi-squared= 99.699, df=3, p<0.0001. Vgat13: Kruskal-Wallis chi-squared= 69.569, df=3, p<0.0001. Vgat8: Kruskal-Wallis chi-squared= 45.965, df=3, p<0.0001. Vgat1: Kruskal-Wallis chi-squared= 120.17, df=3, p<0.0001. Vgat10: Kruskal-Wallis chi-squared= 78.183, df=3, p<0.0001. Vgat18: Kruskal-Wallis chi-squared= 54.178, df=3, p<0.0001. Vgat7: Kruskal-Wallis chi-squared= 34.197, df=3, p<0.0001. Vgat3: Kruskal-Wallis chi-squared= 100.71, df=3, p<0.0001. Vgat9: Kruskal-Wallis chi-squared= 84.388, df=3, p<0.0001. Vgat17: Kruskal-Wallis chi-squared= 24.424, df=3, p<0.0001. Vgat15: Kruskal-Wallis chi-squared= 88.496, df=3, p<0.0001. Vgat12: Kruskal-Wallis chi-squared= 116.7, df=3, p<0.0001. Vgat19: Kruskal-Wallis chi-squared= 55.483, df=3, p<0.0001. Vgat5: Kruskal-Wallis chi-squared= 163.73, df=3, p<0.0001. Vgat14: Kruskal-Wallis chi-squared= 17.716, df=3, p=0.00050. Vgat11: Kruskal-Wallis chi-squared= 61.638, df=3, p<0.0001. Vgat20: Kruskal-Wallis chi-squared= 8.5476, df=3, p = 0.03595.

p values for multi-comparison tests (Bonferroni corrected) were reported for comparison between (P23, GDX) vs (P35, P50) when (P35, P50) was higher: ***p < 0.001.

**Fig. 3c**: Kruskal-Wallis rank sum test followed by multiple comparisons. Kruskal-Wallis chi-squared= 473.01, df=3, p<0.0001. p values for multi-comparison tests (Bonferroni corrected) were reported for comparison between (P23F, OVX) vs (P35F, P50F): ***p < 0.001.

**Fig. 3d**: Kruskal-Wallis rank sum test followed by multiple comparisons. Kruskal-Wallis chi-squared=448.36, df=3, p<0.0001. p values for multi-comparison tests (Bonferroni corrected) were reported for comparison between (P23F, OVX) vs (P35F, P50F): ***p < 0.001.

**Fig. 3e**: Kruskal-Wallis rank sum test followed by multiple comparisons. Kruskal-Wallis chi-squared= 1361.5, df=3, p<0.0001. p values for multi-comparison tests (Bonferroni corrected) were reported for comparison between (P23M, ORX) vs (P35M, P50M): ***p < 0.001.

**Fig. 3f**: Kruskal-Wallis rank sum test followed by multiple comparisons. Kruskal-Wallis chi-squared=712.23, df=3, p<0.0001. p values for multi-comparison tests (Bonferroni corrected) were reported for comparison between (P23M, ORX) vs (P35M, P50M): ***p < 0.001.

**Fig. 3h**: One-way ANOVA followed by multiple comparisons. ANOVA, F (3, 396) = 13016.7483, p<0.0001. Tukey’s multi-comparison test: real Vgat+Esr1+vs shuffle Vgat+Esr1+Vgat^+^, p< 0.0001. real Vgat+Esr1+vs real hormoneR^Low^, p< 0.0001. real Vgat+Esr1+vs shuffle hormoneR^Low^, p< 0.0001. shuffle Vgat+Esr1+vs real hormoneR^Low^, p< 0.0001. shuffle Vgat+Esr1+vs shuffle hormoneR^Low^, p< 0.0001. real hormoneR^Low^ vs shuffle hormoneR^Low^, p< 0.0001.

**Fig. 3j**: One-way ANOVA followed by multiple comparisons. ANOVA, F (3, 396) = 21372.43634, p<0.0001. Tukey’s multi-comparison test: real Vgat+Esr1+vs shuffle Vgat+Esr1+Vgat^+^, p< 0.0001. real Vgat+Esr1+vs real hormoneR^Low^, p< 0.0001. real Vgat+Esr1+vs shuffle hormoneR^Low^, p< 0.0001. shuffle Vgat+Esr1+vs real hormoneR^Low^, p< 0.0001. shuffle Vgat+Esr1+vs shuffle hormoneR^Low^, p< 0.0001. real hormoneR^Low^ vs shuffle hormoneR^Low^, p< 0.0001.

**Fig. 3l**: Wilcoxon test. Female only genes: W=0, p<0.0001; Male only genes: W=583, p=0.23; shared genes: W=0, p<0.0001.

**Fig. 3n**: Wilcoxon test. Female only genes: W=1414, p=0.057; Male only genes: W=0, p<0.0001; shared genes: W=0, p<0.0001.

**Fig. 5b**: Linear regression analysis. Adjusted R-squared=0.51, p=0.00025.

**Fig. 5c**: Linear regression analysis. Adjusted R-squared for each hormone receptor gene was reported.

**Fig. 5i**: unpaired Wilcoxon test. From left to right, p: <0.0001, <0.0001, <0.0001, =0.49, =1, =1, <0.0001, =0.08, =1, =0.40, =0.40, =1, =1, =1, =1, =1, =1, =1, =1, =1

**Fig. 5j**: Linear regression analysis. (Left) Adjusted R-squared=0.12, p=0.073. (right) Adjusted R-squared=0.53, p=0.00017.

Fig. 6a: All dimorphic genes: Kruskal-Wallis rank sum test followed by multiple comparisons. Kruskal-Wallis chi-squared= 38.68, df=3, p<0.0001.

Adolescence related dimorphic genes: Kruskal-Wallis rank sum test followed by multiple comparisons. Kruskal-Wallis chi-squared= 37.977, df=3, p<0.0001.

Adolescence unrelated dimorphic genes: Kruskal-Wallis rank sum test followed by multiple comparisons. Kruskal-Wallis chi-squared= 0.83392, df=3, p=0.8413.

p values for multi-comparison tests (Bonferroni corrected) were reported for comparison between (P23F, OVX) vs (P35F, P50F): ***p < 0.001.

Fig. 6b: All dimorphic genes: Kruskal-Wallis rank sum test followed by multiple comparisons. Kruskal-Wallis chi-squared= 214.46, df=3, p<0.0001.

Adolescence related dimorphic genes: Kruskal-Wallis rank sum test followed by multiple comparisons. Kruskal-Wallis chi-squared= 161.79, df=3, p<0.0001.

Adolescence unrelated dimorphic genes: Kruskal-Wallis rank sum test followed by multiple comparisons. Kruskal-Wallis chi-squared= 2.9488, df=3, p=0.3996.

p values for multi-comparison tests (Bonferroni corrected) were reported for comparison between (P23M, ORX) vs (P35M, P50M): ***p < 0.001.

**Fig. 6c**: One-way ANOVA followed by multiple comparisons.

Adolescence unrelated dimorphic genes: ANOVA, F (7, 792) = 34970, p<0.0001. Tukey’s multi-comparison test: real vs random shuffle, p<0.0001. real P23 vs real OVX, p=1. Real P35 vs real OVX, p=0.35. real P50 vs real OVX, p=0.44. real P35 vs real P23, p=0.49. real P50 vs real P23, p=0.59. real P50 vs real P35, p=1.

Adolescence related dimorphic genes: ANOVA, F (7, 792) = 5095, p<0.0001. Tukey’s multi-comparison test: real vs random shuffle, p<0.0001. real P23 vs real OVX, p<0.0001. Real P35 vs real OVX, p<0.0001. real P50 vs real OVX, p<0.0001. real P35 vs real P23, p<0.0001. real P50 vs real P23, p<0.0001. real P50 vs real P35, p<0.0001.

Post-hoc statistical results of Real P50 vs real others were highlighted when P50 was statistically larger than other real groups.

**Fig. 6d**: One-way ANOVA followed by multiple comparisons.

Adolescence unrelated dimorphic genes: ANOVA, F (7, 792) = 24080, p<0.0001. Tukey’s multi-comparison test: real vs random shuffle, p<0.0001. real P23 vs real OVX, p<0.0001. Real P35 vs real OVX, p<0.0001. real P50 vs real OVX, p<0.0001. real P35 vs real P23, p<0.0001. real P50 vs real P23, p<0.0001. real P50 vs real P35, p=0.186.

Adolescence related dimorphic genes: ANOVA, F (7, 792) = 5040, p<0.0001. Tukey’s multi-comparison test: real vs random shuffle, p<0.0001. real P23 vs real OVX, p<0.0001. Real P35 vs real OVX, p<0.0001. real P50 vs real OVX, p<0.0001. real P35 vs real P23, p<0.0001. real P50 vs real P23, p<0.0001. real P50 vs real P35, p<0.0001.

Post-hoc statistical results of Real P50 vs real others were highlighted when P50 was statistically larger than other real groups.

**Fig. 6f**: Kruskal-Wallis rank sum test followed by multiple comparisons. Kruskal-Wallis chi-squared= 767.39, df=7, p<0.0001. p values for multi-comparison tests (Bonferroni corrected) were reported for comparison between (P23F, OVX) vs (P35F, P50F) and (P23M, ORX) vs (P35M, P50M): ***p < 0.001. **Fig. 6g**: Kruskal-Wallis rank sum test followed by multiple comparisons.

(top) Kruskal-Wallis chi-squared= 1283.8, df=7, p<0.0001. p values for multi-comparison tests (Bonferroni corrected) were reported for comparison between (P35F, P50F) vs others. ***p < 0.001. (bottom) Kruskal-Wallis chi-squared= 446.72, df=7, p<0.0001. p values for multi-comparison tests (Bonferroni corrected) were reported for comparison between (P35F, P50F) vs others. ***p < 0.001.

**Fig. 6h**: Kruskal-Wallis rank sum test followed by multiple comparisons.

(top) Kruskal-Wallis chi-squared= 600.41, df=7, p<0.0001. p values for multi-comparison tests (Bonferroni corrected) were reported for comparison between (P35M, P50M) vs others. ***p < 0.001. (bottom) Kruskal-Wallis chi-squared= 484.69, df=7, p<0.0001. p values for multi-comparison tests (Bonferroni corrected) were reported for comparison between (P35M, P50M) vs others. ***p < 0.001.

**Fig. 7e**: Kruskal-Wallis rank sum test followed by multiple comparisons. Kruskal-Wallis chi-squared= 1583.4, df=4, p<0.0001. p values for multi-comparison tests (Bonferroni corrected) were reported for comparison between (P23F, OVX, Esr1KO) vs (P35F, P50F): ***p < 0.001.

**Fig. 7f**: Kruskal-Wallis rank sum test followed by multiple comparisons. Kruskal-Wallis chi-squared= 1194.6, df=4, p<0.0001. p values for multi-comparison tests (Bonferroni corrected) were reported for comparison between (P23F, OVX, Esr1KO) vs (P35F, P50F): ***p < 0.001.

**Fig. 7g**: Kruskal-Wallis rank sum test followed by multiple comparisons. Kruskal-Wallis chi-squared= 2990.2, df=4, p<0.0001. p values for multi-comparison tests (Bonferroni corrected) were reported for comparison between (P23M, ORX, Esr1KO) vs (P35M, P50M): ***p < 0.001.

**Fig. 7h**: Kruskal-Wallis rank sum test followed by multiple comparisons. Kruskal-Wallis chi-squared=1341.1, df=4, p<0.0001. p values for multi-comparison tests (Bonferroni corrected) were reported for comparison between (P23M, ORX, Esr1KO) vs (P35M, P50M): ***p < 0.001.

**Fig. 7j**: Wilcoxon test. Female only genes: W=0, p<0.0001; Male only genes: W=9327, p=0.17; shared genes: W=0, p<0.0001.

**Fig. 7l**: Wilcoxon test. Female only genes: W=2672, p=0.65; Male only genes: W=0, p<0.0001; shared genes: W=0, p<0.0001.

**Fig. 8c**: Female: unpaired Wilcoxon test. W=197164, p<0.0001. Male: unpaired Wilcoxon test. W=506476, p<0.0001.

**Fig. 8f**: unpaired Wilcoxon test. (Top to bottom) from left to right: W= 653278, 490664, 411540, 534038, 731136, 484492, 711128, 666599, 550366; P: <0.0001, <0.0001, =0.5057, <0.0001, <0.0001, <0.0001, <0.0001, <0.0001, <0.0001

In the figure, p values were reported as the number of stars when values in the cre group were lower than those in control.

**Fig. 8g**: unpaired Wilcoxon test. (Top to bottom) from left to right: W= 154110, 96298, 154052, 94662, 25612, 116958, 133079, 139160, 83886; P: <0.0001, =0.0026, <0.0001, =0.0102, <0.0001, <0.0001, <0.0001, <0.0001, =0.73

In the figure, p values were reported as the number of stars when values in the cre group were lower than those in control.

**Supplementary Fig. 1a:** paired t-test. t(11)=-0.523 , p=0.612.

**Supplementary Fig. 1c:** Two-way repeated measures ANOVA followed by multiple comparisons. ANOVA revealed main effect of age (F (2, 48) = 11.4416728, p<0.0001), main effect of group (F (1, 24) = 16.8147242, p<0.001) and interaction between group and age (F (2, 48) = 0.5084058, p=0.60). Tukey multi-comparison test was conducted. *p < 0.05.

**Supplementary Fig. 1d left:** unpaired t-test. t(24)= -2.966, p=0.0067

**Supplementary Fig. 1d right:** unpaired t-test. t(15.466)= -3.7502, p= 0.0018

**Supplementary Fig. 1e top:** Body weight: Two-way repeated measures ANOVA. ANOVA revealed main effect of age (F (3.062, 73.49) = 51.24, p<0.0001), no effect of group (F (1, 24) = 1.366, p=0.2539) and no interaction between group and age (F (12, 288) = 0.2786, p=0.9922). Locomotion: unpaired t-test. t(24)=1.134, p=0.2682. Time investigating male: unpaired t-test. t(24)=0.2027, p=0.8411. Time investigating female: unpaired t-test. t(24)=1.499, p=0.1469. Social Preference: unpaired t-test. t(24)=0.8857, p=0.3846. Time in open arm: unpaired t-test. t(14)=0.2707, p=0.7906.

**Supplementary Fig. 1e bottom:** Body weight: Two-way repeated measures ANOVA followed by multiple comparisons. ANOVA revealed main effect of age (F (3.182, 73.19) = 202.5, p<0.0001), no effect of group (F (1, 23) = 1.788, p=0.1942) and interaction between group and age (F (12, 276) = 3.375, p=0.0001). Holm-Sidak multi-comparison test was conducted. No significance was observed at any age between groups. Locomotion: unpaired t-test. t(23)=0.06150, p=0.9515. Time investigating male: unpaired t-test. t(23)=0.5738, p=0.5717. Time investigating female: unpaired t-test. t(23)=2.048, p=0.0521. Social Preference: unpaired t-test. t(23)=1.563, p=0.1317. Time in open arm: unpaired t-test. t(14)=0.4188, p=0.6817.

**Supplementary Fig. 1f top:** Body weight: Two-way repeated measures ANOVA. ANOVA revealed main effect of age (F (4.942, 88.95) = 104.5, p<0.0001), no effect of group (F (1, 18) = 2.849, p=0.1087) and no interaction between group and age (F (12, 216) = 1.006, p=0.4442). Locomotion: unpaired t-test. t(18)=0.7283, p=0.4758.Time investigating male: unpaired t-test. t(18)=1.060, p=0.3030. Time investigating female: unpaired t-test. t(18)=0.5965, p=0.5583. Social Preference: unpaired t-test. t(18)=1.684, p=0.1094. Time in open arm: unpaired t-test. t(18)=0.2933, p=0.7727.

**Supplementary Fig. 1f bottom:** Body weight: Two-way repeated measures ANOVA followed by multiple comparisons. ANOVA revealed main effect of age (F (4.126, 66.02) = 197.4, p<0.0001), main effect of group (F (1, 16) = 8.571, p=0.0099) and interaction between group and age (F (12, 192) = 9.574, p<0.0001). Holm-Sidak multi-comparison test was conducted. *p < 0.05. Locomotion: unpaired t-test. t(16)=1.826, p=0.0865. Time investigating male: unpaired t-test. t(16)=0.9004, p=0.3813. Time investigating female: unpaired t-test. t(16)=0.1816, p=0.8582. Social Preference: unpaired t-test. t(16)=0.5261, p=0.6061. Time in open arm: unpaired t-test. t(10)=0.8410, p=0.4200.

**Supplementary Fig. 1g:** #Esr1 and # intromission in female: Linear regression analysis. Adjusted R-squared=0.52, p=0.00014. #Esr1 and # receptivity in female: Linear regression analysis. Adjusted R-squared=0.48, p=0.00028. #Esr1 and # mounts in male: Linear regression analysis. Adjusted R-squared=0.29, p=0.0086. #Esr1 and # thrusts in male: Linear regression analysis. Adjusted R-squared=0.20, p=0.027.

**Supplementary Fig. 1h:** #Esr1 and # intromission in female: Linear regression analysis. Adjusted R-squared=-0.031, p=0.47. #Esr1 and # receptivity in female: Linear regression analysis. Adjusted R-squared=-0.033, p=0.48. #Esr1 and # mounts in male: Linear regression analysis. Adjusted R-squared= -0.056, p=0.76. #Esr1 and # thrusts in male: Linear regression analysis. Adjusted R-squared=0.078, p=0.14.

**Supplementary Fig. 2i:** unpaired Wilcoxon test. W=396364535, p<0.0001.

**Supplementary Fig. 3d left:** one-way repeated measures ANOVA followed by multiple comparisons.

Cluster1: ANOVA, F (135, 405) = 2.9438259, p<0.0001. Holm-Sidak multi-comparison test: P23 vs OVX, t(135)= 0.073054964, p= 0.9419. P23 vs P35, t(135)= 6.6450382, p <0.0001. P23 vs P50, t(135)= 12.347378, p <0.0001. P35 vs OVX, t(135)= 4.0952302, p< 0.0001. P50 vs OVX, t(135)= 13.269141, p <0.0001. P35 vs P50, t(135)= 11.17 6780, p <0.0001.

Cluster2: ANOVA, F (282, 846) = 4.6794342, P<0.0001. Holm-Sidak multi-comparison test: P23 vs OVX, t(282)= 22.482676, p <0.0001. P23 vs P35, t(282)= 29.289408, p <0.0001. P23 vs P50, t(282)= 31.215903, p <0.0001. P35 vs OVX, t(282)= 5.2700226, p< 0.0001. P50 vs OVX, t(282)= 12.850838, p <0.0001. P35 vs P50, t(282)= 9.6808179, p <0.0001.

Cluster3: ANOVA, F (354, 1062) = 3.9834674, p<0.0001. Holm-Sidak multi-comparison test: P23 vs OVX, t(354)= 51.772642, p <0.0001. P23 vs P35, t(354)= 20.469338, p <0.0001. P23 vs P50, t(354)= 40.905873, p <0.0001. P35 vs OVX, t(354)= 28.730343, p< 0.0001. P50 vs OVX, t(354)= 11.831025, p <0.0001. P35 vs P50, t(354)= 20.702854, p <0.0001.

Cluster4: ANOVA, F (356, 1068) = 4.6642539, P<0.0001. Holm-Sidak multi-comparison test: P23 vs OVX, t(356)= 2.6050178, p <0.0096. P23 vs P35, t(356)= 23.110498, p <0.0001. P23 vs P50, t(356)= 6.1529826, p <0.0001. P35 vs OVX, t(356)= 33.395442, p< 0.0001. P50 vs OVX, t(356)= 11.259059, p <0.0001. P35 vs P50, t(356)= 30.387666, p <0.0001.

Cluster5: ANOVA, F (29, 87) = 6.4594382, p<0.0001. Holm-Sidak multi-comparison test: P23 vs OVX, t(29)= 3.9693469, p = 0.0017. P23 vs P35, t(29)= 1.4257659, p = 0.3021. P23 vs P50, t(29)= 0.040574193, p = 0.9679. P35 vs OVX, t(29)= 6.7801762, p< 0.0001. P50 vs OVX, t(29)= 7.3425963, p <0.0001. P35 vs P50, t(29)= 3.2891624, p = 0.0079.

Cluster6: ANOVA, F (339, 1017) = 3.7357024, p<0.0001. Holm-Sidak multi-comparison test: P23 vs OVX, t(339)= 27.925103, p < 0.0001. P23 vs P35, t(339)= 1.1969343, p = 0.2322. P23 vs P50, t(339)= 39.356609, p < 0.0001. P35 vs OVX, t(339)= 26.966204, p< 0.0001. P50 vs OVX, t(339)= 24.477635, p <0.0001. P35 vs P50, t(339)= 41.561976, p < 0.0001.

**Supplementary Fig. 3d right:** one-way repeated measures ANOVA followed by multiple comparisons. Cluster1: ANOVA, F (770, 2310) = 4.8206939, p<0.0001. Holm-Sidak multi-comparison test: P23 vs ORX, t(770)= 12.113758, p < 0.0001. P23 vs P35, t(770)= 9.6124052, p < 0.0001. P23 vs P50, t(770)= 26.901991, p < 0.0001. P35 vs ORX, t(770)= 28.711090, p< 0.0001. P50 vs ORX, t(770)= 54.978801, p <0.0001. P35 vs P50, t(770)= 31.862981, p < 0.0001.

Cluster2: ANOVA, F (1089, 3267) = 5.6620007, p<0.0001. Holm-Sidak multi-comparison test: P23 vs ORX, t(1089)= 77.978449, p < 0.0001. P23 vs P35, t(1089)= 48.804692, p < 0.0001. P23 vs P50, t(1089)= 55.618808, p < 0.0001. P35 vs ORX, t(1089)= 36.331287, p< 0.0001. P50 vs ORX, t(1089)= 19.946091, p <0.0001. P35 vs P50, t(1089)= 10.943465, p < 0.0001.Cluster3: ANOVA, F (189, 567) = 3.4899665, p<0.0001. Holm-Sidak multi-comparison test: P23 vs ORX, t(189)= 5.6579721, p < 0.0001. P23 vs P35, t(189)= 11.699347, p < 0.0001. P23 vs P50, t(189)= 8.0593044, p < 0.0001. P35 vs ORX, t(189)= 14.879007, p< 0.0001. P50 vs ORX, t(189)= 12.488563, p <0.0001. P35 vs P50, t(189)= 14.346162, p < 0.0001.

Cluster4: ANOVA, F (1030, 3090) = 6.0976581, p<0.0001. Holm-Sidak multi-comparison test: P23 vs ORX, t(1030)= 30.413779, p < 0.0001. P23 vs P35, t(1030)= 37.654898, p < 0.0001. P23 vs P50, t(1030)= 38.951160, p < 0.0001. P35 vs ORX, t(1030)= 11.231385, p< 0.0001. P50 vs ORX, t(1030)= 13.001623, p <0.0001. P35 vs P50, t(1030)= 5.5005486, p < 0.0001.

Cluster5: ANOVA, F (118, 354) = 3.3165807, p<0.0001. Holm-Sidak multi-comparison test: P23 vs ORX, t(118)= 6.3281474, p < 0.0001. P23 vs P35, t(118)= 7.1294177, p < 0.0001. P23 vs P50, t(118)= 6.5731275, p < 0.0001. P35 vs ORX, t(118)= 2.0573177, p= 0.0419. P50 vs ORX, t(118)= 11.363377, p <0.0001. P35 vs P50, t(118)= 14.575549, p < 0.0001.

**Supplementary Fig. 3e left:** one-way repeated measures ANOVA followed by multiple comparisons.

Cluster1: ANOVA, F (106, 318) = 1.784, p<0.0001. Holm-Sidak multi-comparison test: P23 vs OVX, t(106)= 17.25, p<0.0001. P23 vs P35, t(106)= 5.938, p<0.0001. P23 vs P50, t(106)= 3.823, p =0.0004. P35 vs OVX, t(106)= 17.54, p< 0.0001. P50 vs OVX, t(106)= 9.249, p <0.0001. P35 vs P50, t(106)= 0.9242, p =0.3575.

Cluster2: ANOVA, F (87, 261) = 6.329, p<0.0001. Holm-Sidak multi-comparison test: P23 vs OVX, t(87)= 10.19, p<0.0001. P23 vs P35, t(87)= 8.772, p<0.0001. P23 vs P50, t(87)= 10.93, p<0.0001. P35 vs OVX, t(87)= 0.6321, p=0.5291. P50 vs OVX, t(87)= 7.370, p <0.0001. P35 vs P50, t(87)= 6.112, p<0.0001.

Cluster3: ANOVA, F (110, 330) = 2.172, p<0.0001. Holm-Sidak multi-comparison test: P23 vs OVX, t(110)= 4.682, p<0.0001. P23 vs P35, t(110)= 16.20, p<0.0001. P23 vs P50, t(110)= 3.059, p=0.0056. P35 vs OVX, t(110)= 11.90, p <0.0001. P50 vs OVX, t(110)= 1.708, p =0.0904. P35 vs P50, t(110)= 15.13, p<0.0001.

Cluster4: ANOVA, F (11, 33) = 6.390, p<0.0001. Holm-Sidak multi-comparison test: P23 vs OVX, t(11)= 3.374, p=0.0185. P23 vs P35, t(11)= 4.954, p=0.0026. P23 vs P50, t(11)= 3.743, p=0.0129. P35 vs OVX, t(11)= 4.328, p =0.0060. P50 vs OVX, t(11)= 1.324, p =0.2122. P35 vs P50, t(11)= 2.866, p=0.0305.

Cluster5: ANOVA, F (201, 603) = 4.559, p<0.0001. Holm-Sidak multi-comparison test: P23 vs OVX, t(201)= 29.20, p<0.0001. P23 vs P35, t(201)= 14.46, p<0.0001. P23 vs P50, t(201)= 32.37, p<0.0001. P35 vs OVX, t(201)= 26.25, p <0.0001. P50 vs OVX, t(201)= 2.931, p =0.0038. P35 vs P50, t(201)= 17.71, p<0.0001.

**Supplementary Fig. 3e right:** one-way repeated measures ANOVA followed by multiple comparisons. Cluster1: F (706, 2118) = 5.635, p<0.0001. Holm-Sidak multi-comparison test: P23 vs ORX, t(706)= 52.31, p < 0.0001. P23 vs P35, t(706)= 28.25, p < 0.0001. P23 vs P50, t(706)= 33.39, p < 0.0001. P35 vs ORX, t(706)= 39.25, p< 0.0001. P50 vs ORX, t(706)= 19.27, p <0.0001. P35 vs P50, t(706)= 10.93, p < 0.0001.

Cluster2: ANOVA, F (520, 1560) = 5.850, p<0.0001. Holm-Sidak multi-comparison test: P23 vs ORX, t(520)= 13.78, p < 0.0001. P23 vs P35, t(520)= 21.64, p < 0.0001. P23 vs P50, t(520)= 22.69, p < 0.0001. P35 vs ORX, t(520)= 9.028, p< 0.0001. P50 vs ORX, t(520)= 10.51, p <0.0001. P35 vs P50, t(520)= 5.259, p < 0.0001.

Cluster3: ANOVA, F (176, 528) = 3.522, p<0.0001. Holm-Sidak multi-comparison test: P23 vs ORX, t(176)= 15.68, p < 0.0001. P23 vs P35, t(176)= 4.169, p < 0.0001. P23 vs P50, t(176)= 8.638, p < 0.0001. P35 vs ORX, t(176)= 17.51, p< 0.0001. P50 vs ORX, t(176)= 26.15, p <0.0001. P35 vs P50, t(176)= 13.70, p < 0.0001.

Cluster4: ANOVA, F (164, 492) = 5.971, p<0.0001. Holm-Sidak multi-comparison test: P23 vs ORX, t(164)= 29.20, p < 0.0001. P23 vs P35, t(164)= 33.47, p < 0.0001. P23 vs P50, t(164)= 31.91, p < 0.0001. P35 vs ORX, t(164)= 5.824, p< 0.0001. P50 vs ORX, t(164)= 9.851, p <0.0001. P35 vs P50, t(164)= 4.902, p < 0.0001.

Cluster5: ANOVAF (55, 165) = 3.797, p<0.0001. Holm-Sidak multi-comparison test: P23 vs ORX, t(55)= 6.306, p < 0.0001. P23 vs P35, t(55)= 6.165, p < 0.0001. P23 vs P50, t(55)= 1.518, p = 0.1348. P35 vs ORX, t(55)= 3.378, p= 0.0027. P50 vs ORX, t(55)= 11.23, p <0.0001. P35 vs P50, t(55)= 7.715, p < 0.0001.

**Supplementary Fig. 3f:** Linear regression analysis. Adjusted R-squared=0.43, p=0.013. **Supplementary Fig. 3g:** Linear regression analysis. Adjusted R-squared for each hormone receptor gene was reported.

**Supplementary Fig. 4a, b:** Fisher’s exact test was conducted to compare each cluster with all clusters. P-values were Bonferroni corrected. **p < 0.01, ***p < 0.001.

p values for multi-comparison tests (Bonferroni corrected) were reported for comparison between (P23F, OVX) vs (P35F, P50F): ***p < 0.001.

**Supplementary Fig. 5c left branch1 genes:** Kruskal-Wallis rank sum test followed by multiple comparisons. Kruskal-Wallis chi-squared= 142.61, df=3, p<0.0001. p values for multi-comparison tests (Bonferroni corrected) were reported for comparison between (P23F, OVX) vs (P35F, P50F): ***p < 0.001.

**Supplementary Fig. 5c left branch2 genes:** Kruskal-Wallis rank sum test followed by multiple comparisons. Kruskal-Wallis chi-squared= 133.17, df=3, p<0.0001. p values for multi-comparison tests (Bonferroni corrected) were reported for comparison between (P23F, OVX) vs (P35F, P50F).

**Supplementary Fig. 5c left branch common genes:** Kruskal-Wallis rank sum test followed by multiple comparisons. Kruskal-Wallis chi-squared= 128.99, df=3, p<0.0001. p values for multi-comparison tests (Bonferroni corrected) were reported for comparison between (P23F, OVX) vs (P35F, P50F): ***p < 0.001.

**Supplementary Fig. 5c right branch1 genes:** Kruskal-Wallis rank sum test followed by multiple comparisons. Kruskal-Wallis chi-squared= 27.447, df=3, p<0.0001. p values for multi-comparison tests (Bonferroni corrected) were reported for comparison between (P23F, OVX) vs (P35F, P50F).

**Supplementary Fig. 5c right branch2 genes:** Kruskal-Wallis rank sum test followed by multiple comparisons. Kruskal-Wallis chi-squared= 510.85, df=3, p<0.0001. p values for multi-comparison tests (Bonferroni corrected) were reported for comparison between (P23F, OVX) vs (P35F, P50F): ***p < 0.001.

**Supplementary Fig. 5c right branch common genes:** Kruskal-Wallis rank sum test followed by multiple comparisons. Kruskal-Wallis chi-squared= 117.43, df=3, p<0.0001. p values for multi-comparison tests (Bonferroni corrected) were reported for comparison between (P23F, OVX) vs (P35F, P50F): ***p < 0.001.

**Supplementary Fig. 5d left branch1 genes:** Kruskal-Wallis rank sum test followed by multiple comparisons. Kruskal-Wallis chi-squared= 278.57, df=3, p<0.0001. p values for multi-comparison tests (Bonferroni corrected) were reported for comparison between (P23M, ORX) vs (P35M, P50M): ***p < 0.001.

**Supplementary Fig. 5d left branch2 genes:** Kruskal-Wallis rank sum test followed by multiple comparisons. Kruskal-Wallis chi-squared= 192.7, df=3, p<0.0001. p values for multi-comparison tests (Bonferroni corrected) were reported for comparison between (P23M, ORX) vs (P35M, P50M).

**Supplementary Fig. 5d left branch common genes:** Kruskal-Wallis rank sum test followed by multiple comparisons. Kruskal-Wallis chi-squared= 137.74, df=3, p<0.0001. p values for multi-comparison tests (Bonferroni corrected) were reported for comparison between (P23M, ORX) vs (P35M, P50M): ***p < 0.001.

**Supplementary Fig. 5d right branch1 genes:** Kruskal-Wallis rank sum test followed by multiple comparisons. Kruskal-Wallis chi-squared= 72.355, df=3, p<0.0001. p values for multi-comparison tests (Bonferroni corrected) were reported for comparison between (P23M, ORX) vs (P35M, P50M).

**Supplementary Fig. 5d right branch2 genes:** Kruskal-Wallis rank sum test followed by multiple comparisons. Kruskal-Wallis chi-squared= 646.97, df=3, p<0.0001. p values for multi-comparison tests (Bonferroni corrected) were reported for comparison between (P23M, ORX) vs (P35M, P50M): ***p < 0.001.

**Supplementary Fig. 5d right branch common genes:** Kruskal-Wallis rank sum test followed by multiple comparisons. Kruskal-Wallis chi-squared=139.53, df=3, p<0.0001. p values for multi-comparison tests (Bonferroni corrected) were reported for comparison between (P23M, ORX) vs (P35M, P50M): ***p < 0.001.

**Supplementary Fig. 5e branch1:** Kruskal-Wallis rank sum test followed by multiple comparisons. Kruskal-Wallis chi-squared= 435.41, df=3, p<0.0001. p values for multi-comparison tests (Bonferroni corrected) were reported for comparison between (P23F, OVX) vs (P35F, P50F): ***p < 0.001.

**Supplementary Fig. 5e branch2:** Kruskal-Wallis rank sum test followed by multiple comparisons. Kruskal-Wallis chi-squared= 271.89, df=3, p<0.0001. p values for multi-comparison tests (Bonferroni corrected) were reported for comparison between (P23F, OVX) vs (P35F, P50F): ***p < 0.001.

**Supplementary Fig. 5f branch1:** Kruskal-Wallis rank sum test followed by multiple comparisons. Kruskal-Wallis chi-squared= 565.82, df=3, p<0.0001. p values for multi-comparison tests (Bonferroni corrected) were reported for comparison between (P23M, ORX) vs (P35M, P50M): ***p < 0.001.

**Supplementary Fig. 5f branch2:** Kruskal-Wallis rank sum test followed by multiple comparisons. Kruskal-Wallis chi-squared= 540.99, df=3, p<0.0001. p values for multi-comparison tests (Bonferroni corrected) were reported for comparison between (P23M, ORX) vs (P35M, P50M): ***p < 0.001.

**Supplementary Fig. 5g branch1:** unpaired t-test. T(152.39)= 216.35, p<0.0001. **Supplementary Fig. 5g branch2:** unpaired t-test. T(147.5)= 153.52, p<0.0001. **Supplementary Fig. 5h branch1:** unpaired t-test. T(130.85)= 244.88, p<0.0001. **Supplementary Fig. 5h branch2:** unpaired t-test. T(124.65)= 204.9, p<0.0001. **Supplementary Fig. 5i branch1 AUCell score:** unpaired Wilcoxon test. p<0.0001. **Supplementary Fig. 5i branch2 AUCell score:** unpaired Wilcoxon test. p<0.0001. **Supplementary Fig. 5i branch1 SVM:** unpaired t-test. T(134.62)= 142.72, p<0.0001. **Supplementary Fig. 5i branch2 SVM:** unpaired t-test. T(176.75)= 66.219, p<0.0001. **Supplementary Fig. 5j branch1 AUCell score:** unpaired Wilcoxon test. p<0.0001.

**Supplementary Fig. 5j branch2 AUCell score:** unpaired Wilcoxon test. P=0.0050. **Supplementary Fig. 5j branch1 SVM:** unpaired t-test. T(164.27)= 83.34, p<0.0001. **Supplementary Fig. 5j branch2 SVM:** unpaired t-test. T(190.61)= 71.154, p<0.0001.

**Supplementary Fig. 6d:** One-way ANOVA followed by multiple comparisons. ANOVA, F (3, 396) = 15075, p<0.0001. Tukey’s multi-comparison test: real male vs shuffle male, p< 0.0001. real male vs real female, p< 0.0001. real male vs shuffle female, p< 0.0001. shuffle male vs real female, p< 0.0001. shuffle male vs shuffle female, p= 0.6720. real female vs shuffle female, p< 0.0001.

**Supplementary Fig. 6f:** Kruskal-Wallis rank sum test followed by multiple comparisons. Kruskal-Wallis chi-squared= 830.08, df=7, p<0.0001. p values for multi-comparison tests (Bonferroni corrected) were reported for comparison between (P23, GDX) vs (P35, P50): ***p < 0.001.

**Supplementary Fig. 6j:** Kruskal-Wallis rank sum test followed by multiple comparisons. Kruskal-Wallis chi-squared= 570.12, df=3, p<0.0001.

**Supplementary Fig. 6k:** unpaired t-test. T(198)= 61.044, p<0.0001.

**Supplementary Fig. 6l:** Kruskal-Wallis rank sum test followed by multiple comparisons. Kruskal-Wallis chi-squared= 643.82, df=3, p<0.0001.

p values for multi-comparison tests (Bonferroni corrected) were reported for comparison between (P23M, ORX) vs (P35M, P50M): ***p < 0.001.

**Supplementary Fig. 6m:** unpaired t-test. T(198)= 99.879, p<0.0001.

**Supplementary Fig. 6o:** Kruskal-Wallis rank sum test followed by multiple comparisons. Kruskal-Wallis chi-squared= 228.06, df=3, p<0.0001.

p values for multi-comparison tests (Bonferroni corrected) were reported for comparison between (P23F, OVX) vs (P35F, P50F): ***p < 0.001.

**Supplementary Fig. 6q:** Kruskal-Wallis rank sum test followed by multiple comparisons. Kruskal-Wallis chi-squared= 310.29, df=3, p<0.0001.

p values for multi-comparison tests (Bonferroni corrected) were reported for comparison between (P23M, ORX) vs (P35M, P50M): ***p < 0.001.

**Supplementary Fig. 7d:** Kruskal-Wallis rank sum test followed by multiple comparisons. Kruskal-Wallis chi-squared= 146.73, df=3, p<0.0001.

p values for multi-comparison tests (Bonferroni corrected) were reported for comparison between (P23F, OVX) vs (P35F, P50F): ***p < 0.001.

**Supplementary Fig. 7e:** Kruskal-Wallis rank sum test followed by multiple comparisons. Kruskal-Wallis chi-squared= 515.86, df=3, p<0.0001.

p values for multi-comparison tests (Bonferroni corrected) were reported for comparison between (P23M, ORX) vs (P35M, P50M): ***p < 0.001.

**Supplementary Fig. 9a:** Kruskal-Wallis rank sum test followed by multiple comparisons.

*Pdzrn4*: Kruskal-Wallis chi-squared=1998, df=5, p<0.001. *Creb3l1*: Kruskal-Wallis chi-squared=507.5, df=5, p<0.001. *Sccpdh*: Kruskal-Wallis chi-squared=2517.7, df=5, p<0.001. *Esr1*: Kruskal-Wallis chi-squared=166.37, df=5, p<0.001. *Socs2*: Kruskal-Wallis chi-squared=1609.2, df=5, p<0.001. *Pgr*: Kruskal-Wallis chi-squared=1928.7, df=5, p<0.001. *Tmem35a*: Kruskal-Wallis chi-squared=1450.7, df=5, p<0.001. *Nts*: Kruskal-Wallis chi-squared=631.36, df=5, p<0.001. *Slc32a1*: Kruskal-Wallis chi-squared=133.17, df=5, p<0.001. *Alcam*: Kruskal-Wallis chi-squared=657.22, df=5, p<0.001. *Ar*: Kruskal-Wallis chi-squared=546.52, df=5, p<0.001.

p values for multi-comparison tests (Bonferroni corrected) were reported for comparison between (P23F, P28C, OVX) vs (P28E, P35F, P50F): *p < 0.05, **p < 0.01, ***p < 0.001.

**Supplementary Fig. 9b:** Kruskal-Wallis rank sum test followed by multiple comparisons.

*Sytl4*: Kruskal-Wallis chi-squared=1532.5, df=5, p<0.001. *Acvr1c*: Kruskal-Wallis chi-squared=936.37, df=5, p<0.001. *Greb1*: Kruskal-Wallis chi-squared=512.3, df=5, p<0.001. *Nrip*: Kruskal-Wallis chi-squared=759.41, df=5, p<0.001. *Lamp5*: Kruskal-Wallis chi-squared=1052.2, df=5, p<0.001. *Apoc3*: Kruskal-Wallis chi-squared=938.64, df=5, p<0.001. *Pgr*: Kruskal-Wallis chi-squared=606.25, df=5, p<0.001. *Esr1*: Kruskal-Wallis chi-squared=87.281, df=5, p<0.001. *Slc32a1*: Kruskal-Wallis chi-squared=192.37, df=5, p<0.001. *Npy2r*: Kruskal-Wallis chi-squared=528.2, df=5, p<0.001. *Pgr151*: Kruskal-Wallis chi-squared=1920.3, df=5, p<0.001. *Ar*: Kruskal-Wallis chi-squared=235.83, df=5, p<0.001.

p values for multi-comparison tests (Bonferroni corrected) were reported for comparison between (P23M, P28C, ORX) vs (P28T, P35M, P50M): *p < 0.05, **p < 0.01, ***p < 0.001.

**Supplementary Fig. 9d:** One-way ANOVA followed by multiple comparisons. ANOVA, F (3, 396) = 43666.7034, p<0.0001. Tukey’s multi-comparison test: real Vgat+Esr1+vs shuffle Vgat+Esr1+Vgat^+^, p< 0.0001. real Vgat+Esr1+vs real hormoneR^Low^, p< 0.0001. real Vgat+Esr1+vs shuffle hormoneR^Low^, p< 0.0001. shuffle Vgat+Esr1+vs real hormoneR^Low^, p< 0.0001. shuffle Vgat+Esr1+vs shuffle hormoneR^Low^, p< 0.0001. real hormoneR^Low^ vs shuffle hormoneR^Low^, p< 0.0001.

**Supplementary Fig. 9g:** One-way ANOVA followed by multiple comparisons. ANOVA, F (3, 396) = 36690.76438, p<0.0001. Tukey’s multi-comparison test: real Vgat+Esr1+vs shuffle Vgat+Esr1+Vgat^+^, p< 0.0001. real Vgat+Esr1+vs real hormoneR^Low^, p< 0.0001. real Vgat+Esr1+vs shuffle hormoneR^Low^, p< 0.0001. shuffle Vgat+Esr1+vs real hormoneR^Low^, p< 0.0001. shuffle Vgat+Esr1+vs shuffle hormoneR^Low^, p< 0.0001. real hormoneR^Low^ vs shuffle hormoneR^Low^, p< 0.0001.

**Supplementary Fig. 9n:** unpaired t-test. T(198)=91.01, p<0.0001.

**Supplementary Fig. 9q:** unpaired t-test. T(198)=157.2, p<0.0001.

**Supplementary Fig. 9r:** Kruskal-Wallis rank sum test followed by multiple comparisons. Kruskal-Wallis chi-squared= 2815.7, df=11, p<0.0001.

p values for multi-comparison tests (Bonferroni corrected) were reported for comparisons between (P23F, P28FC, OVX) vs (P28FE, P35F, P50F) in M or L and for comparisons between M and L within each group: **p < 0.01, ***p < 0.001.

**Supplementary Fig. 9s:** Kruskal-Wallis rank sum test followed by multiple comparisons. Kruskal-Wallis chi-squared= 2847.3, df=1, p<0.0001.

p values for multi-comparison tests (Bonferroni corrected) were reported for comparisons between (P23M, P28MC, ORX) vs (P28MT, P35M, P50M) in M or L and for comparisons between M and L within each group: **p < 0.01, ***p < 0.001.

**Supplementary Fig. 9u:** Kruskal-Wallis rank sum test followed by multiple comparisons. Kruskal-Wallis chi-squared= 73.89, df=3, p<0.0001.

p values for multi-comparison tests (Bonferroni corrected) were reported for comparison between groups. ***p < 0.001.

**Supplementary Figure 10d:** One-way ANOVA followed by multiple comparisons. ANOVA, F (5, 594) = 5120.211650, p<0.0001. Tukey’s multi-comparison test: real P50 vs shuffle P50, p< 0.0001. real P50 vs real P35, p< 0.0001. real P50 vs shuffle P35, p< 0.0001. real P50 vs real P23, p< 0.0001. real P50 vs shuffle P23, p< 0.0001. shuffle P50 vs real P35, p< 0.0001. shuffle P50 vs shuffle P35, p< 0.0001. shuffle P50 vs real P23, p< 0.0001. shuffle P50 vs shuffle P23, p= 0.9968. real P35 vs shuffle P35, p< 0.0001. real P35 vs real P23, p= 0.0004. real P35 vs shuffle P23, p< 0.0001. real P23 vs shuffle P35, p< 0.0001. shuffle P35 vs shuffle P23, p< 0.0001. real P23 vs shuffle P23, p< 0.0001.

**Supplementary Fig.12c left branch1 genes:** Kruskal-Wallis rank sum test followed by multiple comparisons. Kruskal-Wallis chi-squared= 330.26, df=4, p<0.0001. p values for multi-comparison tests (Bonferroni corrected) were reported for comparison between (Esr1KOF, P23F, OVX) vs (P35F, P50F): ***p < 0.001, **p < 0.01.

**Supplementary Fig.12c left branch2 genes:** Kruskal-Wallis rank sum test followed by multiple comparisons. Kruskal-Wallis chi-squared= 26.497, df=4, p<0.0001. p values for multi-comparison tests (Bonferroni corrected) were reported for comparison between (Esr1KOF, P23F, OVX) vs (P35F, P50F). **Supplementary Fig.12c left branch common genes:** Kruskal-Wallis rank sum test followed by multiple comparisons. Kruskal-Wallis chi-squared= 333.74, df=4, p<0.0001. p values for multi-comparison tests (Bonferroni corrected) were reported for comparison between (Esr1KOF, P23F, OVX) vs (P35F, P50F): ***p < 0.001.

**Supplementary Fig.12c right branch1 genes:** Kruskal-Wallis rank sum test followed by multiple comparisons. Kruskal-Wallis chi-squared= 84.947, df=4, p<0.0001. p values for multi-comparison tests (Bonferroni corrected) were reported for comparison between (Esr1KOF, P23F, OVX) vs (P35F, P50F). **Supplementary Fig.12c right branch2 genes:** Kruskal-Wallis rank sum test followed by multiple comparisons. Kruskal-Wallis chi-squared= 319.6, df=4, p<0.0001. p values for multi-comparison tests (Bonferroni corrected) were reported for comparison between (Esr1KOF, P23F, OVX) vs (P35F, P50F): ***p < 0.001.

**Supplementary Fig.12c right branch common genes:** Kruskal-Wallis rank sum test followed by multiple comparisons. Kruskal-Wallis chi-squared= 380.94, df=4, p<0.0001. p values for multi-comparison tests (Bonferroni corrected) were reported for comparison between (Esr1KOF, P23F, OVX) vs (P35F, P50F): ***p < 0.001.

**Supplementary Fig.12d left branch1 genes:** Kruskal-Wallis rank sum test followed by multiple comparisons. Kruskal-Wallis chi-squared= 1136.2, df=4, p<0.0001. p values for multi-comparison tests (Bonferroni corrected) were reported for comparison between (Esr1KOM, P23M, ORX) vs (P35M, P50M): ***p < 0.001.

**Supplementary Fig.12d left branch2 genes:** Kruskal-Wallis rank sum test followed by multiple comparisons. Kruskal-Wallis chi-squared= 109.38, df=4, p<0.0001. p values for multi-comparison tests (Bonferroni corrected) were reported for comparison between (Esr1KOM, P23M, ORX) vs (P35M, P50M).

**Supplementary Fig.12d left branch common genes:** Kruskal-Wallis rank sum test followed by multiple comparisons. Kruskal-Wallis chi-squared= 613.41, df=4, p<0.0001. p values for multi-comparison tests (Bonferroni corrected) were reported for comparison between (Esr1KOM, P23M, ORX) vs (P35M, P50M): ***p < 0.001.

**Supplementary Fig.12d right branch1 genes:** Kruskal-Wallis rank sum test followed by multiple comparisons. Kruskal-Wallis chi-squared= 153.86, df=4, p<0.0001. p values for multi-comparison tests (Bonferroni corrected) were reported for comparison between (Esr1KOM, P23M, ORX) vs (P35M, P50M).

**Supplementary Fig.12d right branch2 genes:** Kruskal-Wallis rank sum test followed by multiple comparisons. Kruskal-Wallis chi-squared= 447.68, df=4, p<0.0001. p values for multi-comparison tests (Bonferroni corrected) were reported for comparison between (Esr1KOM, P23M, ORX) vs (P35M, P50M): ***p < 0.001.

**Supplementary Fig.12d right branch common genes:** Kruskal-Wallis rank sum test followed by multiple comparisons. Kruskal-Wallis chi-squared= 596.01, df=4, p<0.0001. p values for multi-comparison tests (Bonferroni corrected) were reported for comparison between (Esr1KOM, P23M, ORX) vs (P35M, P50M): ***p < 0.001.

**Supplementary Fig.12e branch1:** Kruskal-Wallis rank sum test followed by multiple comparisons. Kruskal-Wallis chi-squared= 1025.1, df=4, p<0.0001. p values for multi-comparison tests (Bonferroni corrected) were reported for comparison between (Esr1KOF,P23F, OVX) vs (P35F, P50F): ***p < 0.001.

**Supplementary Fig.12e branch2:** Kruskal-Wallis rank sum test followed by multiple comparisons. Kruskal-Wallis chi-squared= 1174.6, df=4, p<0.0001. p values for multi-comparison tests (Bonferroni corrected) were reported for comparison between (Esr1KOF,P23F, OVX) vs (P35F, P50F): ***p < 0.001.

**Supplementary Fig.12f branch1:** Kruskal-Wallis rank sum test followed by multiple comparisons. Kruskal-Wallis chi-squared= 1014.8, df=4, p<0.0001. p values for multi-comparison tests (Bonferroni corrected) were reported for comparison between (Esr1KOM, P23M, ORX) vs (P35M, P50M): ***p < 0.001.

**Supplementary Fig.12f branch2:** Kruskal-Wallis rank sum test followed by multiple comparisons. Kruskal-Wallis chi-squared= 1113.1, df=4, p<0.0001. p values for multi-comparison tests (Bonferroni corrected) were reported for comparison between (Esr1KOM, P23M, ORX) vs (P35M, P50M): ***p < 0.001.

## Resource and Reagent Table

| REAGENT or RESOURCE | SOURCE | IDENTIFIER |
| --- | --- | --- |
| Antibodies |  |  |
| Donkey anti-Rabbit IgG (H+L) Highly Cross-Adsorbed Secondary Antibody, Alexa Fluor 568 | Invitrogen | A10042 |
| Alexa Fluor® 647 AffiniPure Donkey Anti-Rabbit IgG (H+L) | Jackson Immuno Research | 711-605-152 |
| Anti-Estrogen Receptor alpha antibody | Santa Cruz | sc-542 |
| Bacterial and Virus Strains |  |  |
| AAV-frtFlex-Cre | Dr. Larry Zweifel laboratory | NA |
| AAV-fDIO-eYFP | UNC Vector Core | NA |
| Chemicals, Peptides, and Recombinant Proteins |  |  |
| Anisomycin from *Streptomyces griseolus* | Sigma-Aldrich | Cat#A9789 |
| Pronase | Sigma-Aldrich | Cat#10165921001 |
| Actinomycin D | Sigma-Aldrich | Cat#A1410 |
| N-Methyl-D-glucamine | Sigma-Aldrich | Cat#66930 |
| (+)-Sodium L-ascorbate | Sigma-Aldrich | Cat#A7631 |
| Sodium pyruvate | Sigma-Aldrich | Cat#P2256 |
| N-Acetyl-L-cysteine | Sigma-Aldrich | Cat#A7250 |
| Critical Commercial Assays |  |  |
| Chromium Single Cell 3' Library & Gel Bead Kit v2 | 10x Genomics | Cat# 120267 |
| Chromium i7 Multiplex Kit Kit | 10x Genomics | Cat#120262 |
| Chromium Single Cell A Chip | 10x Genomics | Cat#1000009 |
| Chromium Controller & Accessory Kit | 10x Genomics | Cat#120223 |
| Dead Cell Removal Kit | Miltenyi Biotec | Cat#130-090-101 |
| Illumina NextSeq 500 v2.5 | Illumina | Cat#20024907 |
| Illumina Hiseq 3000/4000 | Illumina | Cat#PE-410-1001 |
| RNAscope Fluorescent Multiplex Reagent Kit | ACDBio | Cat#320850 |
| HCR RNA-FISH products, V3 | Molecular Instruments | N/A |
| Deposited Data |  |  |
| Mouse scRNA-seq data | Gene Expression Omnibus | GSE172177 |
| MERFISH data | Moffitt et al., 2018 | https://datadryad.org/stash/dataset/doi:10.5061/dryad.8t8s248; deposited at DRYAD. RRID N/A. |
| Experimental Models: Organisms/Strains |  |  |
| Mouse: C57BL/6J males | The Jackson Laboratory | JAX: 000664; RRID: IMSR_JAX:000664 |
| Mouse: VgatFlp | The Jackson Laboratory | JAX: 029591; RRID:IMSR_JAX:029591 |
| Mouse: Esr1lox/lox | The Jackson Laboratory | JAX: 032173; RRID:IMSR_JAX:032173 |
| Mouse: Vglut2Flp | The Jackson Laboratory | JAX: 030212; RRID:IMSR_JAX:030212 |
| Oligonucleotides |  |  |
| RNAscope® Probes, see Table S6 |  |  |
| HCR Probes, see Table S6 |  |  |
| Software and Algorithms |  |  |
| RNAscope HiPlex Image Registration Software | ACDBio | Cat#300065 |
| ImageJ (Fiji) | Schindelin et al., 2012 | RRID: SCR_002285; http://fiji.sc |
| SCENIC (v1.1.0-01) | Aibar et al., 2017 | https://aertslab.org/#scenic RRID: not found |
| R (v3.6) | N/A | https://www.r-project.org/ RRID:SCR_001905 |
| Seurat v3 | Stuart et al., 2019 | RRID: SCR_016341; https://satijalab.org/seurat/get_started.html |
| DoubletDecon (v1.02) | DePasquale et al., 2018 | https://github.com/EDePasquale/DoubletDecon RRID: not found |
| Enrichr | Chen et al., 2013; Kuleshov et al., 2016 | http://amp.pharm.mssm.edu/Enrichr/ RRID:SCR_001575 |
| Cell Ranger | 10x Genomics | RRID: SCR_017344; https://support.10xgenomics.com/single-cell-gene-expression/software/pipelines/latest/what-is-cell-ranger |
| The Gene Ontology (GO) Project | Ashburner et al., 2000 | http://www.informatics.jax.org/mgihome/GO/project.shtml RRID:SCR_006447 |
| GraphPad Prism v9.0.2 | GraphPad Software | https://www.graphpad.com/ RRID:SCR_002798 |
| CateGORizer | Reecy et al., 2008 | https://www.animalgenome.org/tools/catego/<br>RRID:SCR_005737 |
| Monocle (V3) | Cao et al., 2019 | https://cole-trapnell-lab.github.io/monocle3/; RRID:SCR_018685 |
| LISI | Korsunsky et al., 2019 | https://github.com/immunogenomics/LISI |
| scWGCNA | Morabito et al., 2021 | https://github.com/smorabit/scWGCNA |
| MELD | Burkhardt et al., 2021 | https://github.com/KrishnaswamyLab/MELD |
| PAGA | Wolf et al., 2019 | https://github.com/theislab/paga |

---

**Supplementary Fig. 1:**
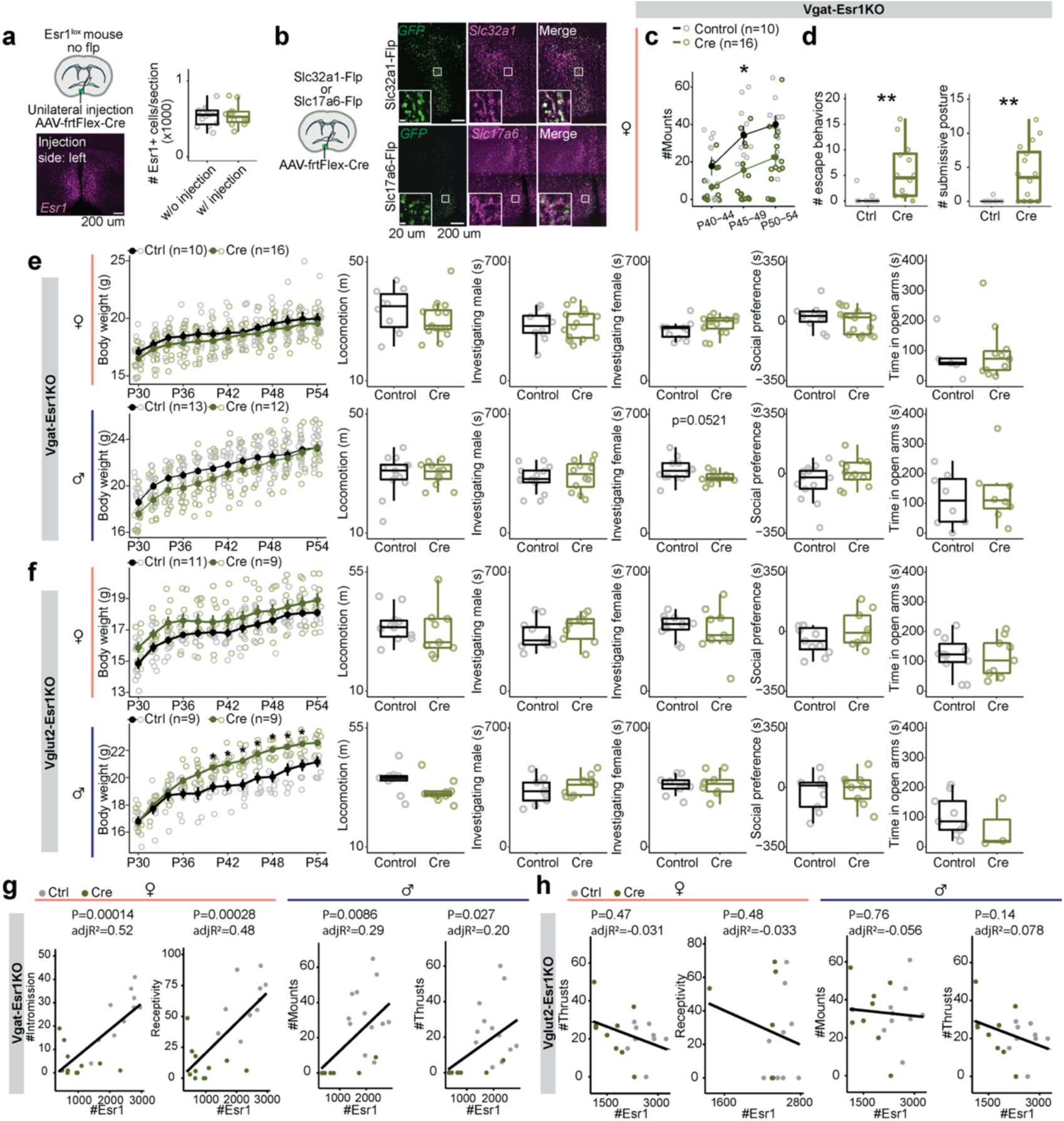
Supporting data for cell-type specific deletion of Esr1 and behavioral experiments, related to Fig. 1 **a,** Schematic illustrating unilateral injection of AAV-frtFlex-cre virus in the MPOA of Esr1^lox/lox^ mice. Representative image of Esr1 expression in MPOA after a left hemisphere injection. Quantification of Esr1 in the MPOA. Note that this is a control experiment to test viral specificity, where mice did not express Flp-recombinase. **b,** Schematic illustrating AAV-frtFlex-Cre injection in the MPOA of *Slc32a1*^Flp^ or *Slc17a6*^Flp^ mice. Representative images showing expression of *GFP* viral-reporter and *Slc32a1* (top) or *Slc17a6* (bottom) in the MPOA of *Slc32a1*^Flp^ (top) or *Slc17a6*^Flp^ (bottom) mice. **c-d,** Quantitative comparisons of female mice: number of mounts received (**c**), escape attempts (**d**, left) and submissive postures (**d**, right) in *Slc32a1*^Flp^::*Esr1*^lox/lox^ mice. **e-f,** From left to right, females (top) to males (bottom): Quantifications of body weight, locomotion, time investigating a male conspecific, time investigating a female conspecific, social preference in three chamber test, and time spent in the open arm of EPM of *Slc32a1*^Flp^::*Esr1*^lox/lox^ (**e**) or *Slc17a6*^Flp^::*Esr1*^lox/lox^ (**f**) mice. **g-h,** Linear regression analysis between number of Esr1 cells and mating-related behavioral measurements in female (left) and male (right) *Slc32a1*^Flp^::*Esr1*^lox/lox^ (**g**) or *Slc17a6*^Flp^::*Esr1*^lox/lox^ (**h**) mice. Line plots are shown in mean ± S.E.M and analyzed with a two-way repeated measures ANOVA followed by multiple comparisons. Box plots are shown with box (25%, median line, and 75%) and whiskers and analyzed with unpaired t-test. **p < 0.01. *p < 0.05. Statistical details in Methods. aR^2^: adjusted R squared; EPM: elevated plus maze.

**Supplementary Fig. 2:**
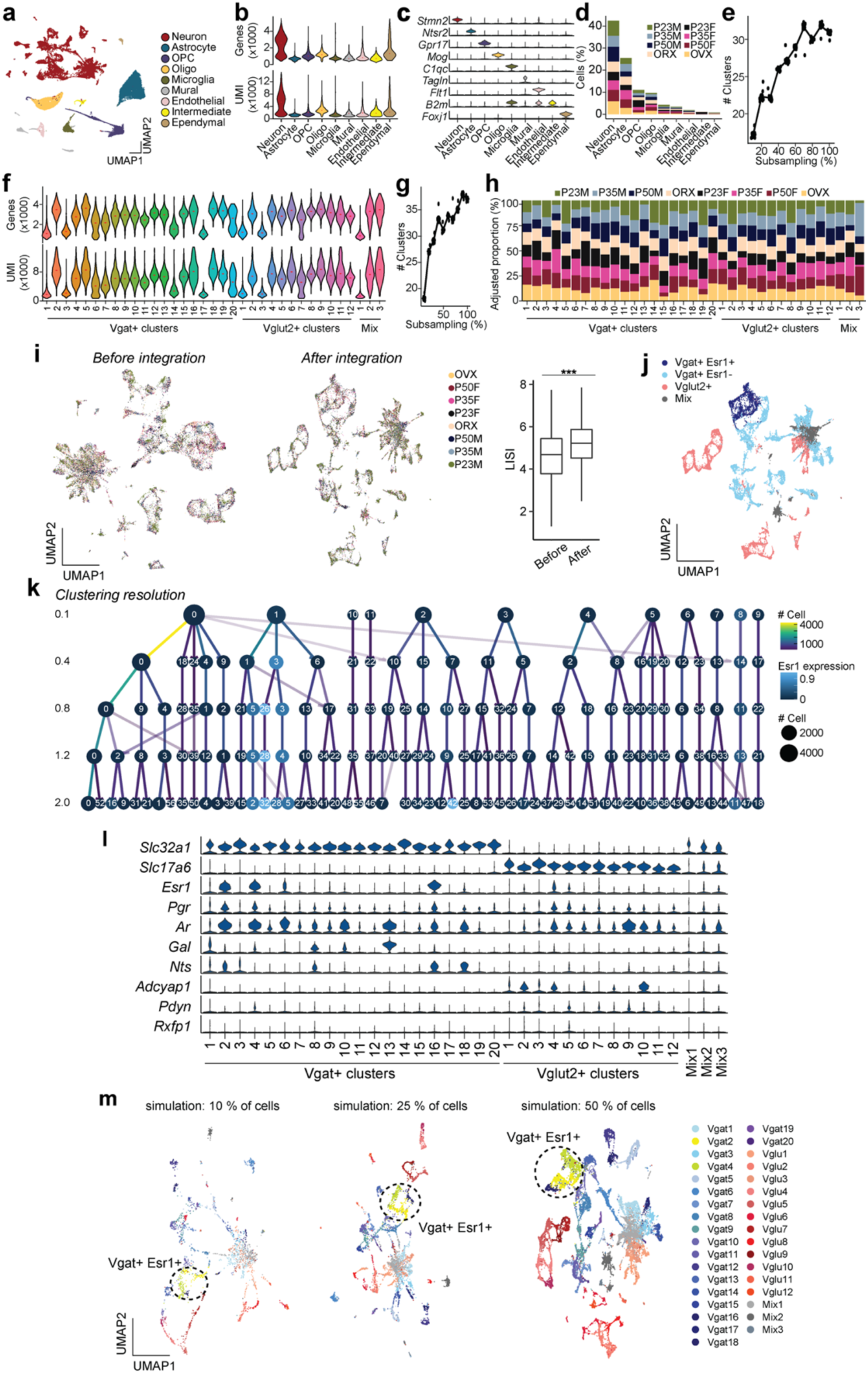
Supporting data for scRNAseq clustering, related to Fig. 2 **a,** Joint clustering of 58,921 cells from all groups (P23, P35, P50, GDX). UMAP plot is color coded by 9 cell category types, listed in legend. **b,** Violin plots showing the gene (top) and UMI (bottom) distributions in each cell category type. **c,** Violin plots of marker gene expression in each cell category type. **d,** Proportion of cells represented in each cell category type from each group. **e,** Graph representation of cluster number(s) from MPOA cell sub-sampling. **f,** Violin plots showing the gene (top) and UMI (bottom) distributions in each neuronal cell type cluster. **g,** Graph representation of cluster number(s) from MPOA neuron sub-sampling. **h,** Proportion of cells represented in each neuron cluster from each group. All neuronal cell types were present irrespective of age, sex, and hormonal states. **i,** UMAP clustering without integration (left) and then joint clustering (right) of neuronal cells from all groups. UMAP plot is color coded by experimental group. Boxplot of LISI (Local Inverse Simpson’s Index, or cell “mixing” index) was computed to assess integration performance. Note that the theoretical maximum of LISI score is 8. **j,** UMAP plot with color code for major neuronal types: Salmon = Vglut2+, Blue = Vgat+Esr1-, Purple = Vgat+Esr1+, Grey = Mixed. **k,** Cluster tree illustrating the lineage of clusters at each clustering resolution. *Esr1* expression and the number of cells in each cluster are represented by the color and the size of the dot, respectively. Number of cells diverging from a node is represented by the color of the connecting line. **l,** Violin plots showing normalized expression values of *Slc32a1*, *Slc17a6*, several steroid hormone receptor genes, and canonical marker genes in the MPOA at each cluster. **m,** *In-silico* analysis generated UMAP plots, using simulated subsets of data. UMAP plots are color coded by neuronal types. Vgat+Esr1+ clusters are highlighted. Box plots are shown with box (25%, median line, and 75%) and whiskers and analyzed with Wilcoxon rank-sum test. ***p < 0.001. Statistical details in Methods. UMIs: unique molecular identifiers; GDX: gonadectomy; OVX: ovariectomy; ORX: orchiectomy.

**Supplementary Fig. 3.**
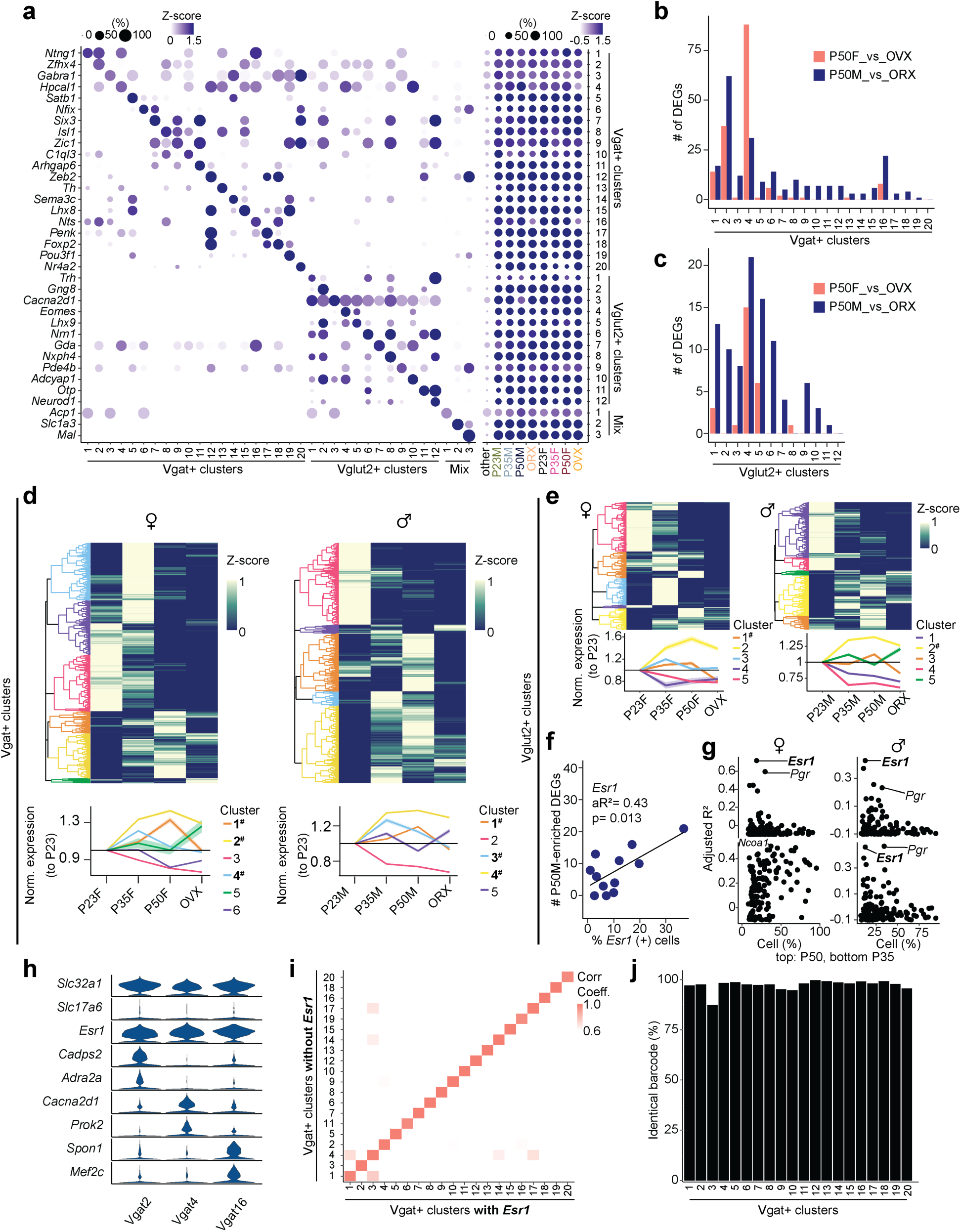
: Supporting data for scRNAseq experiments, related to Fig. 2 **a,** Dot plot illustrating scaled expression (z-score, color intensity) and percentage of expressing cells (dot size) for a list of marker genes (y-axis) in each cluster (x-axis). On the left y-axis each clusters gene expression score is presented for each group: P23, P35, P50, and GDX (females and males). **b, c,** Bar plot to show the difference in number of DEGs between P50 and GDX groups (females: salmon; males: blue) in each Vgat+ cluster (**b**) and Vglut2+ cluster (**c**). **d, e,** Heatmaps with hierarchical clustering dendrograms (top) and line graphs (bottom) to demonstrate gene expression patterns within a cluster relative to other clusters for each group: P23, P35, P50, GDX in females (left) and males (right). Clusters are denoted by color. Cluster numbers that are bolded and marked with a hashtag (#) show higher expression at P35 and P50 than P23 and GDX. Z-scored gene expression is denoted by a separate color intensity scale. Line graph data shows normalized expression of each cluster relative to P23 expression. Vgat+ clusters (**d**); Vglut2+ clusters (**e**). f, Linear regression analysis between percentage of *Esr1*-expressing cells and number of P50M-enriched DEGs compared to GDX for each Vglut2+ cluster (dot). **g,** Scatter plots showing aR^2^ values of hormone receptor genes (dots) in females (left) and males (right) comparing the percentage of percent expressing cells to the number of DEGs in comparison with GDX samples at each Vglut2+ cluster for P50-enriched (top) and P35-enriched (bottom) DEGs. **h,** Violin plots showing the normalized expression values of *Slc32a1*, *Slc17a6*, *Esr1,* and several subtype-specific genes in the MPOA for each Vgat+Esr1+ cluster (Vgat 2, 4, 16). **i,** Heatmap illustrating the Pearson correlation coefficient between Vgat+ clusters including or excluding *Esr1* expression data. Near identical clusters were detected irrespective of *Esr1* expression. **j,** Bar plot showing the percent of identical cell barcodes within each Vgat+ cluster comparing inclusion and exclusion of *Esr1* expression data. Near identical clusters were detected irrespective of *Esr1* expression. Line graphs include standard error and were analyzed using one-way repeated-measures ANOVA followed by multiple comparisons. Statistical details in Methods. GDX: gonadectomy; OVX: ovariectomy; ORX: orchiectomy; aR^2^: adjusted R squared; DEG: differentially expressed gene.

**Supplementary Fig. 4:**
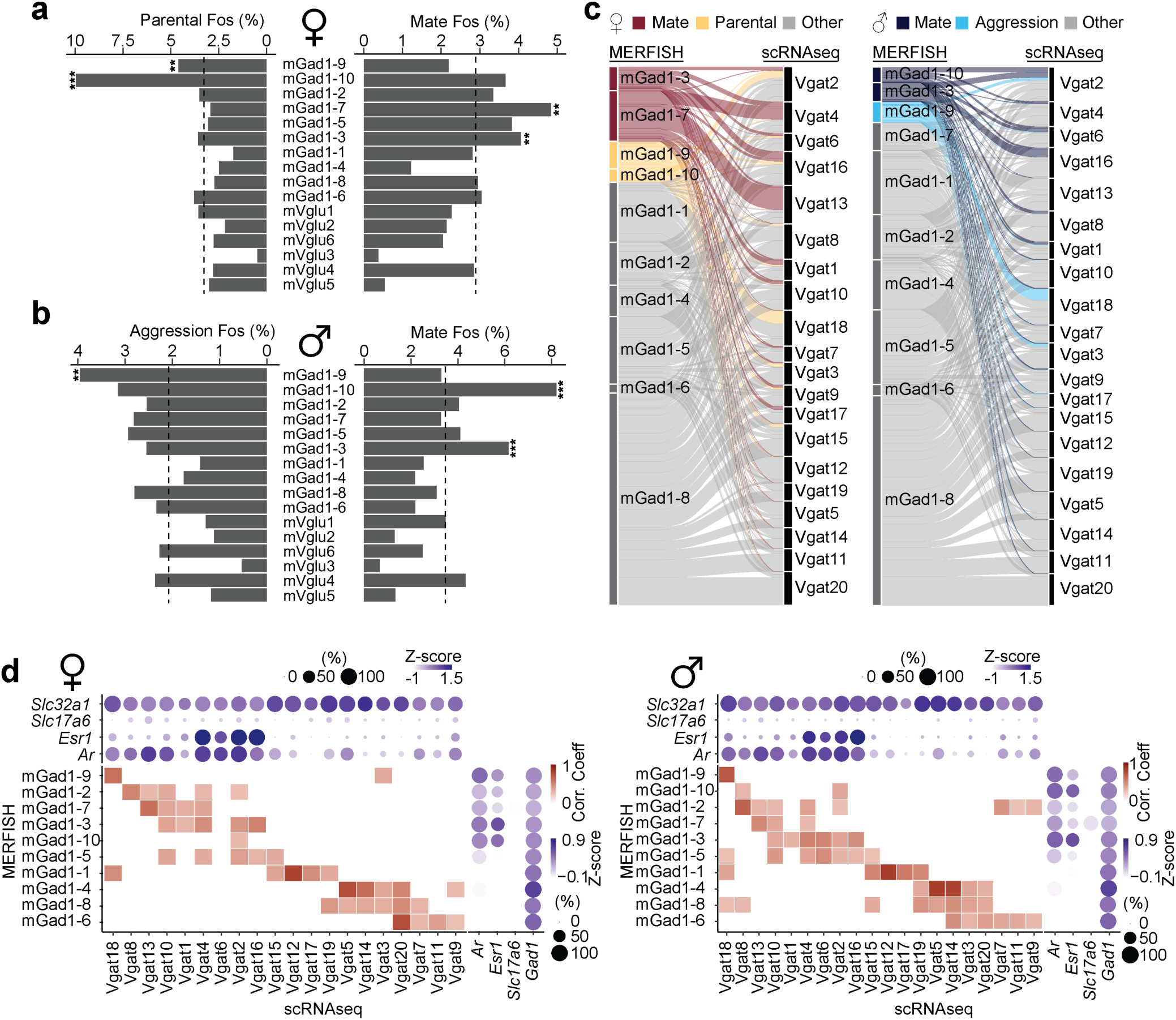
Supporting data for scRNAseq and MERFISH analysis, related to Fig. 2 **a, b,** Percentage of *Fos* positive cells associated with female parental and mating behaviors (**a**) and male aggression and mating behaviors (**b**) from MERFISH cell clusters (Moffitt et al., 2018). Dashed lines indicate the overall mean percentage. Each cluster was compared to the mean using a Fisher’s exact test. **p < 0.01, ***p < 0.001. Statistical details in Methods. **c**, Alluvial plots showing the correspondence between MERFISH and scRNAseq clusters color coded by behavior. **d,** Heatmaps and dot plots showing both Pearson correlation coefficients of MERFISH + scRNAseq GABAergic (mGad; Vgat) clusters and their expression of selected genes: *Slc32a1*, *Slc17a6*, *Esr1*, *Ar,* or *Gad1* in the scRNAseq dataset (top) and the MERFISH dataset (right). Increased color intensity corresponds to greater values. Dot size represents percentage of cells expressing the corresponding gene. Left: females; right: males.

**Supplementary Fig. 5:**
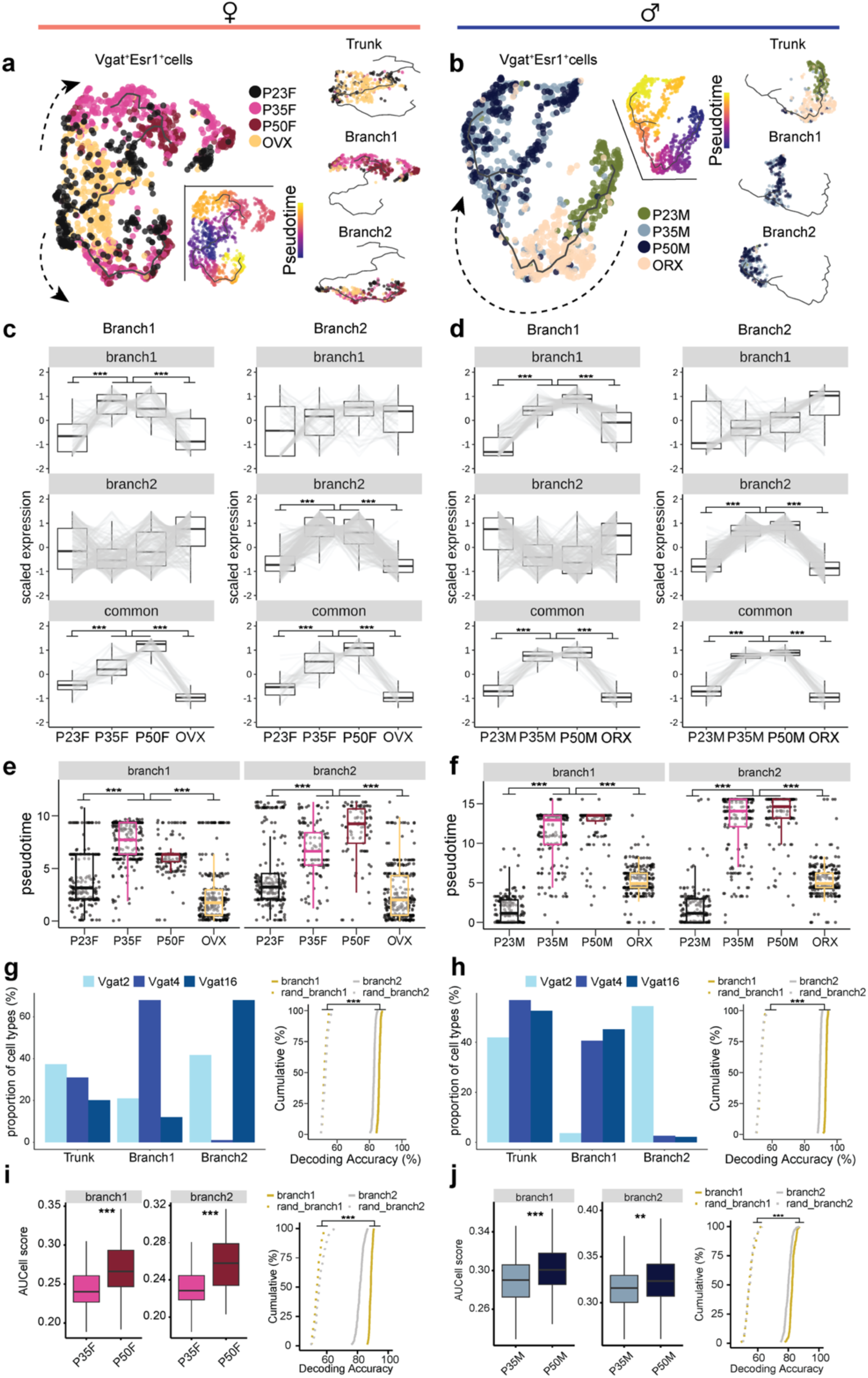
Supporting data for transcriptional trajectory branches from scRNAseq data, related to Fig. 3 **a, b,** UMAP visualization of Vgat+Esr1+ cells and their transcriptional trajectories depicted by a solid black line. Vgat+Esr1+ cells are color coded by group (left) and pseudotime (right), where progression of time is delineated from dark to bright coloring. Dashed arrows indicate the direction of transcriptional progression. Smaller UMAPs on the right visually isolate the trunk and two identified branch trajectories. **a**: females; **b**: males. **c, d,** Box plots show scaled gene expression in Vgat+Esr1+ cells from Branch 1 (left column) and Branch 2 (right column) trajectories for Branch 1-specific genes (top row), Branch 2-specific genes (middle row), and Branch-common genes (bottom row) across all groups. **e, f,** Box plots show pseudotime score for Branch 1 (left column) and Branch 2 (right column) trajectories across groups in females (**e**) and males (**f**). **g, h,** Bar plots represent the percentage of Vgat+Esr1+ cells that belong to cluster Vgat 2, 4, and 16 in the trunk and branches. Cumulative distributions for decoding accuracy between more mature (P35, P50) and immature (P23, GDX) groups using Branch 1 (yellow) or Branch 2 (grey) -specific genes compared to shuffled data (dashed line). **g:** females; **h:** males. **i, j,** Box plots of HA-DEG aggregate expression (AUCell score) separated by branch in P35 and P50. Cumulative distributions for decoding accuracy between P35 and using Branch 1 (yellow) or Branch 2 (grey) -specific genes compared to shuffled data (dashed line). **i:** females; **j:** males. Box plots are shown with box (25%, median line, and 75%) and whiskers and analyzed using Kruskal-Wallis H test followed by multiple comparisons test. p-values were Bonferroni corrected. Cumulative distributions were analyzed with unpaired t-tests for each branch. **p < 0.01, ***p < 0.001. Statistical details in Methods. HA-DEG: hormone-associated differentially expressed gene; GDX: gonadectomy; OVX: ovariectomy; ORX: orchiectomy.

**Supplementary Fig. 6:**
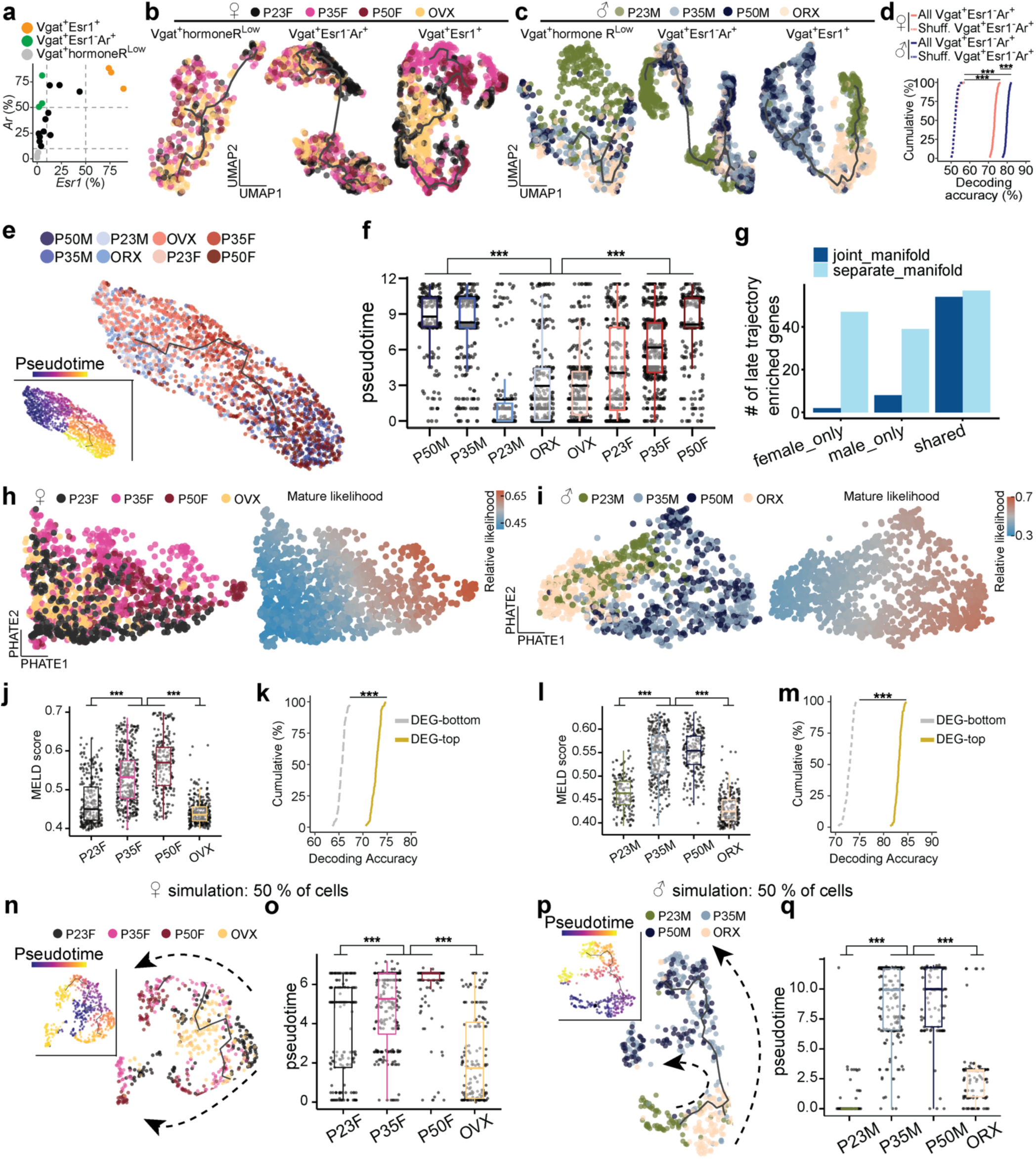
Supporting data for scRNAseq trajectory analysis, related to Fig. 3 **a,** Scatter plot showing the percentage of *Esr1* and/or *Ar* expressing cells in each Vgat+ cluster (dot). **b,c,** UMAP visualization of transcriptional trajectories (black line) color coded by experimental group in Vgat+hormoneR^Low^ (left), Vgat+Esr1-Ar+ (middle), and Vgat+Esr1+ (right) in females (**b**) and males (**c**). **d,** Cumulative distributions of decoding accuracy by SVM classification between mature groups (P50, P35) and immature groups (P23, GDX) using expression data from Vgat+Esr1-Ar+ (salmon: female, blue: male, solid line), or shuffled data (dashed line). **e,** UMAP visualization of Vgat+Esr1+ cells combining males and females, depicting their transcriptional trajectory denoted by a solid black line. Vgat+Esr1+ cells are color coded by group (right) and pseudotime (left), where progression of time is delineated from dark to bright coloring. **f,** Box plot of pseudotime score from combined male and female UMAP (**e**) separated by group. **g,** Bar plot showing the numbers of sex-specific and shared genes enriched in later trajectory, which were identified using a separate manifold for each sex or a joint manifold. Majority of sex-specific gene programs were not identified in joint manifold analysis. **h,i,** PHATE visualization of transcriptional progression in Vgat+Esr1+ cells (dot) color coded by group (left) and relative mature likelihood (right) in females (**h**) and males (**i**). **j, l,** Box plot showing MELD score (mature likelihood) of Vgat+Esr1+ cells in females (**j**) and males (**l**). **k,m,** Cumulative distributions of decoding accuracy between mature groups (P50, P35) and immature groups (P23, GDX) using highest expressing (yellow, used in the HCR experiments) or lowest expressing (grey) HA-DEGs in females (**k**) and males (**m**). **n,p,** *In silico* simulation using a 50% subset of the data. UMAP visualization of transcriptional trajectory (black line) and cells (dots) color coded by group (right) and pseudotime (left) in female (**n**) and male (**p**) Vgat+Esr1+ cells. Dashed arrows indicate the direction of transcriptional progression. **o, q,** *In silico* simulation analysis using a 50% subset of the data. Box plots show pseudotime values assigned to each Vgat+Esr1+ cell across groups in females (**o**) and males (**q**). **o,** UMAP visualization of transcriptional trajectory (black line) and cells (dots) color-coded by group (right) or pseudotime (left) in Vgat+Esr1+populations, including both sexes. Box plots are shown with box (25%, median line, and 75%) and whiskers and analyzed using Kruskal-Wallis H test followed by multiple comparisons test. p-values were Bonferroni corrected. Cumulative distributions were analyzed with unpaired t-tests. ***p < 0.001. Statistical details in Methods. SVM: support vector machine.

**Supplementary Fig. 7:**
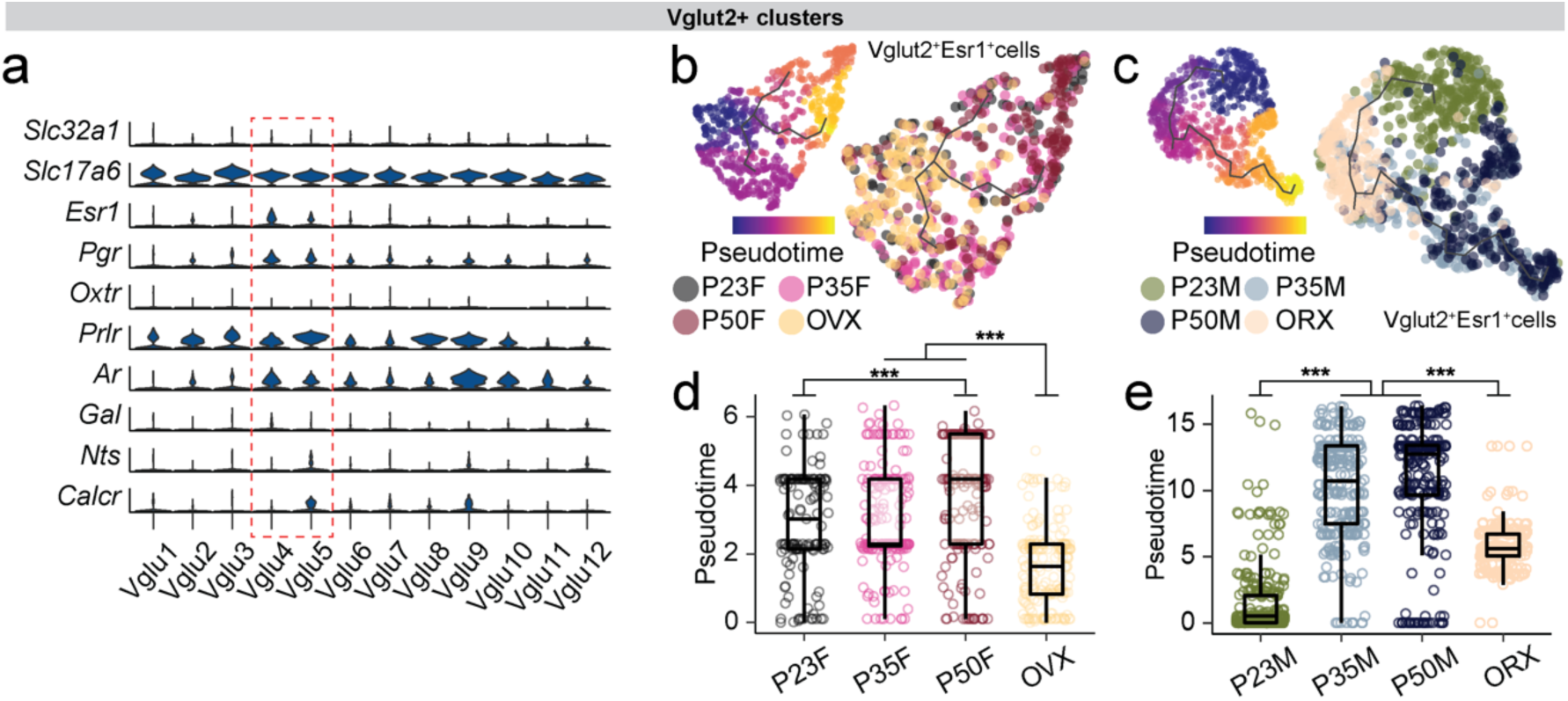
Supporting information for trajectory analysis on scRNAseq data, related to Fig. 3 **a,** Violin plots show normalized expression values of *Slc32a1*, *Slc17a6*, several steroid hormone receptor genes, and canonical marker genes in the MPOA at each Vglut2+ cluster. Vglut2+Esr1+ clusters are highlighted. **b, c,** UMAP visualization of transcriptional trajectory (black line) and cells (dots) color coded by pseudotime (left) or group (right) in female (**b**) and male (**c**) Vglut2+Esr1+ populations. **d, e,** Box plot showing pseudotime of Vglut2+Esr1+ cells in females (**d**) and males (**e**). Box plots are shown with box (25%, median line, and 75%) and whiskers and analyzed using Kruskal-Wallis H test followed by multiple comparisons test. p-values were Bonferroni corrected. ***p < 0.001. Statistical details in Methods.

**Supplementary Fig. 8:**
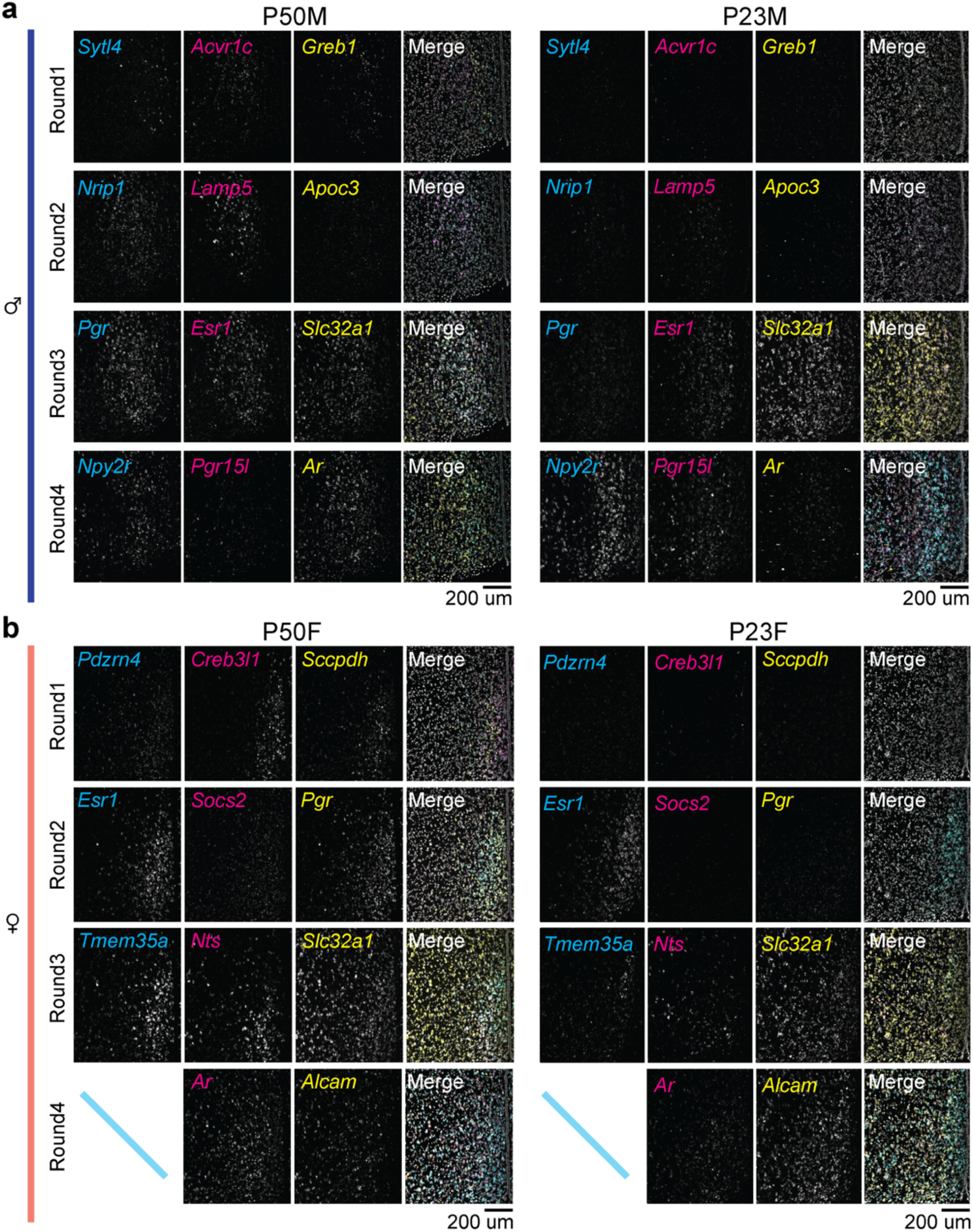
Supporting data for HM-HCR FISH, related to Fig. 4 **a, b,** Representative images showing detected genes in 4 iterative rounds of HM-HCR FISH from the MPOA at P50 (left) and P23 (right). **a**: males; **b**: females. Scale bars: 200 µm.

**Supplementary Fig. 9:**
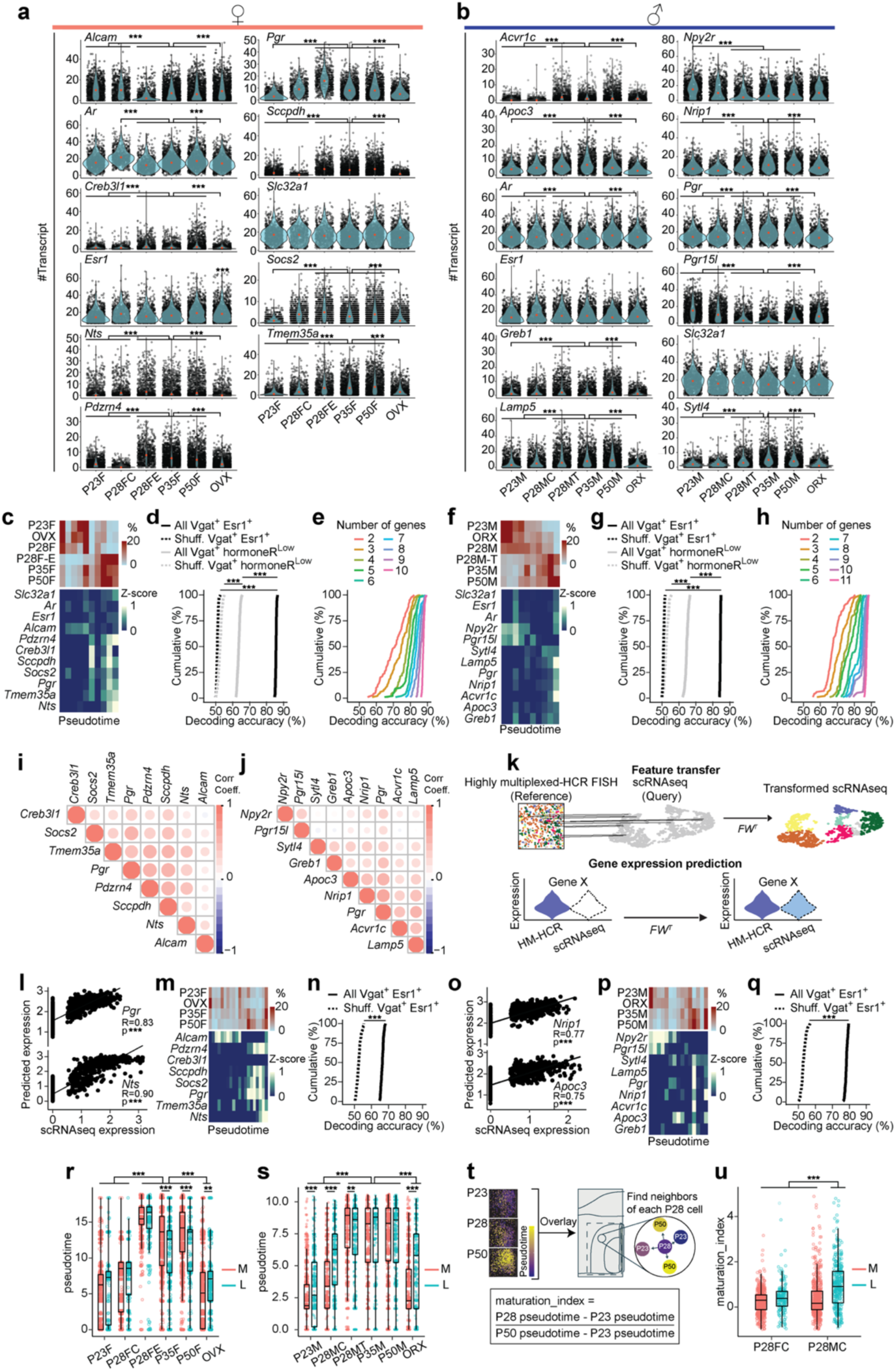
Supporting data for HM-HCR FISH experiments related to Fig. 4. **a, b,** Violin plots showing the expression values of individual genes across experimental groups in MPOA Vgat+Esr1+ cells. **a**: females; **b**: males. **c, f,** Heatmaps showing the percentage of Vgat+Esr1+ cells (top) and scaled gene expression (bottom) across pseudotime. **c**: females; **f**: males. **d, g,** Cumulative distributions of decoding accuracy between mature groups (P50, P35, hormone-supplemented P28) and immature groups (P23, GDX, P28 control) using expression data from Vgat+Esr1+ (black, solid line), Vgat+hormoneR^Low^ (grey, solid line), and shuffled data (dashed lines). **d**: females; **g**: males. **e, h,** Cumulative distributions of decoding accuracy between mature (P50, P35, hormone-supplemented P28) and immature (P23, GDX, P28 control) groups using subsets of Vgat+Esr1+ gene expression data, color coded by number of genes used for decoding. **e**: females; **h**: males. i**, j,** Correlograms of Pearson correlation coefficients between all pairs of DEGs in MPOA Vgat+Esr1+ cells. **i**: females; **j**: males. **k,** Schematic illustrating integrative analysis between HM-HCR FISH and scRNAseq datasets to predict scRNAseq gene expression. **l, o,** Scatter plots showing the correlated expressions of real and predicted scRNAseq data. **l**: females; **o**: males. **m, p,** Heatmaps showing the percentage of Vgat+Esr1+ cells (top) and scaled predicted gene expression (bottom) across pseudotime. **m**: females; **p**: males. **n, p,** Cumulative distributions of decoding accuracy between mature groups (P50, P35) and immature groups (P23, GDX) using predicted expression data in females (**n**) and males (**p**). **r-u,** Quantitative analyses of pseudotime in space, corresponding to **Fig. 4f, g**. **r, s,** Boxplots showing pseudotime of Vgat+Esr1+ cells in the medial and lateral MPOA of females (**r**) and males (**s**). **t,** Schematic illustrating the quantification of sex differences across the transcriptional progression rate. Neighborhood analysis was performed in physical space to compute the maturation index of cells from P28 control groups. **u,** Box plot of maturation index data from Vgat+Esr1+ cells in the medial (M) and lateral (L) MPOA of P28 female and male controls. At P28, male mice showed an early onset of transcriptional progression in the lateral part of the MPOA. Violin plots are outlined with a distribution line, individual dots represent each cell, and red dot indicates the median. Violin plots are analyzed with Wilcoxon rank-sum test. Box plots are shown with box (25%, median line, and 75%) and whiskers and analyzed using Kruskal-Wallis H test followed by multiple comparisons test. p-values were Bonferroni corrected. Cumulative distributions were analyzed with unpaired t-tests. ***p < 0.001. Statistical details in Methods. R: Pearson correlation coefficient; M: medial MPOA; L: lateral MPOA; P28FC: P28 female control; P28FE: P28 females with estrogen treatment; P28MC: P28 male control; P28MT: P28 males with testosterone treatment; GDX: gonadectomy; OVX: ovariectomy; ORX: orchiectomy.

**Supplementary Fig. 10.**
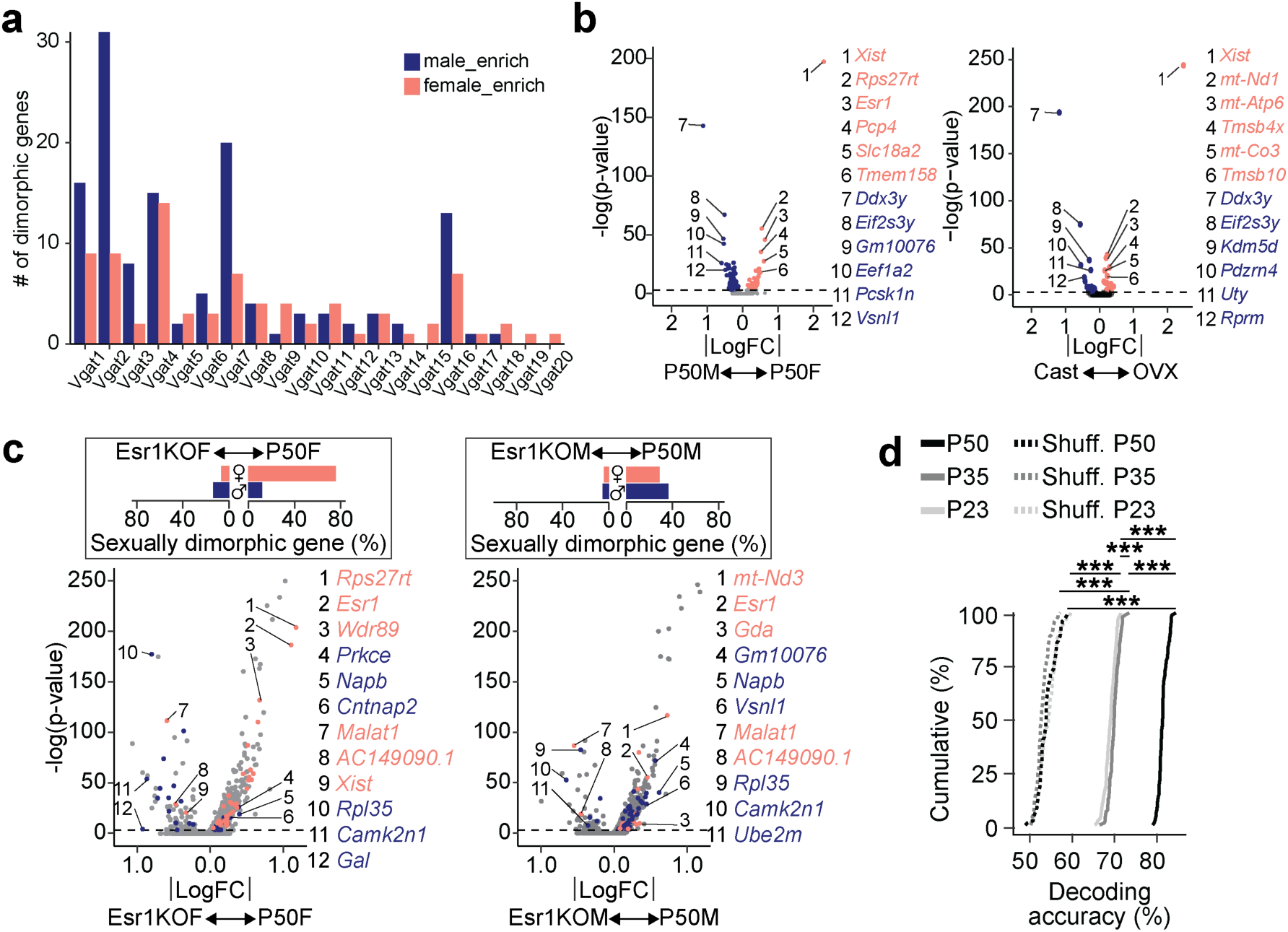
: Supporting data for sexual dimorphism analysis, related to Fig. 5-7 **a,** Bar plot showing the number of sexually dimorphic genes at each Vgat+ cluster comparing female (salmon) and male (blue) groups at P50. **b,** Volcano plot comparing P50 female (salmon) and male (blue) gene expression in Vgat+Esr1+ cells (left). Dimorphic genes are highlighted and indicated numerically. This is the same panel shown in **Fig. 5e** and reused here to facilitate the comparison with GDX data shown on the right. Volcano plot comparing OVX (salmon) and ORX (blue) gene expression in Vgat+Esr1+ cells. Dimorphic genes are highlighted and indicated numerically. **c,** Volcano plots comparing intact P50 to Esr1KO P50 gene expression in Vgat+Esr1+ cells. The percent of sexually dimorphic genes (P50F-enriched genes: salmon, P50M-enriched genes: blue) present in intact P50 and Esr1KOs are quantified in the bar plot (top). Left: females; right: males. **d,** Cumulative distributions of decoding accuracy between males and females (P50: black, P35: dark grey, P23: light grey, shuffled: dashed). Cumulative distributions were analyzed with one-way ANOVA followed by multiple comparisons. Details in Methods. ***p < 0.001. Statistical details in Methods. GDX: gonadectomy; OVX: ovariectomy; ORX: orchiectomy; LogFC: Log fold change.

**Supplementary Fig. 11:**
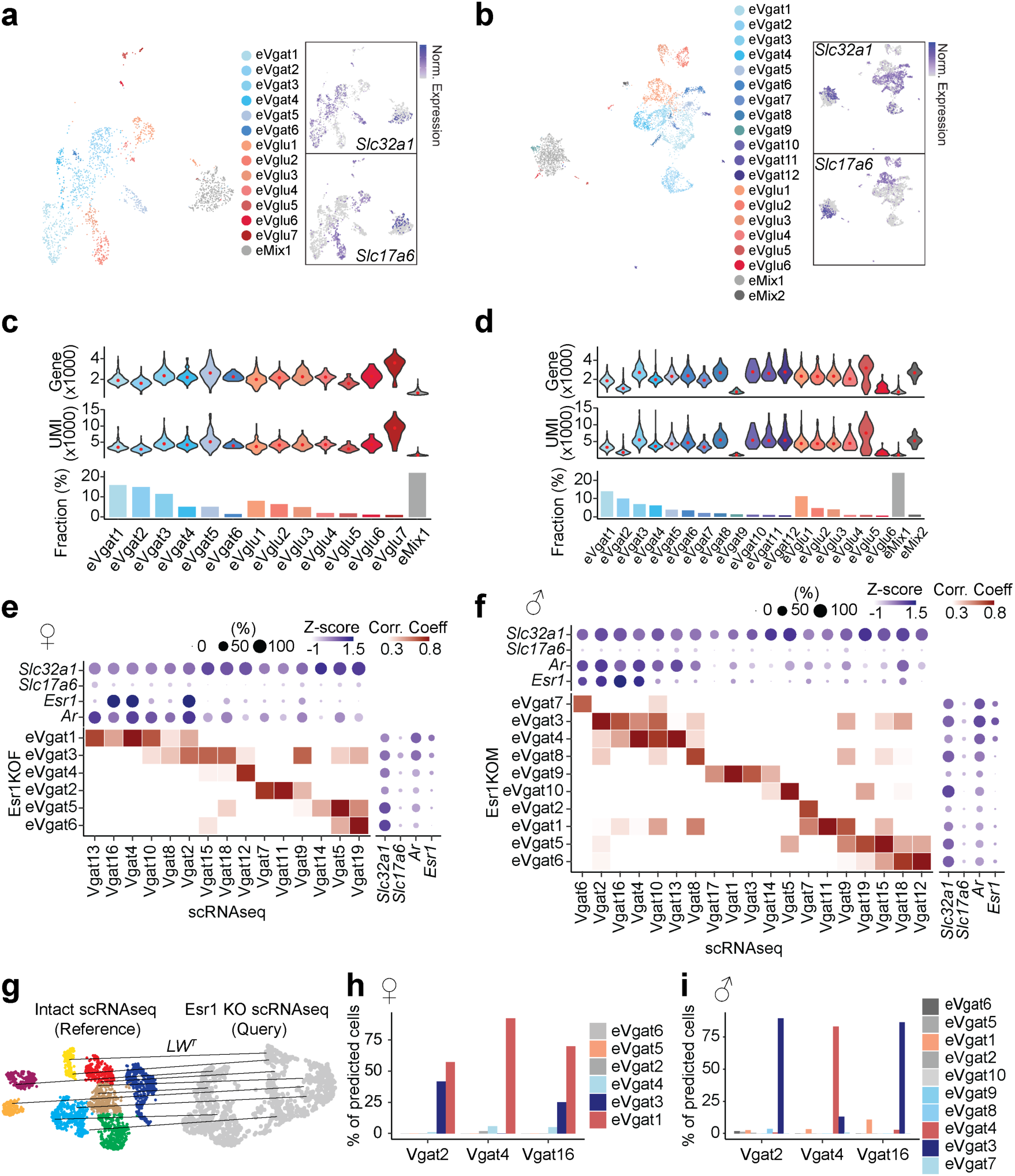
Supporting data for trajectory and gene regulatory network analysis followed by cell type specific deletion of *Esr1*, related to Fig. 7 **a, b,** UMAP visualization of neuronal clusters in Esr1KOF (**a**, left) and Esr1KOM (**b**, left). Feature plots showing normalized expression of *Slc32a1* and *Slc17a6* in Esr1KOF (**a**, right) and Esr1KOM (**b**, right). **c, d,** Violin plots showing gene (top) and UMI (middle) distributions in each neuronal cell type. Bar graphs show the fraction representation of each neuronal cluster in Esr1KOF (**c**, bottom) and Esr1KOM (**d**, bottom). **e, f,** Heatmaps illustrating Pearson correlation coefficient (red) between Vgat+ clusters of Esr1KO and intact groups. Dot plots are attached illustrating scaled expression (color intensity) and the percentage of expressing cells (dot size) of *Slc32a1*, *Slc17a6*, *Esr1,* and *Ar* in intact (top) and Esr1KOs (right side) in females (**e**) and males (**f**). **g,** Schematic illustrating integrative analysis to establish correspondence between cell types of Esr1KO data and the reference scRNAseq data. **h, i,** Bar graphs showing the correspondence between Esr1KO and the reference Vgat+Esr1+ clusters in females (**h**) and males (**i**) using the label transfer technique (details in Methods). Violin plots are outlined with a distribution line and red dot indicates the median.

**Supplementary Fig. 12:**
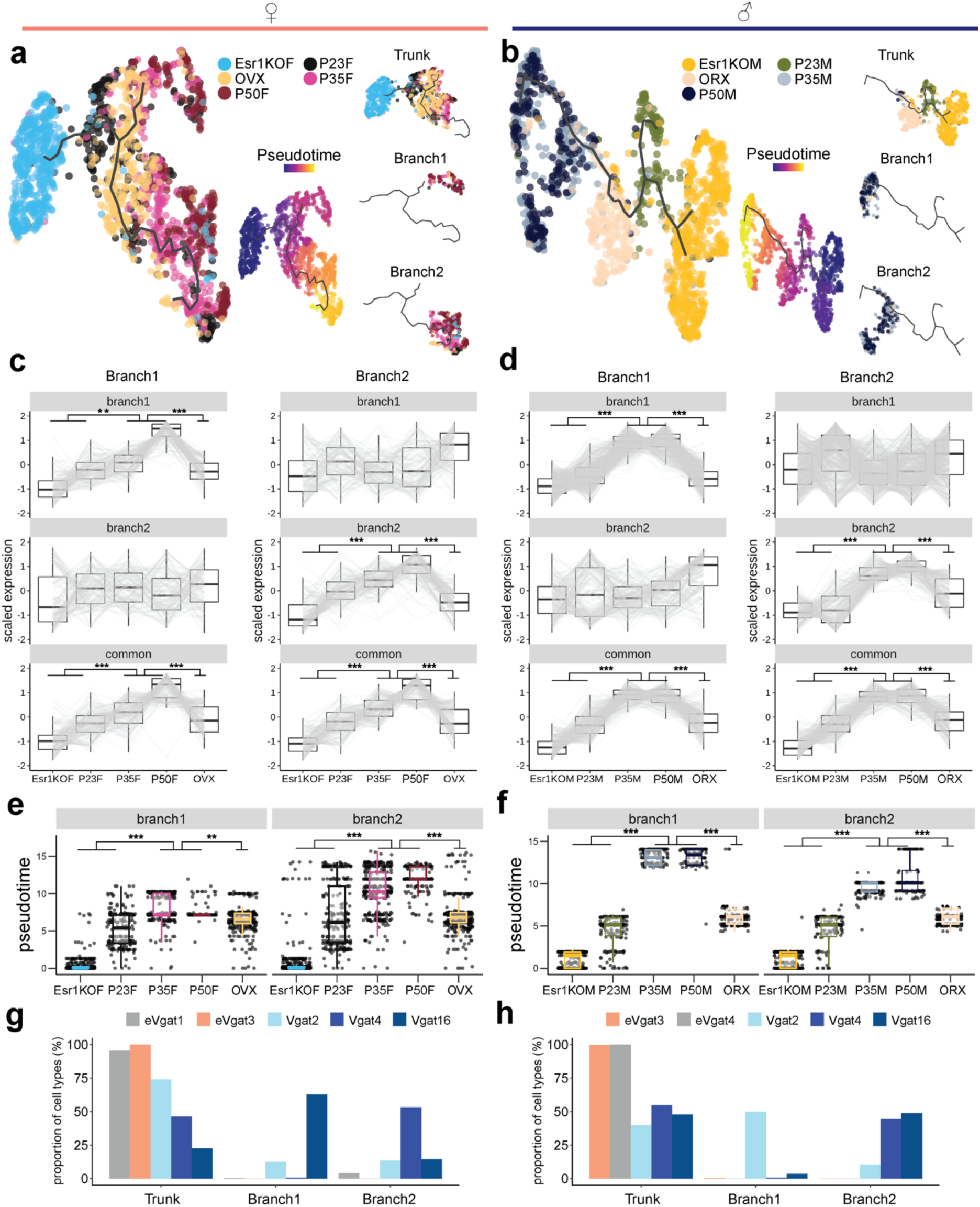
Supporting data for trajectory analysis followed by cell type specific deletion of *Esr1*, related to Fig. 7 **a,b,** UMAP visualization of transcriptional trajectories (black line) and cells (dots) color coded by experimental group (left) and pseudotime (inset) in Vgat+Esr1+ cells. UMAPs on the right visualize where cells from the branches and trunk lie on the trajectories. **a:** females; **b:** males. **c, d,** Box plots show scaled gene expression in Vgat+Esr1+ cells from Branch 1 (left column) and Branch 2 (right column) trajectories for Branch 1-specific genes (top row), Branch 2-specific genes (middle row), and Branch-common genes (bottom row) across all groups. **c:** females; **d:** males. **e, f,** Box plots show pseudotime score for Branch 1 (left column) and Branch 2 (right column) trajectories across groups in females (**e**) and males (**f**). **g,h,** Bar plot of Vgat+Esr1+ cluster (eVgat for Esr1KOs) proportion of cells that reside in the trunk, branch1, or branch2. **g:** females; **h:** males. Box plots are shown with box (25%, median line, and 75%) and whiskers and analyzed using Kruskal-Wallis H test followed by multiple comparisons test. p-values were Bonferroni corrected. ***p < 0.001, **p < 0.01. Statistical details in Methods. GDX: gonadectomy; OVX: ovariectomy; ORX: orchiectomy; KO: knockout.

**Supplementary Fig. 13:**
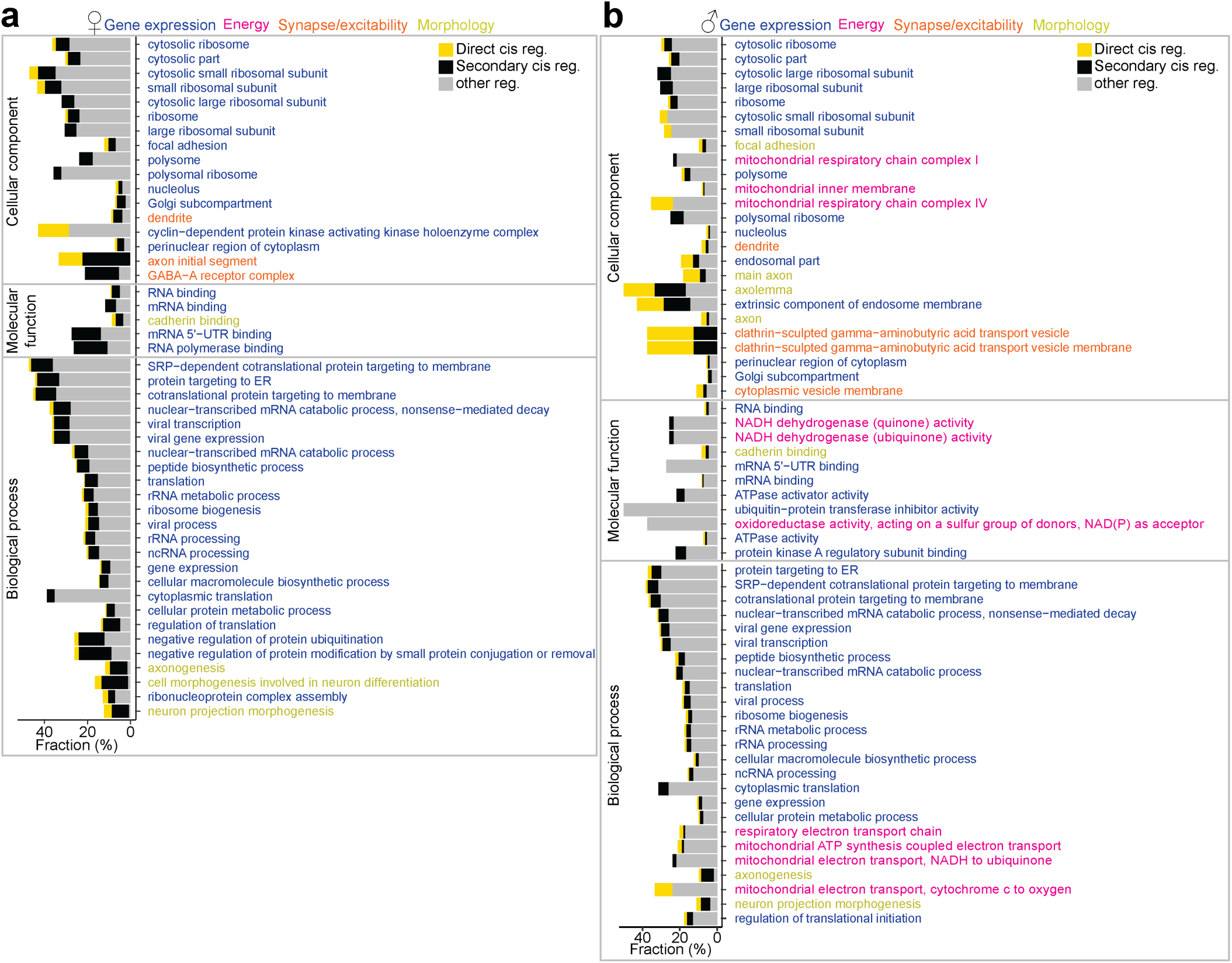
Supporting information for gene regulatory network analysis followed by cell-type specific deletion of Esr1, related to Fig. 8 **a, b,** Enriched Esr1-DEG gene ontology terms are placed alongside bar plots corresponding to gene expression percentage. Top 25 GO terms are shown and color-coded by gene category. Bars are fractionated into direct *Esr1*-regulated genes (yellow), secondary TF-regulated genes (black), and other genes regulated by Esr1-TFs and other TFs (grey). GO Biological Process, GO Molecular Function, and GO Cellular Component are referenced. **a:** females; **b:** males.

**Supplementary Table 1.** Data table showing quality of sequenced libraries and corresponding scRNAseq data from samples used in this study.

**Supplementary Table 2.** Data table showing the marker genes of neuronal clusters, related to **Fig. 2**. **Supplementary Table 3.** Data table showing the DEGs in the pseudobulk populations between groups, related to **Supplementary Fig. 3**.

**Supplementary Table 4.** Data table showing the HA-DEGs in each Vgat+ cluster, related to Supplementary Fig. 3.

**Supplementary Table 5.** Data table showing the HA-DEGs in the Vgat+Esr1+ population, related to Supplementary Fig. 3.

**Supplementary Table 6.** Data table showing co-expression scores between transcription factors and adolescent gene programs or sexually dimorphic programs related to **Fig. 2 and 5**.

**Supplementary Table 7.** Data table showing the marker genes in the clusters associated with mating, related to **Fig. 2 and Supplementary Fig. 4**.

**Supplementary Table 8.** Data table showing the genes enriched in later trajectories or branch-DEGs in the Vgat+Esr1+ population, related to **Fig. 3 and Supplementary Fig. 5**.

**Supplementary Table 9.** Data table showing the Esr1-DEGs in the Vgat+Esr1+ or Vgat+/HormoneR^low^ populations, related to **Fig. 7**.

**Supplementary Table 10.** Data table showing the genes enriched in later trajectories or branch-DEGs in the Vgat+Esr1+ population, related to **Fig. 7** and **Supplementary Fig. 12**.

**Supplementary Table 11.** Data table showing the sexually dimorphic genes in the Vgat+Esr1+ population, related to **Fig. 7**.

**Supplementary Table 12.** *In situ* hybridization probe list.

## References

(1) Blakemore, S.-J.; Burnett, S.; Dahl, R. E. The Role of Puberty in the Developing Adolescent Brain. Human Brain Mapping 2010, 31 (6), 926–933. 10.1002/hbm.21052.

(2) McCarthy, M. M.; Arnold, A. P. Reframing Sexual Differentiation of the Brain. Nat Neurosci 2011, 14 (6), 677–683. 10.1038/nn.2834.

(3) Romeo, R. D.; Richardson, H. N.; Sisk, C. L. Puberty and the Maturation of the Male Brain and Sexual Behavior: Recasting a Behavioral Potential. Neuroscience & Biobehavioral Reviews 2002, 26 (3), 381–391. 10.1016/S0149-7634(02)00009-X.

(4) Schulz, K. M.; Sisk, C. L. Pubertal Hormones, the Adolescent Brain, and the Maturation of Social Behaviors: Lessons from the Syrian Hamster. Molecular and Cellular Endocrinology 2006, 254–255, 120–126. 10.1016/j.mce.2006.04.025.

(5) Juntti, S. A.; Tollkuhn, J.; Wu, M. V.; Fraser, E. J.; Soderborg, T.; Tan, S.; Honda, S.-I.; Harada, N.; Shah, N. M. The Androgen Receptor Governs the Execution, but Not Programming, of Male Sexual and Territorial Behaviors. Neuron 2010, 66 (2), 260–272. 10.1016/j.neuron.2010.03.024.

(6) Wu, M. V.; Manoli, D. S.; Fraser, E. J.; Coats, J. K.; Tollkuhn, J.; Honda, S.-I.; Harada, N.; Shah,

7. N. M. Estrogen Masculinizes Neural Pathways and Sex-Specific Behaviors. Cell 2009, 139 (1), 61–72. 10.1016/j.cell.2009.07.036.

(7) Wu, M. V.; Tollkuhn, J. Estrogen Receptor Alpha Is Required in GABAergic, but Not Glutamatergic, Neurons to Masculinize Behavior. Hormones and Behavior 2017, 95, 3–12. 10.1016/j.yhbeh.2017.07.001.

(8) Hou, H.; Uusküla-Reimand, L.; Makarem, M.; Corre, C.; Saleh, S.; Metcalf, A.; Goldenberg, A.; Palmert, M. R.; Wilson, M. D. Gene Expression Profiling of Puberty-Associated Genes Reveals Abundant Tissue and Sex-Specific Changes across Postnatal Development. Hum Mol Genet 2017, 26 (18), 3585–3599. 10.1093/hmg/ddx246.

(9) Guo, J.; Nie, X.; Giebler, M.; Mlcochova, H.; Wang, Y.; Grow, E. J.; Kim, R.; Tharmalingam, M.; Matilionyte, G.; Lindskog, C.; Carrell, D. T.; Mitchell, R. T.; Goriely, A.; Hotaling, J. M.; Cairns, B. R. The Dynamic Transcriptional Cell Atlas of Testis Development during Human Puberty. Cell Stem Cell 2020, 26 (2), 262–276.e4. 10.1016/j.stem.2019.12.005.

(10) Clemens, L. G.; Yang, L.-Y. MPOA Lesions Affect Female Pacing of Copulation in Rats. Behavioral Neuroscience 2000, 114 (6), 1191–1202. 10.1037/0735-7044.114.6.1191.

(11) Heimer, L.; Larsson, K. Impairment of Mating Behavior in Male Rats Following Lesions in the Preoptic-Anterior Hypothalamic Continuum. Brain Research 1967, 3 (3), 248–263. 10.1016/0006-8993(67)90076-5.

(12) Oomura, Y.; Aou, S.; Koyama, Y.; Fujita, I.; Yoshimatsu, H. Central Control of Sexual Behavior. Brain Research Bulletin 1988, 20 (6), 863–870. 10.1016/0361-9230(88)90103-7.

(13) Rosenblatt, J. S.; Ceus, K. Estrogen Implants in the Medial Preoptic Area Stimulate Maternal Behavior in Male Rats. Horm Behav 1998, 33 (1), 23–30. 10.1006/hbeh.1997.1430.

(14) Sano, K.; Nakata, M.; Musatov, S.; Morishita, M.; Sakamoto, T.; Tsukahara, S.; Ogawa, S. Pubertal Activation of Estrogen Receptor α in the Medial Amygdala Is Essential for the Full Expression of Male Social Behavior in Mice. PNAS 2016, 113 (27), 7632–7637. 10.1073/pnas.1524907113.

(15) Sano, K.; Tsuda, M. C.; Musatov, S.; Sakamoto, T.; Ogawa, S. Differential Effects of Site-Specific Knockdown of Estrogen Receptor α in the Medial Amygdala, Medial Pre-Optic Area, and Ventromedial Nucleus of the Hypothalamus on Sexual and Aggressive Behavior of Male Mice. European Journal of Neuroscience 2013, 37 (8), 1308–1319. 10.1111/ejn.12131.

(16) Wei, Y.-C.; Wang, S.-R.; Jiao, Z.-L.; Zhang, W.; Lin, J.-K.; Li, X.-Y.; Li, S.-S.; Zhang, X.; Xu, X.-H. Medial Preoptic Area in Mice Is Capable of Mediating Sexually Dimorphic Behaviors Regardless of Gender. Nature Communications 2018, 9 (1), 1–15. 10.1038/s41467-017-02648-0.

(17) Fang, Y.-Y.; Yamaguchi, T.; Song, S. C.; Tritsch, N. X.; Lin, D. A Hypothalamic Midbrain Pathway Essential for Driving Maternal Behaviors. Neuron 2018, 98 (1), 192–207.e10. 10.1016/j.neuron.2018.02.019.

(18) Karigo, T.; Kennedy, A.; Yang, B.; Liu, M.; Tai, D.; Wahle, I. A.; Anderson, D. J. Distinct Hypothalamic Control of Same- and Opposite-Sex Mounting Behaviour in Mice. Nature 2020, 1–6. 10.1038/s41586-020-2995-0.

(19) McHenry, J. A.; Otis, J. M.; Rossi, M. A.; Robinson, J. E.; Kosyk, O.; Miller, N. W.; McElligott, Z. A.; Budygin, E. A.; Rubinow, D. R.; Stuber, G. D. Hormonal Gain Control of a Medial Preoptic Area Social Reward Circuit. Nature Neuroscience 2017, 20 (3), 449–458. 10.1038/nn.4487.

(20) Wu, Z.; Autry, A. E.; Bergan, J. F.; Watabe-Uchida, M.; Dulac, C. G. Galanin Neurons in the Medial Preoptic Area Govern Parental Behaviour. Nature 2014, 509 (7500), 325–330. 10.1038/nature13307.

(21) Brock, O.; Mees, C. D.; Bakker, J. Hypothalamic Expression of Oestrogen Receptor α and Androgen Receptor Is Sex-, Age- and Region-Dependent in Mice. Journal of Neuroendocrinology 2015, 27 (4), 264–276. 10.1111/jne.12258.

(22) Moffitt, J. R.; Bambah-Mukku, D.; Eichhorn, S. W.; Vaughn, E.; Shekhar, K.; Perez, J. D.; Rubinstein, N. D.; Hao, J.; Regev, A.; Dulac, C.; Zhuang, X. Molecular, Spatial, and Functional Single-Cell Profiling of the Hypothalamic Preoptic Region. Science 2018, 362 (6416). 10.1126/science.aau5324.

(23) Gegenhuber, B.; Wu, M. V.; Bronstein, R.; Tollkuhn, J. Gene Regulation by Gonadal Hormone Receptors Underlies Brain Sex Differences. Nature 2022, 606 (7912), 153–159. 10.1038/s41586-022-04686-1.

(24) Kim, D.-W.; Yao, Z.; Graybuck, L. T.; Kim, T. K.; Nguyen, T. N.; Smith, K. A.; Fong, O.; Yi, L.; Koulena, N.; Pierson, N.; Shah, S.; Lo, L.; Pool, A.-H.; Oka, Y.; Pachter, L.; Cai, L.; Tasic, B.; Zeng, H.; Anderson, D. J. Multimodal Analysis of Cell Types in a Hypothalamic Node Controlling Social Behavior. Cell 2019, 179 (3), 713–728.e17. 10.1016/j.cell.2019.09.020.

(25) Welch, J. D.; Kozareva, V.; Ferreira, A.; Vanderburg, C.; Martin, C.; Macosko, E. Z. Single-Cell Multi-Omic Integration Compares and Contrasts Features of Brain Cell Identity. Cell 2019, 177 (7), 1873–1887.e17. 10.1016/j.cell.2019.05.006.

(26) Ogawa, S.; Lubahn, D. B.; Korach, K. S.; Pfaff, D. W. Behavioral Effects of Estrogen Receptor Gene Disruption in Male Mice. PNAS 1997, 94 (4), 1476–1481. 10.1073/pnas.94.4.1476.

(27) Cheong, R. Y.; Czieselsky, K.; Porteous, R.; Herbison, A. E. Expression of ESR1 in Glutamatergic and GABAergic Neurons Is Essential for Normal Puberty Onset, Estrogen Feedback, and Fertility in Female Mice. J. Neurosci. 2015, 35 (43), 14533–14543. 10.1523/JNEUROSCI.1776-15.2015.

(28) Karigo, T.; Deutsch, D. Flexibility of Neural Circuits Regulating Mating Behaviors in Mice and Flies. Front. Neural Circuits 2022, 16. 10.3389/fncir.2022.949781.

(29) Hrvatin, S.; Sun, S.; Wilcox, O. F.; Yao, H.; Lavin-Peter, A. J.; Cicconet, M.; Assad, E. G.; Palmer, M. E.; Aronson, S.; Banks, A. S.; Griffith, E. C.; Greenberg, M. E. Neurons That Regulate Mouse Torpor. Nature 2020, 583 (7814), 115–121. 10.1038/s41586-020-2387-5.

(30) Wolf, F. A.; Hamey, F. K.; Plass, M.; Solana, J.; Dahlin, J. S.; Göttgens, B.; Rajewsky, N.; Simon, L.; Theis, F. J. PAGA: Graph Abstraction Reconciles Clustering with Trajectory Inference through a Topology Preserving Map of Single Cells. Genome Biology 2019, 20 (1), 59. 10.1186/s13059-019-1663-x.

(31) Aibar, S.; González-Blas, C. B.; Moerman, T.; Huynh-Thu, V. A.; Imrichova, H.; Hulselmans, G.; Rambow, F.; Marine, J.-C.; Geurts, P.; Aerts, J.; Oord, J.; Atak, Z. K.; Wouters, J.; Aerts, S. SCENIC: Single-Cell Regulatory Network Inference and Clustering. Nat Methods 2017, 14 (11), 1083–1086. 10.1038/nmeth.4463.

(32) Stuart, T.; Butler, A.; Hoffman, P.; Hafemeister, C.; Papalexi, E.; Mauck, W. M.; Hao, Y.; Stoeckius, M.; Smibert, P.; Satija, R. Comprehensive Integration of Single-Cell Data. Cell 2019, 177 (7), 1888–1902.e21. 10.1016/j.cell.2019.05.031.

(33) Heumos, L.; Schaar, A. C.; Lance, C.; Litinetskaya, A.; Drost, F.; Zappia, L.; Lücken, M. D.; Strobl, D. C.; Henao, J.; Curion, F.; Schiller, H. B.; Theis, F. J. Best Practices for Single-Cell Analysis across Modalities. Nat Rev Genet 2023, 24 (8), 550–572. 10.1038/s41576-023-00586-w.

(34) Cao, J.; Spielmann, M.; Qiu, X.; Huang, X.; Ibrahim, D. M.; Hill, A. J.; Zhang, F.; Mundlos, S.; Christiansen, L.; Steemers, F. J.; Trapnell, C.; Shendure, J. The Single-Cell Transcriptional Landscape of Mammalian Organogenesis. Nature 2019, 566 (7745), 496–502. 10.1038/s41586-019-0969-x.

(35) Qiu, X.; Mao, Q.; Tang, Y.; Wang, L.; Chawla, R.; Pliner, H. A.; Trapnell, C. Reversed Graph Embedding Resolves Complex Single-Cell Trajectories. Nat. Methods 2017, 14 (10), 979–982. 10.1038/nmeth.4402.

(36) Pape, M.; Miyagi, M.; Ritz, S. A.; Boulicault, M.; Richardson, S. S.; Maney, D. L. Sex Contextualism in Laboratory Research: Enhancing Rigor and Precision in the Study of Sex-Related Variables. Cell 2024, 187 (6), 1316–1326. 10.1016/j.cell.2024.02.008.

(37) Weinreb, C.; Rodriguez-Fraticelli, A.; Camargo, F. D.; Klein, A. M. Lineage Tracing on Transcriptional Landscapes Links State to Fate during Differentiation. Science 2020, 367 (6479). 10.1126/science.aaw3381.

(38) Burkhardt, D. B.; Stanley, J. S.; Tong, A.; Perdigoto, A. L.; Gigante, S. A.; Herold, K. C.; Wolf, G.; Giraldez, A. J.; Dijk, D.; Krishnaswamy, S. Quantifying the Effect of Experimental Perturbations at Single-Cell Resolution. Nature Biotechnology 2021, 1–11. 10.1038/s41587-020-00803-5.

(39) Moon, K. R.; Dijk, D.; Wang, Z.; Gigante, S.; Burkhardt, D. B.; Chen, W. S.; Yim, K.; Elzen, A.; Hirn, M. J.; Coifman, R. R.; Ivanova, N. B.; Wolf, G.; Krishnaswamy, S. Visualizing Structure and Transitions in High-Dimensional Biological Data. Nature Biotechnology 2019, 37 (12), 1482–1492. 10.1038/s41587-019-0336-3.

(40) Buchholtz, L. J.; Ghitani, N.; Lam, R. M.; Licholai, J. A.; Chesler, A. T.; Ryba, N. J. P. Decoding Cellular Mechanisms for Mechanosensory Discrimination. Neuron 2021, 109 (2), 285–298.e5. 10.1016/j.neuron.2020.10.028.

(41) Choi, H. M. T.; Schwarzkopf, M.; Fornace, M. E.; Acharya, A.; Artavanis, G.; Stegmaier, J.; Cunha, A.; Pierce, N. A. Third-Generation in Situ Hybridization Chain Reaction: Multiplexed, Quantitative, Sensitive, Versatile, Robust. Development 2018, 145 (12). 10.1242/dev.165753.

(42) Knoedler, J. R.; Inoue, S.; Bayless, D. W.; Yang, T.; Tantry, A.; Davis, C.; Leung, N. Y.; Parthasarathy, S.; Wang, G.; Alvarado, M.; Rizvi, A. H.; Fenno, L. E.; Ramakrishnan, C.; Deisseroth, K.; Shah, N. M. A Functional Cellular Framework for Sex and Estrous Cycle-Dependent Gene Expression and Behavior. Cell 2022, 185 (4), 654–671.e22. 10.1016/j.cell.2021.12.031.

(43) Xu, X.; Coats, J. K.; Yang, C. F.; Wang, A.; Ahmed, O. M.; Alvarado, M.; Izumi, T.; Shah, N. M. Modular Genetic Control of Sexually Dimorphic Behaviors. Cell 2012, 148 (3), 596–607. 10.1016/j.cell.2011.12.018.

(44) Piekarski, D. J.; Boivin, J. R.; Wilbrecht, L. Ovarian Hormones Organize the Maturation of Inhibitory Neurotransmission in the Frontal Cortex at Puberty Onset in Female Mice. Curr Biol 2017, 27 (12), 1735–1745.e3. 10.1016/j.cub.2017.05.027.

(45) Ishii, K. K.; Hashikawa, K.; Chea, J.; Yin, S.; Fox, R. E.; Kan, S.; Shah, M.; Zhou, C.; Navarrete, J.; Murry, A. D.; Szelenyi, E.; Golden, S.; Stuber, G. Post-Ejaculatory Inhibition of Female Sexual Drive via Heterogeneous Neuronal Ensembles in the Medial Preoptic Area. bioRxiv December 2, 2024, p 2023.09.08.556711. 10.1101/2023.09.08.556711.

(46) Shimogori, T.; Lee, D. A.; Miranda-Angulo, A.; Yang, Y.; Wang, H.; Jiang, L.; Yoshida, A. C.; Kataoka, A.; Mashiko, H.; Avetisyan, M.; Qi, L.; Qian, J.; Blackshaw, S. A Genomic Atlas of Mouse Hypothalamic Development. Nature Neuroscience 2010, 13 (6), 767–775. 10.1038/nn.2545.

(47) Romanov, R. A.; Tretiakov, E. O.; Kastriti, M. E.; Zupancic, M.; Häring, M.; Korchynska, S.; Popadin, K.; Benevento, M.; Rebernik, P.; Lallemend, F.; Nishimori, K.; Clotman, F.; Andrews, W. D.; Parnavelas, J. G.; Farlik, M.; Bock, C.; Adameyko, I.; Hökfelt, T.; Keimpema, E.; Harkany, T. Molecular Design of Hypothalamus Development. Nature 2020, 1–7. 10.1038/s41586-020-2266-0.

(48) Gegenhuber, B.; Tollkuhn, J. Signatures of Sex: Sex Differences in Gene Expression in the Vertebrate Brain. WIREs Developmental Biology 2020, 9 (1), e348. 10.1002/wdev.348.

(49) McCarthy, M. M.; Arnold, A. P.; Ball, G. F.; Blaustein, J. D.; De Vries, G. J. Sex Differences in the Brain: The Not So Inconvenient Truth. Journal of Neuroscience 2012, 32 (7), 2241–2247. 10.1523/JNEUROSCI.5372-11.2012.

(50) Shansky, R. M. Are Hormones a “Female Problem” for Animal Research? Science 2019, 364 (6443), 825–826. 10.1126/science.aaw7570.

(51) Chen, J. Y.; Campos, C. A.; Jarvie, B. C.; Palmiter, R. D. Parabrachial CGRP Neurons Establish and Sustain Aversive Taste Memories. Neuron 2018, 100 (4), 891–899.e5. 10.1016/j.neuron.2018.09.032.

(52) Heymann, G.; Jo, Y. S.; Reichard, K. L.; McFarland, N.; Chavkin, C.; Palmiter, R. D.; Soden, M. E.; Zweifel, L. S. Synergy of Distinct Dopamine Projection Populations in Behavioral Reinforcement. Neuron 2020, 105 (5), 909–920.e5. 10.1016/j.neuron.2019.11.024.

(53) Hashikawa, Y.; Hashikawa, K.; Rossi, M. A.; Basiri, M. L.; Liu, Y.; Johnston, N. L.; Ahmad, O. R.; Stuber, G. D. Transcriptional and Spatial Resolution of Cell Types in the Mammalian Habenula. Neuron 2020, 106 (5), 743–758.e5. 10.1016/j.neuron.2020.03.011.

(54) Mayer, C.; Acosta-Martinez, M.; Dubois, S. L.; Wolfe, A.; Radovick, S.; Boehm, U.; Levine, J. E. Timing and Completion of Puberty in Female Mice Depend on Estrogen Receptor α-Signaling in Kisspeptin Neurons. Proceedings of the National Academy of Sciences 2010, 107 (52), 22693–22698. 10.1073/pnas.1012406108.

(55) Korsunsky, I.; Millard, N.; Fan, J.; Slowikowski, K.; Zhang, F.; Wei, K.; Baglaenko, Y.; Brenner, M.; Loh, P.; Raychaudhuri, S. Fast, Sensitive and Accurate Integration of Single-Cell Data with Harmony. Nat Methods 2019, 16 (12), 1289–1296. 10.1038/s41592-019-0619-0.

(56) Huynh-Thu, V. A.; Irrthum, A.; Wehenkel, L.; Geurts, P. Inferring Regulatory Networks from Expression Data Using Tree-Based Methods. PLOS ONE 2010, 5 (9), e12776. 10.1371/journal.pone.0012776.

(57) Morabito, S.; Miyoshi, E.; Michael, N.; Shahin, S.; Martini, A. C.; Head, E.; Silva, J.; Leavy, K.; Perez-Rosendahl, M.; Swarup, V. Single-Nucleus Chromatin Accessibility and Transcriptomic Characterization of Alzheimer’s Disease. Nat Genet 2021, 1–13. 10.1038/s41588-021-00894-z.

(58) Chen, E. Y.; Tan, C. M.; Kou, Y.; Duan, Q.; Wang, Z.; Meirelles, G. V.; Clark, N. R.; Ma’ayan, A. Enrichr: Interactive and Collaborative HTML5 Gene List Enrichment Analysis Tool. BMC Bioinformatics 2013, 14 (1), 128. 10.1186/1471-2105-14-128.

(59) Kuleshov, M. V.; Jones, M. R.; Rouillard, A. D.; Fernandez, N. F.; Duan, Q.; Wang, Z.; Koplev, S.; Jenkins, S. L.; Jagodnik, K. M.; Lachmann, A.; McDermott, M. G.; Monteiro, C. D.; Gundersen, G. W.; Ma’ayan, A. Enrichr: A Comprehensive Gene Set Enrichment Analysis Web Server 2016 Update. Nucleic Acids Res 2016, 44 (W1), W90–W97. 10.1093/nar/gkw377.

(60) DePasquale, E. A. K.; Schnell, D. J.; Van Camp, P.-J.; Valiente-Alandí, Í.; Blaxall, B. C.; Grimes, H. L.; Singh, H.; Salomonis, N. DoubletDecon: Deconvoluting Doublets from Single-Cell RNA-Sequencing Data. Cell Reports 2019, 29 (6), 1718–1727.e8. 10.1016/j.celrep.2019.09.082.

(61) Butler, A.; Hoffman, P.; Smibert, P.; Papalexi, E.; Satija, R. Integrating Single-Cell Transcriptomic Data across Different Conditions, Technologies, and Species. Nat. Biotechnol. 2018, 36 (5), 411–420. 10.1038/nbt.4096.

(62) Haghverdi, L.; Lun, A. T. L.; Morgan, M. D.; Marioni, J. C. Batch Effects in Single-Cell RNA-Sequencing Data Are Corrected by Matching Mutual Nearest Neighbors. Nature Biotechnology 2018, 36 (5), 421–427. 10.1038/nbt.4091.

(63) Tasic, B.; Yao, Z.; Graybuck, L. T.; Smith, K. A.; Nguyen, T. N.; Bertagnolli, D.; Goldy, J.; Garren, E.; Economo, M. N.; Viswanathan, S.; Penn, O.; Bakken, T.; Menon, V.; Miller, J.; Fong, O.; Hirokawa, K. E.; Lathia, K.; Rimorin, christine; Tieu, M.; Larsen, R.; Casper, T.; Barkan, E.; Kroll, M.; Parry, S.; Shapovalova, N. V.; Hirschstein, D.; Pendergraft, J.; Sullivan, H. A.; Kim, T. K.; Szafer, A.; Dee, N.; Groblewski, P.; Wickersham, ian; cetin, A.; Harris, J. A.; Levi, B. P.; Sunkin, S. M.; Madisen, L.; Daigle, T. L.; Looger, L.; Bernard, A.; Phillips, J.; Lein, E.; Hawrylycz, M.; Svoboda, K.; Jones, A. R.; Koch, christof; Zeng, H. Shared and Distinct Transcriptomic Cell Types across Neocortical Areas. Nature 2018, 563 (7729), 72–78. 10.1038/s41586-018-0654-5.

(64) Zeisel, A.; Hochgerner, H.; Lönnerberg, P.; Johnsson, A.; Memic, F.; Zwan, J.; Häring, M.; Braun, E.; Borm, L. E.; La Manno, G.; Codeluppi, S.; Furlan, A.; Lee, K.; Skene, N.; Harris, K. D.; Hjerling-Leffler, J.; Arenas, E.; Ernfors, P.; Marklund, U.; Linnarsson, S. Molecular Architecture of the Mouse Nervous System. Cell 2018, 174 (4), 999–1014.e22. 10.1016/j.cell.2018.06.021.

(65) Zappia, L.; Oshlack, A. Clustering Trees: A Visualization for Evaluating Clusterings at Multiple Resolutions. GigaScience 2018, 7 (7), giy083. 10.1093/gigascience/giy083.

(66) Davie, K.; Janssens, J.; Koldere, D.; Waegeneer, M. D.; Pech, U.; Kreft, Ł.; Aibar, S.; Makhzami, S.; Christiaens, V.; González-Blas, C. B.; Poovathingal, S.; Hulselmans, G.; Spanier, K. I.; Moerman, T.; Vanspauwen, B.; Geurs, S.; Voet, T.; Lammertyn, J.; Thienpont, B.; Liu, S.; Konstantinides, N.; Fiers, M.; Verstreken, P.; Aerts, S. A Single-Cell Transcriptome Atlas of the Aging Drosophila Brain. Cell 2018, 174 (4), 982–998.e20. 10.1016/j.cell.2018.05.057.

(67) Otis, J. M.; Namboodiri, V. M. K.; Matan, A. M.; Voets, E. S.; Mohorn, E. P.; Kosyk, O.; McHenry, J. A.; Robinson, J. E.; Resendez, S. L.; Rossi, M. A.; Stuber, G. D. Prefrontal Cortex Output Circuits Guide Reward Seeking through Divergent Cue Encoding. Nature 2017, 543 (7643), 103–107. 10.1038/nature21376.

(68) Rossi, M. A.; Basiri, M. L.; McHenry, J. A.; Kosyk, O.; Otis, J. M.; Munkhof, H. E.; Bryois, J.; Hübel, C.; Breen, G.; Guo, W.; Bulik, C. M.; Sullivan, P. F.; Stuber, G. D. Obesity Remodels Activity and Transcriptional State of a Lateral Hypothalamic Brake on Feeding. Science 2019, 364 (6447), 1271–1274. 10.1126/science.aax1184.

